# Codebook: sequence specificity and genomic binding of poorly-characterized human transcription factors

**DOI:** 10.1101/2024.11.11.622097

**Authors:** Arttu Jolma, Kaitlin U. Laverty, Ali Fathi, Ally W.H. Yang, Isaac Yellan, Ilya E. Vorontsov, Sachi Inukai, Judith F. Kribelbauer-Swietek, Antoni J. Gralak, Rozita Razavi, Mihai Albu, Alexander Brechalov, Zain M. Patel, Vladimir Nozdrin, Georgy Meshcheryakov, Andrey Buyan, Ivan Kozin, Sergey Abramov, Alexandr Boytsov, The Codebook Consortium, Matthew T. Weirauch, Oriol Fornes, Vsevolod J. Makeev, Jan Grau, Ivo Grosse, Philipp Bucher, Bart Deplancke, Ivan V. Kulakovskiy, Timothy R. Hughes, The Codebook consortium, Philipp Bucher, Bart Deplancke, Oriol Fornes, Jan Grau, Ivo Grosse, Timothy R. Hughes, Arttu Jolma, Fedor A. Kolpakov, Ivan V. Kulakovskiy, Vsevolod J. Makeev, Mihai Albu, Marjan Barazandeh, Alexander Brechalov, Zhenfeng Deng, Ali Fathi, Chun Hu, Samuel A. Lambert, Kaitlin U. Laverty, Zain M. Patel, Sara E. Pour, Rozita Razavi, Mikhail Salnikov, Ally W.H. Yang, Isaac Yellan, Hong Zheng, Georgy Meshcheryakov, Giovanna Ambrosini, Antoni J. Gralak, Sachi Inukai, Judith F. Kribelbauer-Swietek, Marie-Luise Plescher, Semyon Kolmykov, Ivan Yevshin, Nikita Gryzunov, Ivan Kozin, Mikhail Nikonov, Vladimir Nozdrin, Arsenii Zinkevich, Katerina Faltejskova, Pavel Kravchenko, Sergey Abramov, Alexandr Boytsov, Vasilii Kamenets, Dmitry Penzar, Anton Vlasov, Ilya E. Vorontsov, Aldo Hernandez-Corchado, Hamed S. Najafabadi, Quaid Morris, Xiaoting Chen, Matthew T. Weirauch

**Affiliations:** Donnelly Centre, University of Toronto, Toronto, ON M5S 3E1, Canada; Sloan Kettering Institute, Memorial Sloan Kettering Cancer Center, New York, NY 10065, USA; Department of Molecular Genetics, University of Toronto, Toronto, ON M5S 1A8, Canada; Vavilov Institute of General Genetics, Russian Academy of Sciences, 119991, Moscow, Russia; Laboratory of Systems Biology and Genetics, Institute of Bioengineering, School of Life Sciences, École Polytechnique Fédérale de Lausanne, 1015, Lausanne, Switzerland; Swiss Institute of Bioinformatics, 1015, Lausanne, Switzerland; Faculty of Bioengineering and Bioinformatics, Lomonosov Moscow State University, 119991, Moscow, Russia; Institute of Protein Research, Russian Academy of Sciences, 142290, Pushchino, Russia; Altius Institute for Biomedical Sciences, Seattle, WA 98121, USA; Center for Autoimmune Genomics and Etiology, Cincinnati Children’s Hospital Medical Center, Cincinnati, OH, USA; Divisions of Allergy and Immunology, Biomedical Informatics, Human Genetics, and Developmental Biology, Cincinnati Children’s Hospital Medical Center, Cincinnati, OH, USA; Department of Pediatrics, University of Cincinnati College of Medicine, Cincinnati, OH, USA; Department of Medical Genetics, Centre for Molecular Medicine and Therapeutics, BC Children’s Hospital Research Institute, University of British Columbia, Vancouver, BC V5Z 4H4, Canada; Institute of Computer Science, Martin Luther University Halle-Wittenberg, 06099, Halle, Germany; Institute of Biochemistry and Genetics, Ufa Federal Research Centre of Russian Academy of Sciences, 450054, Ufa, Russia; Department of Computational Biology, Sirius University of Science and Technology, 354340, Sirius, Krasnodar region, Russia; Bioinformatics Laboratory, Federal Research Center for Information and Computational Technologies, 630090, Novosibirsk, Russia; Bioinformatics Competence Center, Université de Lausanne, 1015 Lausanne, Switzerland; Chugai Pharmaceutical Co., Ltd, Tokyo, 103-8324, Japan; Biosoft.Ru LLC, 630058, Novosibirsk, Russia; Institute of Organic Chemistry and Biochemistry of the Czech Academy of Sciences, 160 00 Praha 6, Czech Republic; Computer Science Institute, Faculty of Mathematics and Physics, Charles University, 118 00 Praha 1, Czech Republic; Max Planck Institute of Biochemistry, 82152, Planegg, Germany; TUD Dresden University of Technology, Center for Molecular and Cellular Bioengineering (CMCB), Biotechnologisches Zentrum (BIOTEC), Tatzberg 47/49, 01307 Dresden, Germany; Department of Human Genetics, McGill University, Montréal, QC H3A 0C7, Canada; Victor P. Dahdaleh Institute of Genomic Medicine, Montréal, QC H3A 0G1, Canada

**Keywords:** Transcription factor, TF, ChIP-seq, HT-SELEX, GHT-SELEX, SELEX, SMiLE-seq, Motif, DNA-binding specificity, PWM, PBM, Codebook

## Abstract

Gene expression is regulated by transcription factors (TFs), which recognize specific DNA sequence motifs. Several hundred putative human TFs, identified mainly by an apparent DNA-binding domain, lack known binding motifs^1^, and even for well-characterized TFs, it remains controversial to what degree motifs accurately reflect binding sites in living cells^2,3^. Here, we describe a systematic effort (“Codebook”) to determine the sequence specificity of 332 putative and poorly characterized human TFs. Over 4,000 independent experiments, encompassing multiple *in vitro* and *in vivo* assays, produced motifs for just over half (177, or 53%), of which most are unique to a single protein, thereby extending the vocabulary of sequence recognition encoded by human TFs by ∼100 distinct motifs. Moreover, binding motifs identified *in vitro* are strongly enriched within cellular binding sites. Collectively, the data reveal tens of thousands of previously unknown, conserved, and direct TF binding sites across the human genome. These sites are concentrated in promoter regions, and are predictive of gene expression, illustrating that this new data atlas provides an important step forward in decoding the human genome.

## Introduction and motivations

A 2018 survey of putative human transcription factors (TFs) concluded that over a quarter of the estimated ∼1,600 possible TFs lacked established binding motifs^1^. This is a striking deficit given that most of the conserved DNA in the human genome is noncoding^4^, and that a primary hypothesis for the function of noncoding DNA is gene regulation^5^. Moreover, most of the uncharacterized and putative TFs are not close paralogs of any established TF. Thus, despite possessing either literature evidence of DNA-binding or protein domains that would typically bind DNA, their binding motifs cannot be readily inferred, although it also cannot be excluded that these putative TFs lack DNA sequence specificity entirely.

TF DNA-binding motifs are commonly modelled as Position Weight Matrices (PWMs), which describe the relative preference of a TF for each nucleotide base pair in the binding site^6,7^, and can be visualized as a sequence logo^8^. Different methods for measuring TF binding, and for deriving PWMs from the resultant data, have different inherent limitations and biases^6^. There is also long-standing controversy regarding the contribution of a TF’s inherent sequence specificity to its cellular binding, relative to the influence of chromatin and cofactors^2^. These uncertainties represent fundamental hurdles for the analysis of gene regulation, as well as a myriad of related tasks in genome analysis, including the interpretation of conserved genomic elements and sequence variants.

To address these issues, we analyzed a large majority of the poorly characterized human TFs^1^ (defined here as having no confidently known binding motif), as well as several dozen previously studied control TFs^9,10^, using a panel of assays that provide different perspectives on DNA sequence specificity. We refer to this international collaborative project as the “Codebook/GRECO-BIT Collaboration”: the reagent set and laboratory experiments were initiated as the “Codebook Project”, alluding to the fact that TFs decode individual “words” in the genome, and the <u>G</u>ene <u>RE</u>gulation <u>CO</u>nsortium <u>B</u>enchmarking Ini<u>T</u>iative, GRECO-BIT rooted in GRECO^11^, was then engaged for much of the data analysis and benchmarking.

In this paper, we present an overview of the Codebook data, major outcomes and findings of the study, and examples of prevalent phenomena and applications. These include:

- Just over half of the 332 putative (i.e., “Codebook”) TFs (177) display DNA sequence specificity, most with the same motifs observed *in vitro* and in living cells.
- The resulting 177 motifs are largely different from one another, and from the motifs of previously-studied human TFs.
- Tens of thousands of genomic binding sites for these TFs display evidence of purifying selection at the motif matches, indicating that they are functional.
- These conserved binding sites are abundant in the promoters of coding genes, and promoter binding sites of the Codebook TFs are predictive of gene expression across tissues and cell types.
- Many TFs bind to genomic dark matter, such as transposons, or are derived from transposons, suggesting they may have had roles in adaptive evolution.

We also introduce web resources that can be used to access the primary and processed data, including the PWMs. Accompanying manuscripts and SI provide greater technical detail and depth, including biological findings, new assays, and intriguing TF families, as well as methods for identifying and benchmarking PWMs^12–15^.

### Overview of the Codebook project

**Fig. 1** provides a schematic of the Codebook project. We analyzed 393 proteins (332 putative “Codebook” TFs, and 61 control TFs) (**Supplementary Table 1**) using up to five different assays encompassing both *in vitro* and living cellular contexts: HT-SELEX, SMiLE-seq, Protein Binding Microarrays (PBMs), ChIP-seq (with inducible tagged transgenes in HEK293 cells), and GHT-SELEX, a novel variant of HT-SELEX in which the selections are performed using fragmented genomic DNA^12^, instead of random sequences. The GHT-SELEX, SMiLE-seq, and ChIP-seq data yielded significant and extensive biological insight on their own that is detailed in accompanying manuscripts^12,13,15^, with highlights and joint conclusions summarized here. Different assays accommodate different protein tags and expression systems, and many TFs were analyzed using multiple constructs (e.g., full-length and DNA-binding domain(DBD)(s)) and multiple expression systems, leading to variable numbers of constructs analyzed and experiments per protein. The study employed 622 and 93 protein-coding inserts cloned into 1267 and 118 protein expression constructs for the Codebook TFs and controls, respectively (**Supplementary Table 2** and **Supplementary Table 3**). In total, we performed 4,804 experiments, 4,377 for Codebook proteins and 427 for control TFs (**Supplementary Table 4**).

**Figure 1.**
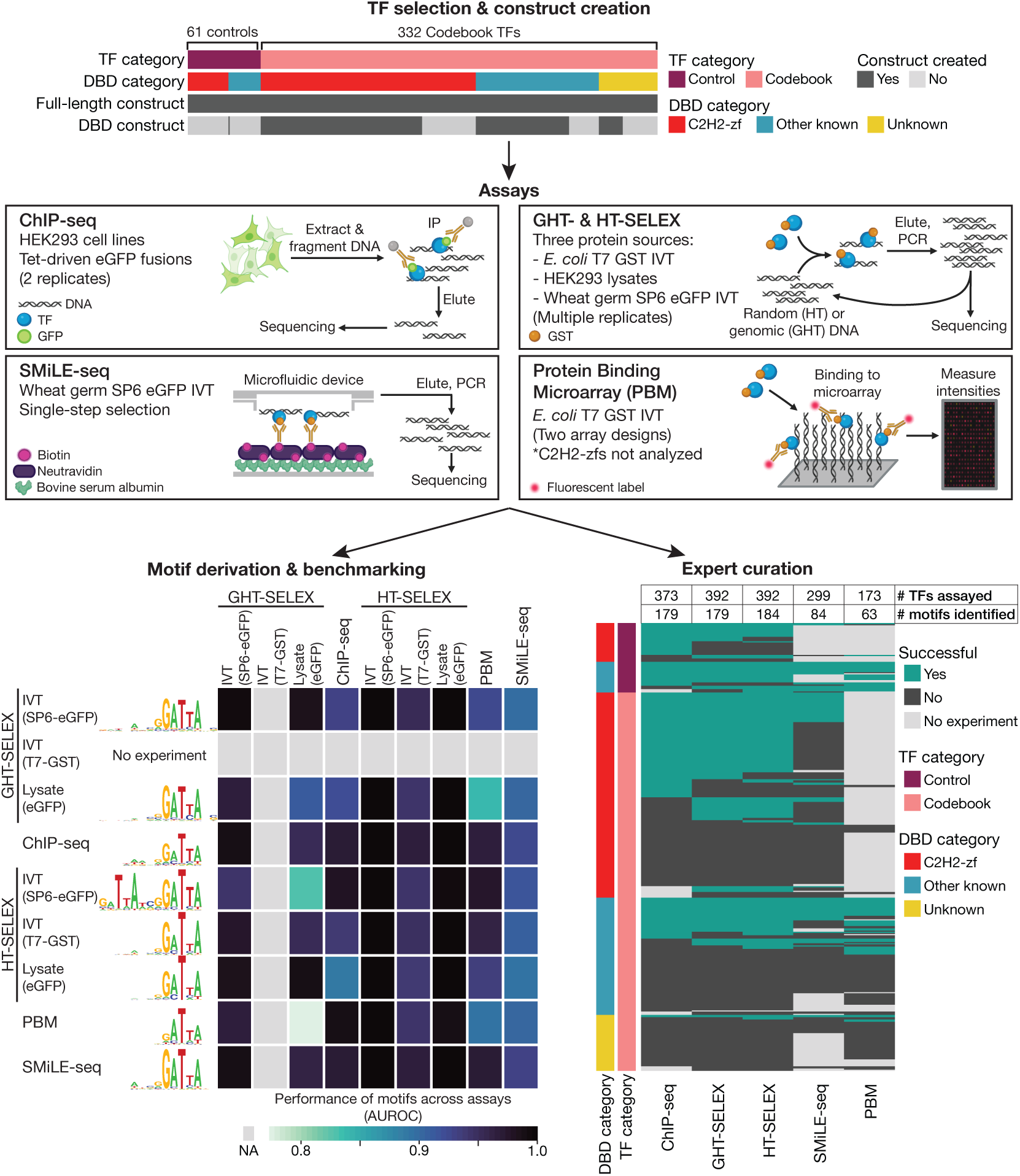
Codebook project overview. ***Top***, Categories of 393 TFs assayed and their associated constructs. ***Middle***, Graphical summary of assays employed. ***Bottom left***, Example of performance (as AUROC) of the best performing PWM for TPRX1, for each combination of experiment type – one for PWM derivation (rows), and one for PWM testing (columns). ***Bottom right***, Heatmap shows successful experiments for all 393 TFs across all experiment types. AUROC: area under precision-recall curve, PWM: position weight matrix, TF: transcription factor, C2H2-zfs: C2H2 zinc finger proteins, IP: immunoprecipitation, DBD: DNA-binding domain, IVT: in vitro transcription/translation

Because the Codebook proteins had no known motif, the process of determining which experiments were successful was, by necessity, coupled to the process of deriving motifs from each of the data sets, as described in detail in a separate manuscript^14^.

Briefly, if we obtained similar motifs for the same protein from two or more different experimental platforms, and these motifs were predictive of data across platforms, then we considered the experiments producing these motifs to be successful. In total, 1,073 experiments for Codebook proteins and 280 for controls were deemed successful (i.e., produced a motif that was reproducible across experiments) (**Supplementary Table 4**). In total, we obtained motifs for 177/332 Codebook proteins and 58/61 control TFs using the same process, suggesting a low overall false-negative rate. The number of Codebook proteins successful in a particular assay was 130 for ChIP-seq (41% of those tested), 144 for HT-SELEX (44% of those tested), 139 for GHT-SELEX (42% of those tested), 64 for SMiLE-seq (23% of those tested), and 39 for PBMs (27% of those tested) (**Extended Data Fig. 1a**,**b**, **Supplementary Table 1**). For each of the proteins, we selected a single representative PWM that scored highly across data types, for use in subsequent analyses (see **Methods**, **Supplementary Table 5**).

The 177 successful Codebook TFs are dominated by the large and diverse C2H2 zinc finger (C2H2-zf) class, for which 67% (121/180) produced motifs. C2H2-zf proteins can bind RNA, protein, or other ligands^16–18^; this outcome indicates that most of the Codebook C2H2-zf proteins are DNA-binding proteins, although it does not exclude other functions. Among the Codebook TFs with other DBD types, roughly half (49%, 50/103) also produced motifs. The Codebook TFs also encompassed 49 proteins that lack a well-established DBD, which were included due to literature evidence for DNA sequence specificity. Among these, only six (12%) yielded motifs. Upon closer inspection, all six do, in fact, contain plausible DNA-binding regions or domains (see **Supplementary Discussion**). For example, DACH1 and DACH2, which recognize the same motif, are homologs of Drosophila *Dachshund*, sharing the SKI/SNO/DAC domain. There is prior evidence of DNA-binding by SKI/SNO/DAC domains^19,20^, although it is also known to have other functions^21^. Subsequent to the main Codebook study, we tested all five additional human SKI/SNO/DAC domain-containing proteins with HT-SELEX, and identified a motif for one, SKIDA1, providing further support for SKI/SNO/DAC acting as a DNA-binding domain (**Extended Data Fig. 2a-c** and **Supplementary Discussion**).

Nearly half of the Codebook proteins did not yield motifs (PWMs) that passed our success criteria. False negatives could arise from the lack of an obligate binding partner, a requirement for epigenetically modified DNA, lack of requisite post-translational modification in our experiments, or other limitations of the methods. We manually re-assessed each of the failed proteins, and deemed that the majority (83/155) likely represent true negatives. C2H2-zf proteins with a single C2H2-zf domain, or composed only of unusual C2H2-zf domains, typically failed in our assays, as did those that lack the canonical spacing of C2H2-zf domains that supports binding of tandem C2H2-zf arrays to longer sites^22^ (see **Supplementary Discussion**, **Extended Data Fig. 3**, **Supplementary Table 6**, and **Supplementary Document 1** for discussion of the unsuccessful proteins), suggesting that these C2H2-zf domains may have roles other than DNA-binding. Some subtypes of other bona fide DBD classes also lack sequence specificity (e.g., HMG); we found almost two dozen such cases. In addition, the 155 putative TFs that did not yield a motif in any assay are enriched for those with no known DBD (of which 43/49 failed); among these, 33 have no evidence for DNA binding beyond a single study, most of which were published over two decades ago.

### Motifs identified *in vitro* are highly enriched at cellular binding sites and predict differential binding to SNVs

The Codebook project generated technically successful ChIP-seq data for 130 of the 177 successful Codebook TFs^13^, allowing us to gauge utilization of the motif in a cellular context. There is clear enrichment of motif matches within ChIP-seq peaks, indicating that the intrinsic sequence specificities of the TFs are used in the selection of cellular binding sites (**Fig. 2**). Some TFs display motif enrichment in a wide region around the peak summit due to multiple repeated motif matches; this phenomenon is particularly common at CpG island promoters (discussed below). An accompanying manuscript^12^ demonstrates that genomic binding *in vitro* and *in vivo* are more similar than generally believed. This conclusion could only be reached by a combination of GHT-SELEX and ChIP-seq data coupled with optimization of PWM derivation and scanning methods.

**Figure 2.**
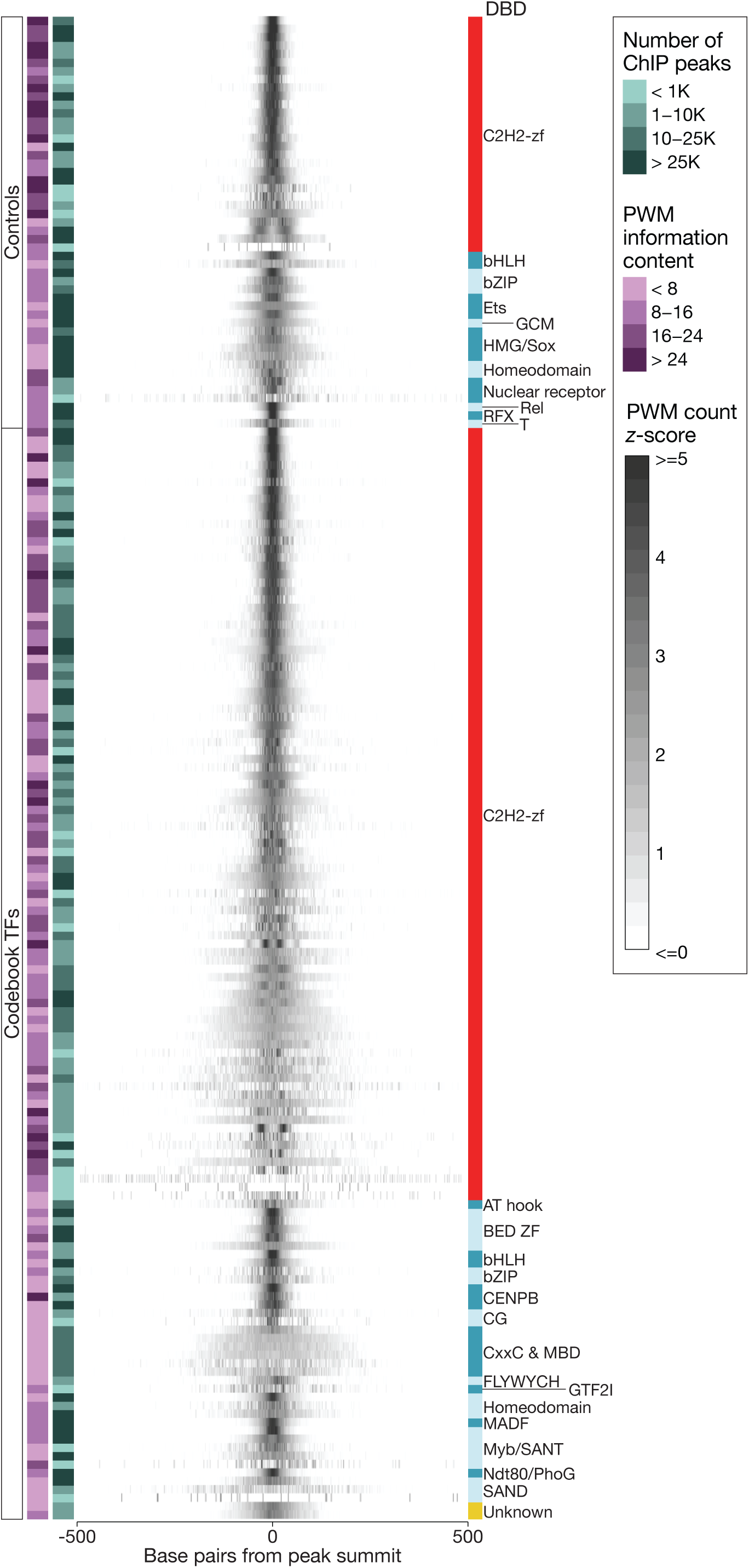
Motif enrichment at ChIP-seq binding sites. Heatmap displays the standardized count of representative PWM matches at each base +/-500 bp from the TF’s ChIP-seq peak summits. PWM matches were identified with MOODS (P-value < 0.001) and aggregated for each TF at each base position. For visualization, counts were standardized (z-score across all base positions) for each TF. PWM information content and the number of ChIP-seq peaks for each TF are indicated on the left. Control and Codebook TFs are separated and grouped by DBD type. PWM: position weight matrix, DBD: DNA-binding domain, ChIP: chromatin immunoprecipitation, TF: transcription factor

The Codebook project was conducted over a period of nearly six years, and during this time, several large-scale studies aimed at systematic ChIP-seq analysis of human TFs (e.g., ENCODE) were published, mainly employing cell lines other than HEK293^23–25^. Peaks from these external datasets overlap only partially with the Codebook ChIP-seq peaks for the same TF, which is expected because chromatin and cofactors can impact site accessibility and selection (**Extended Data Fig. 4a-d**, **Supplementary Table 7**). Nonetheless, Codebook PWMs were generally able to discriminate between binding sites in these other ChIP-seq experiments and random sequences (median AUROC of 0.71 on HEK293, and slightly lower on other cell types), illustrating that the motifs are predictive across studies and cell types (**Extended Data Fig. 4e**).

Sequence variants can disrupt TF binding sites, impacting gene regulation^26^. We examined allelic read imbalance^27^ in ChIP-seq and GHT-SELEX peak sets (**Extended Data Fig. 5**, **Supplementary Table 8**), detecting a total of 12,060 apparent allele-specific binding (ASB) events at 10,009 common single-nucleotide polymorphisms (SNPs) for 190 individual TFs at 5% FDR. Importantly, for highly motif-disrupting variants (with at least two-fold difference in P-values between PWM predictions for the reference and the alternative alleles), the allelic preference of ASBs was concordant with PWM predictions for 1,682 out of 2,260 (74%) of ASBs. Thus, while the specific ASBs detected experimentally are limited to SNPs present in HEK293 cells, the Codebook motifs can be applied to predict differential TF binding to any other sequence variants. Considering aggregated ASBs across all tested TFs and experiments (see Methods), the corresponding SNPs were enriched for previously characterized ASBs^28^ (2.4 odds-ratio, P-value 3.90e-162), disease-associated SNPs^29^ (1.35 odds-ratio, P-value 3.79e-18,), and GTEx eQTLs^30^ (1.34 odds-ratio, P-value 1.77e-41) (two-sided Fisher’s exact test) (**Supplementary Table 9**).

### Most of the Codebook TF motifs are unique, and many require degeneracy for highest motif performance

The Codebook protein sequences are typically not closely related to each other or to those of known TFs, suggesting that their DNA sequence preferences may also differ^1^. **Fig. 3** shows an overview of similarity^31^ among the representative Codebook PWMs, illustrating that most of the Codebook TF motifs are indeed unique. The majority are also dissimilar to any other known PWM^14^ (**Extended Data Fig. 6**). This result is partly explained by the large number of C2H2-zf proteins, which differ in their DNA-contacting “specificity residues”^32^. Small clusters along the diagonal in **Fig. 3** mostly correspond to the handful of paralogs analyzed (e.g., TIGD4 and 5, SP140 and SP140L, DACH1 and 2, CAMTA1 and 2, and ZXDA, B, and C). In addition, the middle of **Fig. 3** contains a set of eight TFs for which the motif is dominated by a single CG sequence. Five of them (CXXC4, FBXL1, TET3, MBD1, and KDM2A) contain CXXC-zf domains, which are expected to bind clusters of unmethylated CpGs^33^. Similarly, in the lower right is a group of five AT-hook proteins that prefer A/T containing sequences. Overall, however, the 177 Codebook TFs encompass 129 distinct motifs (**Fig. 3**, **Methods**). When we performed motif clustering jointly with previously-catalogued, representative motifs for 1,211 human TFs, we obtained only 4.75-fold more distinct motif clusters (613), of which 92 (15%) contained only Codebook proteins (**Extended Data Fig. 6**, **Supplementary Table 10**). These numbers are robust to methodology: utilization of a different public motif database, a different similarity metric and a different clustering procedure led to a similar outcome, with a total of 653 motif clusters, 135 (21%) of which contained only Codebook motifs (**Supplementary Table 11, Methods**). Thus, the Codebook data expands the lexicon of human TFs by at least 15%, providing ∼100 previously unknown motifs.

**Figure 3.**
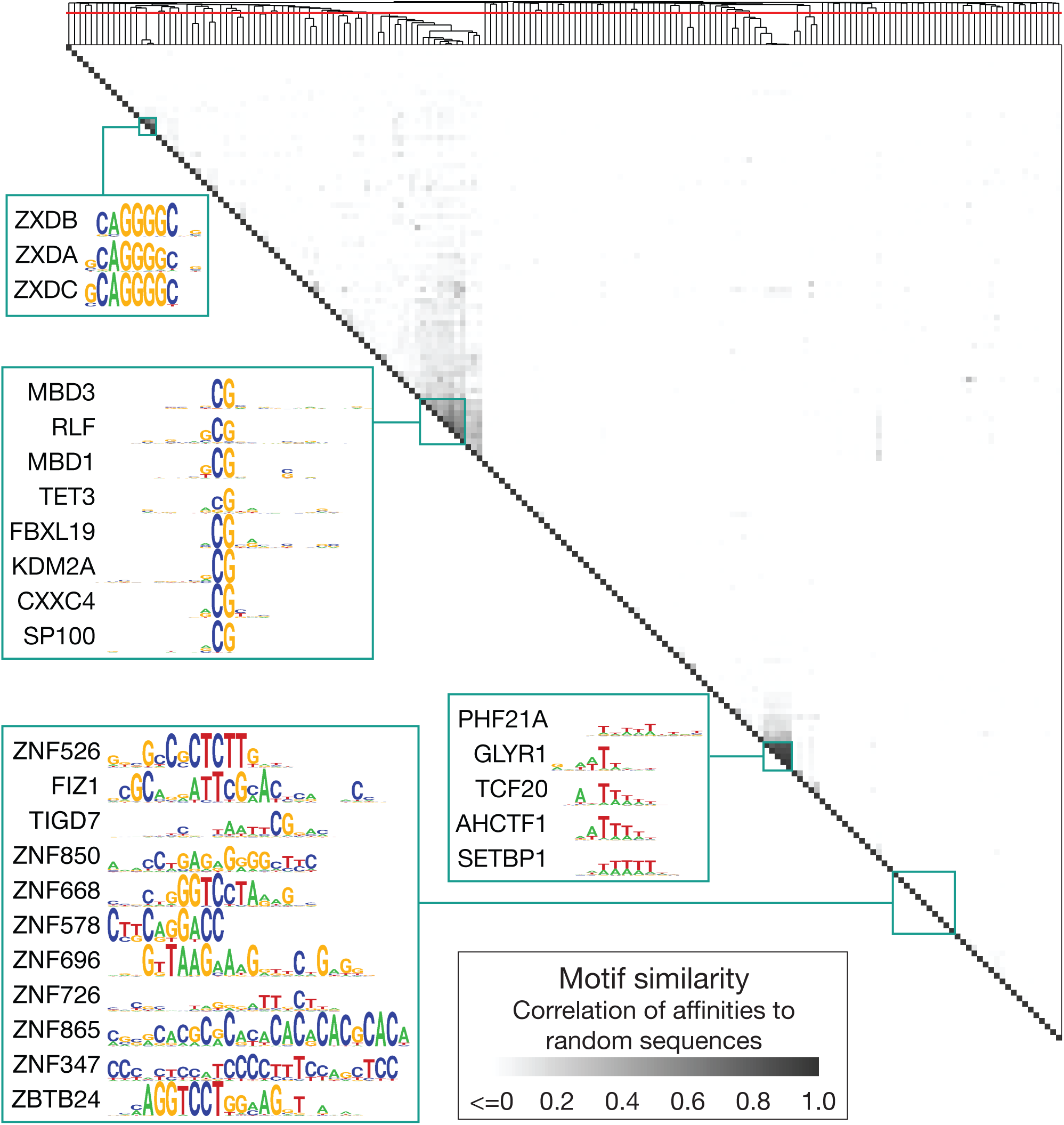
Similarity of Codebook TF motifs. Symmetric heatmap displays the similarity between representative PWMs for Codebook TFs, clustered by Pearson correlation with average linkage. PWM similarity is the correlation between pairwise affinities to 150,000 random sequences of length 100, as calculated by MoSBAT^31^. The red line across the dendrogram indicates the clustering threshold determined by the optimal silhouette value^84^. Pullouts and labels illustrate specific points in the main text. TF: transcription factor, PWM: position weight matrix

For dozens of TFs, the representative PWM, selected for highest accuracy across data sets, had a degenerate quality, i.e. there were few or no positions at which a specific base is absolutely required (**Extended Data Fig. 7a**). Artificially increasing the information content (IC) (i.e., “unflattening” the sequence logo, and thus increasing the apparent “specificity” of the low-IC representative PWMs) almost universally reduced motif performance (**Extended Data Fig. 7b,c**), indicating that the degeneracy is required for accuracy across data types. We also found that, overall, IC is not predictive of motif performance^14^. It is counterintuitive that degeneracy (i.e., lower apparent specificity) would lead to better predictive capacity, but similar findings by others support the validity of the result^34–36^. The degeneracy may be a consequence of forcing a single PWM to represent the specificity of TFs that, in reality, recognize multiple related motifs^14^. It is also conceivable that TFs having the ability to tolerate mutations in binding sites is beneficial for evolution^37^.

### Codebook TF binding sites suggest functions for tens of thousands of conserved elements, which often co-occur in promoters

Together, the Codebook assays and PWMs pinpoint genomic loci that are bound directly by each TF *in vivo* (i.e., in ChIP-seq), by identifying sites that are also bound *in vitro* (i.e., GHT-SELEX), and that contain a PWM match, thus allowing base-level resolution. We refer to these as “triple <u>o</u>verla<u>p</u>” (TOP) sites, which are taken as the overlap of the three sets (ChIP-seq, GHT-SELEX, and PWM matches) after applying optimized score thresholds for each (see **Methods** for details). To further gauge functionality of the TOP sites, we examined whether the pattern of per-nucleotide conservation^38^ at each site is consistent with the TF’s sequence preference driving local sequence constraint. To do this, we tested whether the motif matches at TOPs are more highly conserved than neighboring sequence, and whether the degree of conservation at each position within the motif match correlates with the contribution of that base to predicted relative affinity (see **Methods** for details). **Fig. 4a** shows several examples illustrating apparent conservation of PWM matches, for both control and Codebook TFs. Overall, 85 of the 101 Codebook TFs (as well as 31/36 controls) with successful GHT-SELEX and ChIP-seq data displayed conservation of at least one TOP site (FDR < 0.1). In total we identified 113,577 such conserved TOP sites (“CTOP” sites) (82,760 for Codebook TFs and 30,817 for controls), encompassing 1,394,530 bases within the human genome. These results provide strong support for the functional importance of Codebook TF binding sites in the genome.

**Figure 4.**
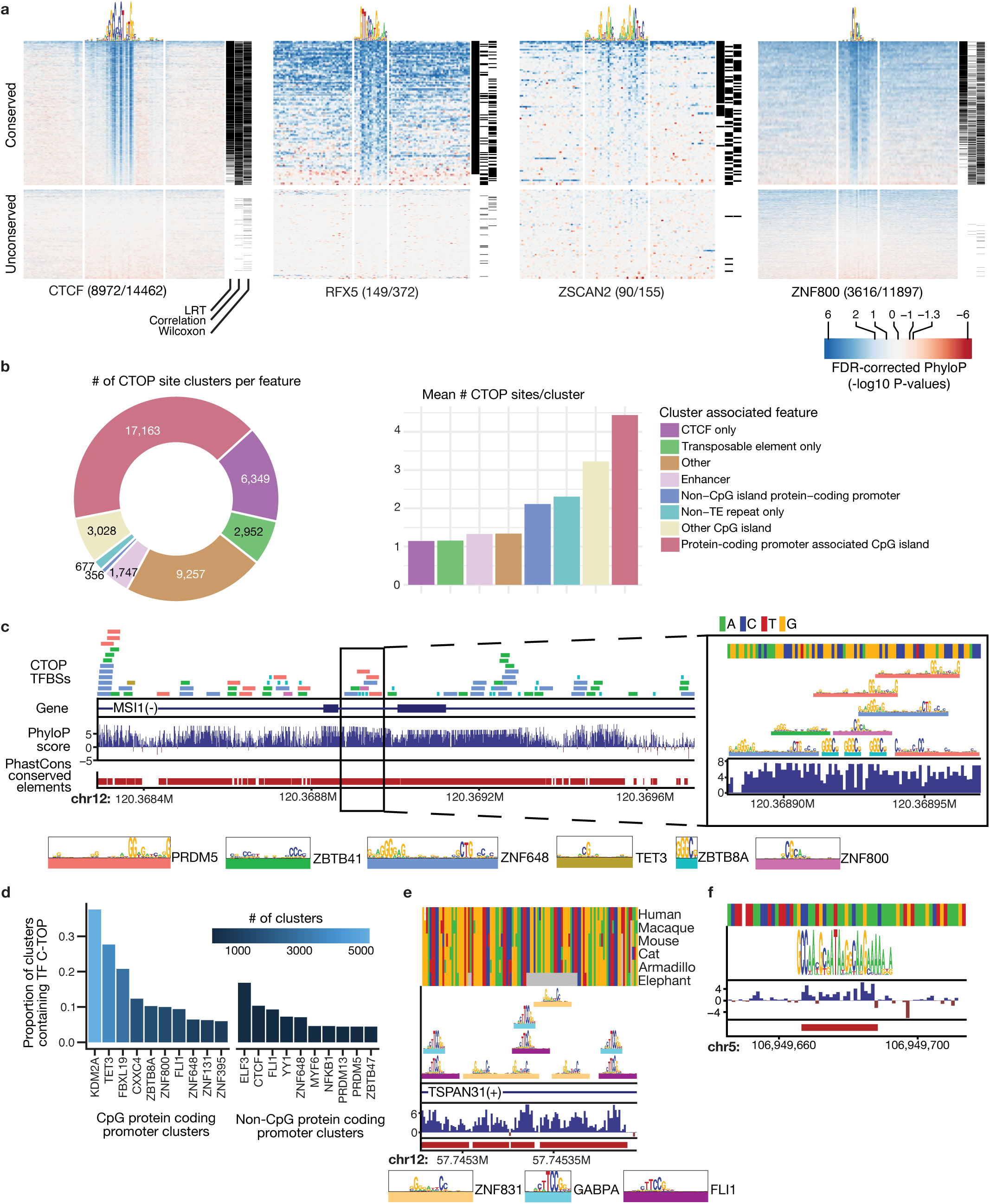
Conservation of Codebook TF binding sites and association with genomic features. a, Heatmaps of FDR-corrected phyloP scores over the PWM match and 50 bp flanking for TOP sites for four TFs (two controls and two Codebook TFs). Statistical test results (see main text and **Methods**) are indicated at right. **b,** *Left,* Donut plot displays the proportion and number of clusters of conserved TOP (CTOP) sites that overlap the genomic features indicated. *Middle,* Bar plot displays the mean # of individual CTOPs contained within clusters that overlap the examined genomic regions. **c,** A 1,420-base, CpG-island-overlapping CTOP cluster (chr12:120368293-120369713). Zoonomia 241-mammal phyloP scores and Multiz 471 Mammal alignment PhastCons Conserved Elements are shown. **d,** Bar plot of the frequency of TFs with CTOPs that occur most frequently in CTOP clusters that overlap CpG and non-CpG protein coding promoters, respectively. **e,** CTOP cluster overlapping the non-CpG promoter at chr12:57,745,278-57,745,396. **f,** CTOP site for the KRAB-C2H2-zf protein ZNF689, overlapping an L1M4-type L1 element located at chr5:106949666-106949692. TF: transcription factor, TOP: triple-overlap, CTOP: conserved triple-overlap

Many of the CTOP sites were either overlapping or adjacent to other CTOP sites for the same or different TFs, suggesting that these sites may be part of the same regulatory element. We therefore grouped the CTOPs that were within 100bp from each other, yielding 41,529 CTOP clusters (of which 24,733 are singletons and 16,796 contain multiple CTOPs). The majority (35,583, or 85.7%) overlapped with the UCSC PhastCons track. Thus, binding of these TFs presents a likely biochemical function for these previously-identified conserved elements. The remaining 5,946 CTOPs appear to represent new functional annotations that were not readily detected by comparative genomics alone. It is known that TF binding motifs can provide additional functional information (e.g., degenerate positions within motif matches) that can explain complex conservation patterns^39^.

Codebook TFs with the largest number of CTOP sites were typically located within promoters (36.9% of all CTOP clusters) and/or CpG islands (47.3% of CTOP clusters) (**Fig. 4b**). The majority of CpG islands that are within protein-coding genes (60.4%, 8,111/13,427) contained CTOP sites, with an average of 4.55 CTOP sites per CpG island. Moreover, 56/101 of Codebook TFs analyzed had at least one CTOP site within a CpG island. An example CTOP that overlaps a CpG island is shown in **Fig. 4c**. The extent of specific, conserved, and intrinsic occupancy of CpG islands by many TFs of diverse classes is surprising: CpG islands are known to be enriched for binding sites for specific TFs (SP1, NRF1, E2F, and ETS classes), which we do observe (**Fig. 4c-e**), but CpG islands typically have little long-range conservation^40^. The CXXC proteins are known to specifically recognize unmethylated CG dinucleotides and to modulate chromatin at promoters^41^, and we do observe that the CXXC proteins KDM2A, CXXC4, FBXL19, and TET3 predominantly bind CpG islands. The abundance of CG dinucleotides in CpG islands has been attributed primarily to their lack of methylation in the germline, however, rather than primary sequence constraint or binding to TFs^42^. The Codebook data thus provide a new lens for study of CpG islands. Intriguingly, many of the Codebook TFs with CTOP sites in CpG islands recognize elaborate C/G rich motifs, often with degenerate positions and long gaps between invariant bases (**Fig. 4c**).

CTOP clusters were also found in non-CpG island protein-coding promoters (**Fig. 4b**) (678/6,606 such promoters, defined as -1000 to +500 relative to TSS). These clusters are not dominated by any specific TFs, although some TFs are more prevalent than others (e.g. controls ELF3 and CTCF, and Codebook TF ZNF648) (**Fig. 4d**). **Fig. 4e** shows an example of a CTOP cluster in a non-CpG promoter, occurring early in the first intron of the *TSPAN31* gene, which exhibits apparent conserved spacing and orientation of multiple Codebook TF binding sites. In contrast, CTOP clusters outside of promoters and CpG islands often contain just one or two CTOP sites (**Fig. 4b)**. One example is a very strongly conserved ZNF689 binding site found in a lncRNA intron and overlapping an L1M4 transposon (**Fig. 4f**).

CTOP clusters also overlapped with known and putative enhancers: 1,825 overlap with HEK293 enhancers (defined by ChromHMM^13^), and 4,057 are found in the extensive GeneHancer annotation set^43^. These lower numbers, relative to promoters, could be attributed to the relatively rapid evolution of enhancers^44^, lack of complete knowledge of enhancer identities, depletion of enhancer-related functions among the Codebook TFs, or to other, non-enhancer functions of these conserved sites, such as control of heterochromatin or other aspects of chromosome architecture.

### Codebook TF motifs predict gene expression levels across tissues and cell types

Conservation of TF binding sites is a strong indicator that they are functional elements, but it does not inherently reveal their biological purpose, i.e. in what conditions they are active, and what processes they regulate. To link Codebook motifs to biological functions, we employed Motif Activity Response Analysis (MARA)^45,46^, a framework that identifies relationships between the promoter motif occurrences and gene expression in different biological samples. For MARA, we chose a representative motif from each of the 653 motif clusters (**Supplementary Table 11**), scanned the FANTOM5 promoters^47^ with the representative motifs, and used the resulting scores to predict the promoter activity (normalized CAGE tag counts) in 583 biological samples (tissues, primary cells, and cell lines) using a linear model (**See Methods, Extended Data Fig. 8a, Supplementary Table 12**). Next, for each motif cluster, we calculated the Pearson correlation between the predicted motif activity and the total expression of the corresponding TFs across all samples. The distribution of resulting correlation coefficients for 135 motif clusters containing only Codebook motifs is similar to that of motif clusters for previously known TFs (**Extended Data Fig. 8b**), supporting the notion that the Codebook proteins are bona fide regulatory factors.

To explore the cellular functions the Codebook TFs might regulate, we examined the 25 Codebook-only motif clusters with the highest variance in activity across the FANTOM5 samples. Bi-clustering activities yielded four major TF groups (**Fig. 5**): (**1**) TFs targeting genes that are preferentially expressed in immune cells, including transcriptional repressors ZBTB8A^48^ and ZFTA, whose targets are specifically repressed in the brain, and are often activated in ependymomas caused by ZFTA fusion with RELA^49^; (**2**) TFs active in both immune cells and cancer, including proteins that bind unmethylated CG-containing motifs (CGGBP1, CXXC4, and SP140), the transcriptional activator ZNF131^50^, and MBD1; (**3**) TFs with target genes specifically expressed in nervous tissues, including neuron maturation regulators CAMTA1^51^ and ZNF292^52^, neurogenesis-associated repressors MBD3^53^ and DNTTIP1^54^, as well as MYRFL, the paralog of myelin regulatory factor (MYRF), critical for central nervous system myelination^55^, exclusively active in the adult nervous tissues; (**4**) TFs mostly restricted to mesodermal primary cells, including rRNA transcription regulator TTF1^56^ and zinc finger and BTB domain-containing proteins ZBTB41 and ZBTB5, which were found to stimulate cell proliferation^57,58^.

**Figure 5.**
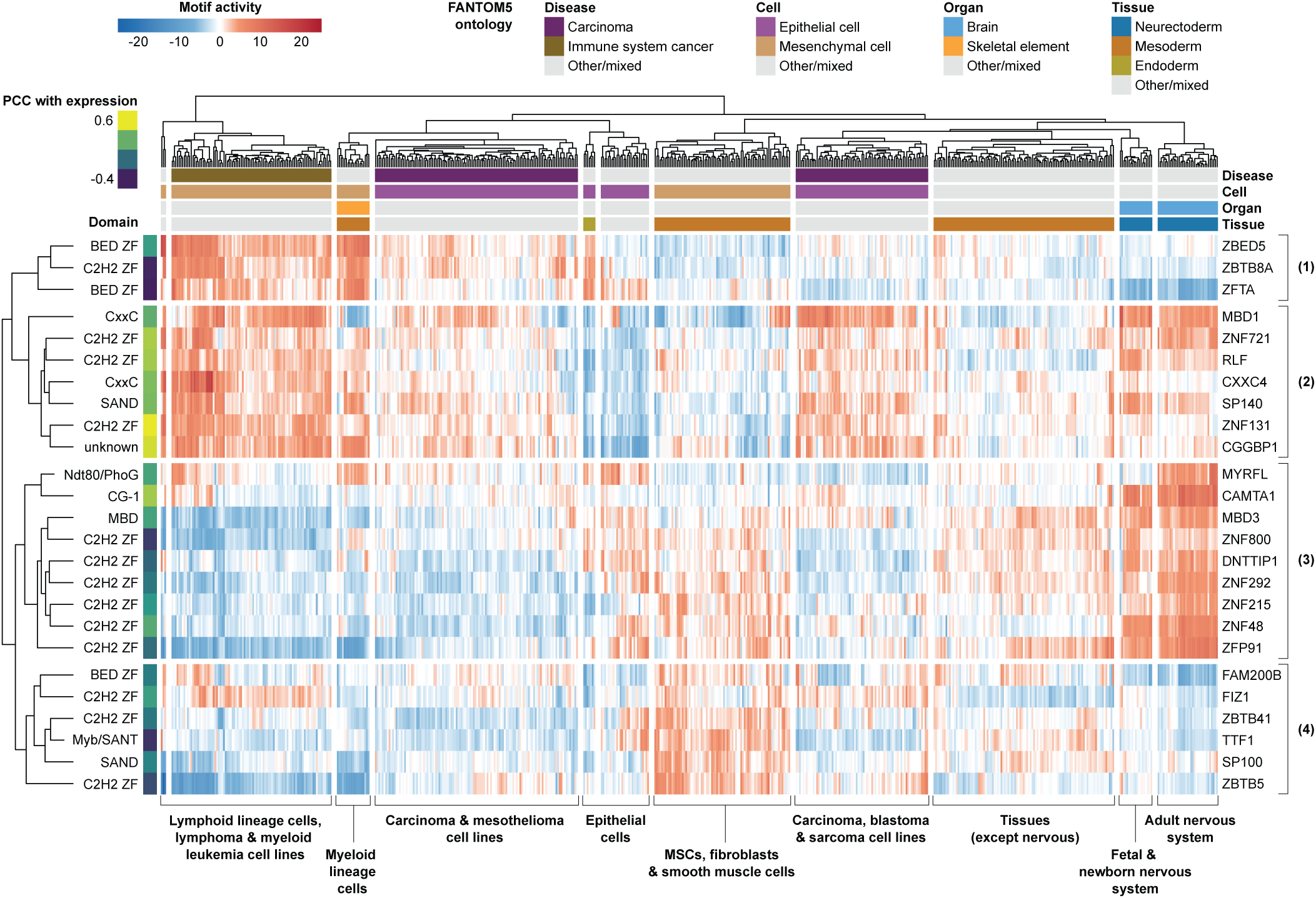
Motif activity response analysis highlights the Codebook motifs’ functionality. Heatmap of bi-clustered MARA-predicted motif activities. X-axis: 583 FANTOM5 samples, clusters are labeled manually according to the predominant cell and tissue types. Y-axis: TFs representing 25 Codebook motif clusters with the most significant differential activity across FANTOM5 samples. Additional columns, left-to-right: DNA-binding domain of the representative motif and the correlation between TF expression and motif activity. Extra rows, top-to-bottom: FANTOM5 ontology-based sample classification using the selected groups from disease state (DOID:4), cell type (CL:0000003), organ (UBERON:0000062), and tissue (UBERON:0000479) terms, clusters are colored by the predominant term if assigned to at least 40% of the samples. PCC: Pearson correlation coefficient, TF: transcription factor

We obtained similar outcomes when we analyzed the global patterns of activity among all 135 Codebook-only motif clusters and FANTOM5 samples using PCA: the first component mainly separates nervous system from all other samples, while the second one separates the developing nervous systems from adult (**Extended Data Fig. 8c**). Taken together, the motif activities of Codebook TFs distinguish FANTOM5 tissues and cell types, and provide many candidates for new transcriptional regulators of genes involved in the inflammatory response, cell cycle control, and nervous system development.

### Codebook TFs and the dark matter genome

TF binding sites in promoters and enhancers are easily rationalized as likely being involved in the regulation of adjacent genes. A principal result of the Codebook project is that dozens of Codebook TFs instead bind preferentially to other regions of the genome^13^. An accompanying manuscript^13^ focuses on the finding that roughly half of the examined TFs bind predominantly outside of promoters and enhancers in ChIP-seq, with most TFs having a very distinct set of binding sites that are supported by both GHT-SELEX and motif matches. Among these TFs are KRAB-containing C2H2 zinc fingers (KZNFs), which are best known for binding to specific classes of endogenous retroelements and nucleate regions of H3K9 trimethylation^59^. Until now, it has not been proven that the precise binding of KZNFs to specific retroelement subtypes is defined entirely by the sequence specificity of the KZNFs alone^60^, but the Codebook GHT-SELEX data abundantly demonstrate that this is indeed the case^12^.

The Codebook data also underscore the fact that transposons contribute to the TF repertoire itself^61^. Sixteen of the Codebook TFs (and two controls) that successfully yielded motifs possess a DBD that has itself been derived from a DNA transposase. These include CGGBP1^62^, six proteins containing BED-zf domains^63^, six with the related CENBP or Brinker domains^64^, three with transposon-derived Myb/SANT/MADF domains^65^, and FLYWCH1^66^. The PWMs obtained for CENPB/Brinker TFs are often long and information-rich, consistent with their presumed ancestral role in binding specifically to the transposons within a large genome (**Extended Data Fig. 9a, Supplementary Discussion**). A striking example is JRK, a TF that is found broadly in mammals^67^ and derived from an ancient domesticated Tigger DNA transposon^68^. All DNA transposons, including Tigger, have been transpositionally inactive in the human lineage for over 40 million years, and are presumably “fossils”^69^. Genomic binding of JRK is enriched for the same family of Tigger elements from which it appears to be derived (**Supplementary Discussion**). The consensus sequence for this Tigger element family has PWM-predicted binding sites for JRK in the terminal repeats (**Extended Data Fig. 9b**), consistent with its presumed ancestral role in transposition. To our knowledge, it is only the second human protein known to retain this quality (the other is SETMAR^70^). These cases apparently represent the simultaneous introduction of a multitude of *cis*-regulatory elements, and the TF that binds them, by the same transposon.

### Completing the human TF codebook

The primary purpose of Codebook was to assess 332 uncharacterized and putative human TFs for sequence-specific DNA binding activity, and produce trustworthy motifs for them, in order to assemble a complete human TF motif collection. This goal was mostly achieved. Codebook alone succeeded for slightly more than half of the tested TFs, and there is reason to believe that many of the unsuccessful putative TFs are likely not DNA-binding proteins or bind DNA with little or no sequence specificity (see

**Supplementary Discussion** and **Supplementary Document 1**). As noted above, a subset of the Codebook TFs, as well as other poorly characterized TFs, have also been analyzed by others since our study began. To evaluate the current scope of known human TF specificities, we surveyed other databases for PWMs for putative TFs that were not included in this study or were not found among the 177 Codebook successes. A conservative manual curation identifies 33 additional human TFs (i.e., beyond the 177 that were successful in Codebook) that have at least one plausible motif available in datasets that have been released since our 2018 TF census^1^ (**Supplementary Table 13**), leading to a new total of 1,421 human TFs with characterized sequence specificities (**Fig. 6** and **Supplementary Table 14**). **Supplementary Data 1** contains representative PWMs for all 1,421 TFs (see **Supplementary Table 10** for motif details).

**Figure 6.**
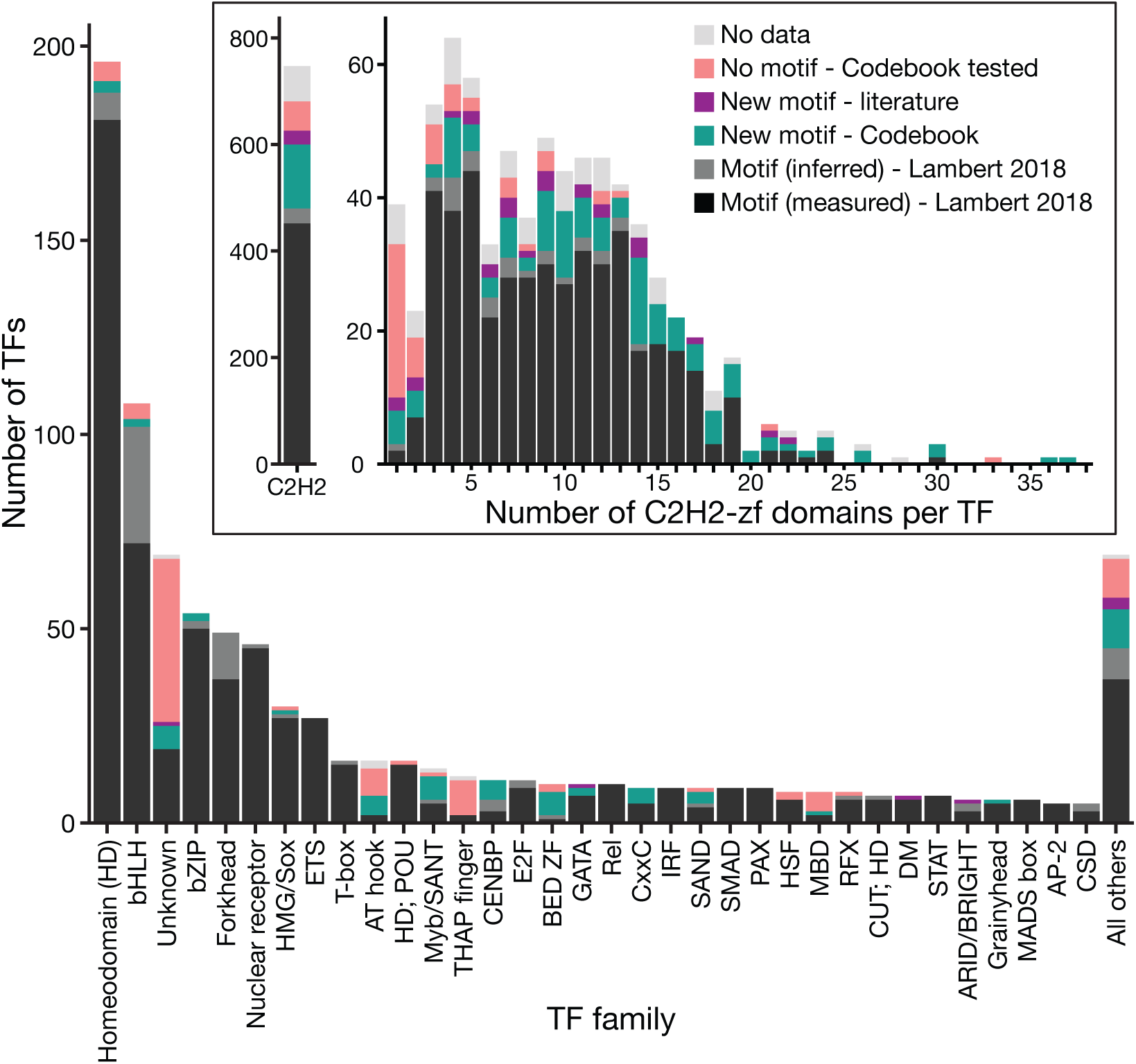
Motif coverage of human TFs by DBD family. TFs are categorized into structural classes from Lambert et al.^1^. See **Supplementary Table 14** for underlying information. TF: transcription factor, DBD: DNA-binding domain

This enumeration leaves 175 proteins, mostly with conventional DBDs, that lack known sequence specificity. Not all proteins with such domains are necessarily TFs, as noted above, and our re-curation of TFs that were unsuccessful in our assays suggests that 83 of these – nearly half – appear unlikely to be bona fide TFs (**Supplementary Document 1**). At the same time, new DBD classes continue to be identified (e.g. WH^71,72^). In addition, some TFs may bind only to methylated DNA, a possibility we explored in a separate manuscript in this collection^15^. Ongoing advances in the prediction of protein and protein-DNA structures^73^ have the potential to identify additional candidates for sequence-specific DNA binding. Thus, while completion of the objective to obtain a motif for every human TF now appears much closer, the list of likely human TFs continues to evolve.

An important technical demonstration of the Codebook project is that the simultaneous application of multiple experimental strategies and multiple motif-derivation and motif-scoring strategies was highly beneficial, with no method either dominating or dispensable^14^, consistent with previous benchmarking efforts^36,74^. As a result, many of the Codebook TFs are now among the best characterized human DNA-binding proteins in terms of their sequence specificity. In particular, obtaining *in vivo* and *in vitro* genomic binding profiles, together with assessing binding to randomly-generated artificial sequences, is advantageous in several ways: it facilitates disentanglement of direct and indirect binding, the contribution of the cellular environment, and the highly non-random nature of the genome sequence. Only for a small handful of the 1,000+ previously characterized TFs is there such a rich combination of data types. A much better perspective on human gene regulation and genome function, and evolution could presumably be obtained from the generation of such data for all human TFs.

## METHODS

### Plasmids and inserts

Sequences and accompanying information are given in **Supplementary Tables 2**, **3**, and **4**. Briefly, we selected Codebook TFs (and their DNA-binding domains) from information accompanying Lambert 2018^1^ and posted at https://humantfs.ccbr.utoronto.ca. Inserts named with an “-FL” suffix correspond to the full-length ORF of a representative isoform of the protein. Those with a “-DBD” suffix contain all of the predicted DBDs in the protein flanked by either 50 amino acids, or up to the N or C-terminus of the protein. Those with a “-DBD1”, “-DBD2” or “-DBD3” suffix contain a subset of the DBDs present in the proteins; these were designed manually, mainly for large C2H2-zf arrays. Inserts were obtained as recoded synthetic ORFs (BioBasic, US) flanked by AscI and SbfI sites, and subcloned into up to three plasmids: (i) pTH13195, a tetracycline-inducible, N-terminal eGFP-tagged expression vector with FLiP-in recombinase sites^10^; (ii) pTH6838, a T7-promoter driven, N-terminal GST-tagged bacterial expression vector^75^, and (iii) pTH16500 (pF3A-ResEnz-egfp), an SP6-promoter driven, N-terminal eGFP-tagged bacterial expression vector, modified from pF3A–eGFP^9^ to contain the two restriction sites after the eGFP.

### Protein production

Each experiment used a protein expressed from one of the following systems: (a) FLiP-in HEK293 cells (catalog number: R78007), induced with Doxycycline for 24 hours, used for inserts in pTH13195; (b) PURExpress T7 recombinant IVT system (NEB Cat.#E6800L), for inserts in pTH6838; or (c) SP6-driven wheat germ extract-based IVT (Promega Cat#L3260), for inserts in pTH16500.

### DNA binding assays

We followed previously-described protocols for ChIP-seq^10^, PBMs^36^, and SMiLE-seq^9^. Detailed descriptions of GHT-SELEX, HT-SELEX, ChIP-seq, and SMiLE-seq data collection and initial analysis are provided in the accompanying papers^14,15,76,77^. For PBMs, we analyzed proteins on two different PBM arrays (HK and ME), with differing probe sequences^78^.

### Data processing and motif derivation

The accompanying paper^14^ describes motif derivation and evaluation in detail. Briefly, after initial preprocessing, we obtained a set of ’true positive’ (likely bound) sequences for each individual experiment. (721 / 4,873) experiments were removed at this step, due to a low number of peaks, or other technical issues, as documented in **Supplementary Table 4**. We then applied a suite of tools (listed in **Supplementary Table 15**) to a training subset of the data from each experiment, and tested the resulting motifs on a test subset of the data from the same experiment, and also on the independent data for the same TF (i.e. the test sets from all other experiments done for the same TF). We employed a binary classification regime for all experiments and all motifs, and scored the motifs by a variety of criteria such as the areas under the receiver operating characteristic (AUROC) or the precision-recall curve (AUPRC).

### Systematic filtering of artefactual motifs

While curating the datasets and assembling a reliable motif collection, we accounted for enrichment of similar artifact motifs. These recurrent DNA patterns were detected due to systematic experimental noise or peculiarities of particular motif discovery tools. For example, in HT-SELEX experiments the “ACGACG” motif is often enriched. The sequence matches the constant flanking region of the ligand and likely abundant due to selective pressure for partially single-stranded DNA. In experiments with cell lysates, motifs of abundant native proteins in HEK293 cells were sometimes enriched (e.g., NFI, YY1, ETS-family TFs). To minimize the influence of these artifacts, we (**1**) manually compiled a list of known artifact motifs (**Supplementary Table 16**), (**2**) scanned the entire motif collection with MACRO-APE^79^, and removed motifs that were highly similar to those in our catalog of artifacts. Additionally, we removed motifs that matched the constant, non-variable regions of the DNA used in HT-SELEX, GHT-SELEX, and SMiLE-Seq. We did not filter out ETS-related motifs for ETS-family positive controls, such as ELF3, FLI1, and GABPA. Subsequent to the expert curation, we confirmed that enriched k-mers in HT-SELEX experiments did not correspond to potential artifacts associated with individual expression systems^76^.

### Motif discovery success rate evaluation

To estimate the success rate of different motif discovery tools across different platforms, we started with the successful experiments and TFs. For each combination of a platform X (e.g. ChIP-seq) and a motif discovery tool Y (e.g. Autoseed), we computed the number of experiments (**Extended Data Fig. 1c**) or TFs (**Extended Data Fig. 1d**) that yielded a highly similar motif to the reference motif (i.e. manually curated) in that combination. For that, for a specific experiment or TF, we took the whole set of candidate motifs generated by that combination of platform X and tool Y, and checked whether any of those motifs pass the similarity threshold using a one-pass scan of MACRO-APE^79^ ScanCollection command with the following parameters: -c 0.05 --rough-discretization 10 --precise 100500. Note that prior to comparison, the motifs were converted to log-odds PWMs (as in ^14^).

### Experiment evaluation by expert curation

To gauge the success of individual experiments, we employed an “expert curation” workflow with an initial voting scheme in which a committee of annotators gauged whether individual experiments should be “successful”, i.e. included in subsequent analyses. All experiments were examined by at least three annotators. A subcommittee (AJ, IVK, and TRH) jointly resolved all cases of disagreement among initial annotators (∼300 experiments) and then reviewed all successful experiments. Annotators had available an early version of the MEX portal (https://mex.autosome.org, described in ^14^) containing results of all PWMs scored against all experiments, and were tasked with gauging whether the experiments yielded PWMs that were similar across experiments, or scored highly across experiments.

Annotators also considered whether the motif was consistent with those for other members of their protein family (e.g. BHLHA9 yielded an E-box-like motif, CAnCTG), and/or similar between closely related paralogs (e.g. ZXDA, ZXDB, and ZXDC all yielded similar motifs). We also considered whether (and how many) “peaks” were obtained from ChIP-seq or GHT-SELEX, and whether these peaks were common to independent experiments (e.g. both ChIP-seq and GHT-SELEX). Annotators were further given a measure of similarity between Codebook PWMs and any PWMs in the public domain, as well as enrichment of known or suspected common contaminant motifs in any experiment.

### Post-evaluation peak processing

After identification of successful experiments, we re-derived peak sets for ChIP-seq and GHT-SELEX experiments in order to obtain a single peak set for each TF, as described in the accompanying papers^12,13^. Briefly, for ChIP-seq we repeated the peak calling using MACS2 and experiment-specific background sets, using a procedure previously described^10^, then merged the peak sets for replicates of the same TF with BEDTools merge^80^ (see accompanying manuscript^13^: “ChIP peak replicate analysis and merging”). We derived GHT-SELEX peaks using a novel method, MAGIX, that calculates enrichment of reads in each cycle, and treats different experiments as independent statistical samples in order to obtain a single enrichment coefficient per peak^12^.

### Expert motif curation

For this study, to identify a single representative PWM for each TF, we first compiled a set of the highest-scoring candidate PWMs for each TF (as summarized above and elsewhere^14^), then ran additional tests with them, utilizing the reprocessed peak data, and manually evaluated the outputs. We started with the union of three sets of 20 PWMs for each TF: the 20 PWMs with the highest AUROC (as calculated elsewhere^14^) on (i) any successful ChIP-seq experiment for the given TF, (ii) any successful GHT-SELEX experiment for the given TF, and (iii) any successful HT-SELEX experiment for the given TF. These PWMs were selected regardless of the data set from which they were derived. We then reassessed these PWMs against ChIP-seq and GHT-SELEX data with two parallel methodologies. First, we recalculated AUROC for each of the candidate top PWMs on the merged, thresholded sets of ChIP-seq peaks (P < 10^-10^)^13^ using AffiMX^31^ to score each peak. We generated negative sets using BEDTools shuffle^80^ with the *-noOverlapping* option to create sets of random genomic regions with the same number of peaks, and the same peak width distribution as the corresponding ChIP peak sets. We used the same technique to calculate AUROC values for GHT-SELEX, with thresholded peak sets (using a “kneedle”^81^ specificity value of 30 in the sorted enrichment values^13^). In parallel, we calculated the Jaccard index to measure the overlap between PWM matches (identified by MOODS^82^ with -p 0.001) vs. ChIP-seq peaks and GHT-SELEX peaks, as two separate measures. The overlap in each case was maximized by applying different thresholds on the peak sets and choosing the cutoff at which the Jaccard index was the highest^12^. We then applied expert curation (by a committee consisting of AJ, TRH, KUL, AF, RR, MA, and IY) to choose a single representative PWM with high performance on all compiled scores that, all else equal, also reflected reasonable expectation from the DBD class (including recognition-code predicted motifs, see accompanying manuscript^12^) and did not have low information content.

### Motif similarity analysis

We took two different approaches to determine the number of distinct motifs represented by the Codebook TFs, and the number of novel motifs added to the human TF repertoire. In the first approach, we identified the similarity between 1,582 PWMs representing the Codebook TFs (177 PWMs, 177 TFs) and TFs with previously known specificities (1,405 PWMs, 1,211 TFs). The Codebook TF PWM set is the set of representative PWMs for the 177 Codebook proteins with identified motifs. For TFs with previously known specificities, the set of PWMs identified as the “best” in Lambert et al. 2018^1^ was used, except for two TFs (MTF2 and PHF1) that did not have an assigned PWM. For these two TFs, the PWMs were retrieved from CisBP^83^. We used the correlation between pairwise affinities to 150,000 random sequences of length 100, as calculated by MoSBAT^31^, as the metric for PWM similarity. We then clustered the 177 Codebook TFs based on these PWM similarities with Pearson correlation and average linkage and identified the ideal number of clusters (129) by determining the optimal silhouette value^84^, which corresponded to splitting PWMs into clusters at a distance of 0.76. We then clustered the full set of 1,582 motifs and used the same distance to split the PWMs into clusters, resulting in 613 clusters, 92 of which contained only Codebook TFs. The set of motifs used in the clustering and their cluster membership are provided in **Supplementary Table 10**.

Independently, and to obtain a non-redundant set of motifs for MARA analysis, we followed the procedure as in^85^ with the addition of Codebook motifs. First, we merged HOCOMOCO^38^ v12 motifs with those evaluated in Codebook, preferably selecting those built with ChIPMunk and ranking high in benchmarking^14^ to maintain consistency with HOCOMOCO v12, which was fully built with ChIPMunk. Having the joint motif collection, we estimated the motif similarities with MACRO-APE^79^ at the motif P-value cutoff of 0.0005 and a default matrix discretization parameter (-d) of 1 (increased to 10 for improved precision only for motif pairs with the Jaccard similarity over 0.01 at -d 1). Next, using the pairwise motif similarity matrix, we performed agglomerative clustering (’average’ linkage) with sklearn. The number of clusters was taken to maximize the silhouette score. Clusters of low-quality motifs (HOCOMOCO ’D’) were discarded. Finally, for each cluster, a single representative motif was taken according to the highest average similarity to all other motifs in the cluster (**Supplementary Table 11**). The non-redundant set of representative motifs is available for download from Zenodo (https://zenodo.org/records/17307845) [doi:10.5281/zenodo.17307845], and the motif clusters annotation is available in **Supplementary Table 11.**

### Motif degeneracy analysis

To explore if low information content (IC) is an intrinsic feature of some binding motifs, we adjusted the IC of PWMs and tested prediction accuracy of ChIP-seq and GHT-SELEX binding sites. PWM IC was adjusted on a per-base-pair basis, by iteratively scaling probabilities for each base at each position until the PWM reached an average IC of 1 bit per base pair. The script, “logo_rescale.pl”, is available at https://gitlab.sib.swiss/EPD/pwmscan.

### Comparison to external peak sets and PWMs

For comparison, we downloaded ChIP-seq peak sets from GTRD^86^ and ENCODE (4.12.2023)^87^, for all Codebook TFs. We then divided these data into four categories corresponding to the cell type: HEK293/HEK293T, HepG2, K562, and other cells. We preferentially selected the peak sets from GTRD, because it contained more diverse peak sets: since GTRD has processed the majority of ENCODE consortium experiments, together with many non-ENCODE experiments. When multiple experiments were available for a TF in a cell type category, we selected the experiment with higher peak counts. If multiple computational methods had been used to derive peak sets for the selected experiment, we chose the peak set using a preferential order MACS, GEM, SISSRS, PICS and PEAKZILLA. See **Supplementary Table 7** for identifiers and metadata of the reference datasets.

For PWM scoring, the external peak sets were used as downloaded, with the exception of peak sets that were generated with the GEM peak caller, which have a peak width of 1 (summit only), and were therefore expanded 250 bases in both directions. For Codebook data, we used the merged and thresholded Codebook ChIP peak sets as in “Expert motif curation”. We generated negative peak sets for each ChIP-seq peak set using BEDTools shuffle^80^ with the *-noOverlapping* option to create sets of random genomic regions with the same number of peaks and the same peak width distribution as the corresponding ChIP peak sets. We downloaded PWMs for all Codebook TFs from JASPAR^88^ (2024 version), HOCOMOCO^38^ (v12) and Factorbook^89^ (downloaded 15.12.2023) (**Supplementary Table 13**). We scanned Codebook and external peak sets (and corresponding negative sets) with the representative (i.e., expert-curated) Codebook motifs (PWMs) using AffiMX^31^, and calculated AUROC values. Additionally, for the 19 Codebook TFs with a successful Codebook ChIP-seq experiment, a Codebook PWM, an external ChIP-seq experiment, and an external PWM, we compared the performance of PWMs across the different peak sets as follows. We first selected a single external PWM for each of the 20 TFs by scanning each PWM for a given TF on each external peak set for the same TF and identifying the PWM that produced the highest AUROC. We then used these highest scoring PWMs to scan the corresponding Codebook data and calculate AUROC values.

### Curation of external motifs for previously uncharacterized TFs

For this analysis we considered all proteins that were annotated as putative TFs in Lambert et al. 2018^1^ but did not then have a credible motif. We downloaded all motifs for these TFs from JASPAR^88^ (2024 version), HOCOMOCO^38^ (v12) and Factorbook^89^ (downloaded 15.12.2023), resulting in a set of 484 PWMs, that were then assessed manually by AJ considering: 1) Whether similar motifs were obtained for the protein from multiple independent datasets; 2) If the motif was consistent with the structural class of the TF and; 3) If the motif was likely to describe inherent specificity of the protein and not, for example, reflect a target site of another TF or a likely artifact. The curated motifs are listed in **Supplementary Table 13** with more details in **Supplementary Table 10**. The PWMs themselves are available in **Supplementary Data 1**, and examples of artifactual and correct external motifs are shown in **Extended Data Fig. 10**.

### Motif activity response analysis (MARA)

The promoter locations and activity data were taken from FANTOM5^47^. After filtering (see **Supplementary Methods**), the set comprised 209,374 individual promoters and 1,020 samples (including replicates), belonging to 583 unique samples (142 tissues, 187 primary cells, and 254 cell lines, see **Supplementary Table 12** and Zenodo [https://zenodo.org/records/17307845]). The sum-occupancy scores^90^ were computed in [250 bp upstream; 10 bp downstream] regions relative to the FANTOM5 TSS locations by scanning with a subset of 632 non-redundant motifs representing TFs with expression detectable in the FANTOM5 data.

We employed MARADONER (MARA-done-right) v 0.13, a command-line Python reimplementation of MARA available in PyPI (pip install maradoner). Conceptually, MARADONER extends the original logic of MARA^45,46^ to model the promoter activity in each sample as a linear function of the motifs’ scores and sample-specific motif activities, where each motif represents a set of TFs with shared binding specificity. Compared to the classical MARA, MARADONER explicitly models sample and motif means (see **Supplementary Methods** for more details).

### TOP (Triple Overlap) and CTOP (Conserved Triple Overlap) peak set analyses

To obtain the TOP sites, we first identified thresholds for ChIP-seq peaks, GHT-SELEX peaks, and PWM-derived “peaks” (see below) that maximize the three-way Jaccard metric (overlap/union) of the three sets, with the thresholds calculated for each TF independently. We converted PWM matches (derived from MOODS^82^ using a P-value cut-off of 0.001) into peaks by merging neighboring matches with a distance less than 200 bp and re-scoring them using the sum-occupancy for clusters. We then identified TOPs as peaks exceeding these thresholds in all three sets and overlap in all three. To obtain the CTOP sites, we then extracted PhyloP scores for each base at each TOP site (and 100 flanking bases) from the Zoonomia consortium^91^, removed sites overlapping the ENCODE Blacklist^92^ or protein coding sequences (due to the skew in phyloP scores caused by codons), and applied three different statistical tests for significance of phyloP scores over the PWM match: two that tested correlation between the IC and the phyloP value at each base position of the PWM (using either Pearson correlation or linear regression), and one that tested for higher phyloP scores over the PWM match (Wilcoxon test). Greater detail on these specific operations is given in the accompanying manuscripts^12,13^.

### Intersection of TOPs/CTOPs and genomic features

We first clustered all CTOPs using BEDTools merge^80^, with a max distance of 100 bp, then intersected with the following genomic feature sets: basic canonical protein coding promoters from GENCODE v44^93^, defined as 1000 bp upstream and 500 bp downstream of the canonical TSS; the “Unmasked CpG Island” track, PhastCons Conserved Elements from the Multiz 470 Mammalian alignment, and RepeatMasker track from UCSC^94^; ChromHMM HEK293 enhancers^13^. We classified promoters as CpG island or non-CpG island based on the GENCODE basic TSS being within +/- 50 bp of a CpG island from the unmasked track. We classified the CTOP clusters as associated with a single type of genomic feature in the following order of priority: CpG island associated with a protein coding promoter; other CpG islands; a non-CpG island-associated protein-coding promoter; an enhancer; clusters containing a CTCF binding site but not overlapping a CpG island, promoter or enhancer; overlapping a transposable element and none of the previous categories; overlapping a non-TE repeat and none of the prior categories; and “Other” for CTOP clusters not intersecting any examined features.

### Analysis of allele-specific binding

The ASB analysis was performed as in Buyan et al.^27^. Briefly, we performed SNP calling with bcftools^95^, corrected mapping bias with *WASP*^96^, generated the maps of background allelic dosage with *BABACHI*^97^ (Abstract O3), called ASBs with *MIXALIME*^27^, and performed PWM scoring with *PERFECTOS-APE*^98^, see **Supplementary Methods** for details. To obtain the non-redundant set of ASB-SNPs across Codebook data for intersecting with existing regulatory SNP databases (GTEx v.8^30^, ADASTRA v.6.1^28^, and EBI GWAS Catalog v1 e115_r2025-12-03_full)^29^, we performed a joint aggregation of the allelic imbalance P-values over all processed datasets (ChIP-Seq and GHT-SELEX combined).

## Supporting information

Supplementary Discussion & Methods

## DATA AVAILABILITY

The sequencing raw data for the ChIP-seq, GHT-SELEX, and HT-SELEX experiments have been deposited into the SRA database under identifiers PRJEB78913 (ChIP-seq), PRJEB76622 (GHT-SELEX), and PRJEB61115 (HT-SELEX). Genomic interval information generated for ChIP-seq and GHT-SELEX have been deposited into GEO under accessions GSE280248 (ChIP-seq) and GSE278858 (GHT-SELEX). PWMs can be browsed at https://mex.autosome.org and downloaded at from Zenodo (https://zenodo.org/records/15667805) [doi:10.5281/zenodo.15667805]. An updated list of human TFs is available at https://humantfs.ccbr.utoronto.ca. The final motifs generated in this study are available in build 3.0 of the CisBP database^99^. Information on constructs, experiments, analyses, processed data, comparison tracks, and browsable pages with information and results for each TF is available at https://codebook.ccbr.utoronto.ca. SNP calls and ASBs are available at Zenodo (https://doi.org/10.5281/zenodo.18224872) [doi:10.5281/zenodo.18224871]. The data underlying MARA are available at Zenodo https://zenodo.org/records/17307845.

## CODE AVAILABILITY

The script for adjusting PWM information content, logo_rescale.pl, is available at https://gitlab.sib.swiss/EPD/pwmscan, the code for the identification of TOPs and cTOPs is available at https://github.com/imyellan/Codebook_CTOP_scripts. The code used for the SNP-centric analysis is available at GitHub: https://github.com/autosome-ru/perspectives-on-codebook-allele-specificity, the MIXALIME is available on GitHub: https://github.com/autosome-ru/mixalime. MARADONER is available on GitHub: https://github.com/autosome-ru/MARADONER.

## ACKNOWLEDGEMENTS

We thank the IT Group of the Institute of Computer Science at Halle University for computational resources, Maximilian Biermann for valuable technical support, Gherman Novakovsky for providing feedback, Berat Dogan for testing earlier versions of RCADEEM, and Debashish Ray for assistance with database depositions.

This work was supported by the following:

- Canadian Institutes of Health Research (CIHR) grants FDN-148403, PJT-186136, PJT-191768, and PJT-191802, and NIH grant R21HG012258 to T.R.H
- CIHR grant PJT-191802 to T.R.H. and H.S.N.
- Natural Sciences and Engineering Research Council of Canada (NSERC) grant RGPIN-2018-05962 to H.S.N.
- A Swiss National Science Foundation grant (no. 310030_197082) to B.D.
- Assignment 125091010189-3 to I.V.K.
- MSHERF grant number no. 075-15-2025-014 (previously no. 075-15-2024-666)
- Marie Skłodowska-Curie (no. 895426) and EMBO long-term (1139-2019) fellowships to J.F.K.
- NIH grants R01HG013328 and U24HG013078 to M.T.W., T.R.H., and Q.D.M.
- NIH grants R01AR073228, P30AR070549, and R01AI173314 to M.T.W.
- NIH grant P30CA008748 partially supported Q.M.
- Canada Research Chairs funded by CIHR to T.R.H. and H.S.N.
- Ontario Graduate Scholarships to K.U.L and I.Y.
- A.J. was supported by Vetenskapsrådet (Swedish Research Council) Postdoctoral Fellowship (2016-00158)
- The Billes Chair of Medical Research at the University of Toronto to T.R.H.
- EPFL’s Center for Imaging
- Resource allocations from Digital Research Alliance of Canada

## AUTHOR CONTRIBUTIONS

T.R.H. conceived of the Codebook Consortium. T.R.H. and B.D. directed the laboratory work. I.V.K. led the GRECO-BIT data analysis team. A.J., O.F., V.J.M., J.G., I.G., P.B., B.D., I.V.K., and T.R.H. oversaw the data analysis. Experiments were performed by: A.J. and A.W.H.Y (HT- and GHT-SELEX); A.W.H.Y (PBM); R.R., A.Bo., A.Br., H.Z., and M.B (ChIP-seq), and A.J.G, J.F.K.S., S.I, A.W.H.Y., and B.D. (SMiLE-seq). J.G., P.B., O.F., V.J.M., I.G., I.K., I.V.K., K.U.L., A.F., S.A., A.Br., A.Bo., J.F.K.S., A.J.G., M.A., M.T.W., I.V., and the Codebook Consortium members contributed to PWM derivation and motif analysis. All authors contributed to the initial assessment of experiments. I.V.K., A.J., and T.R.H. performed the secondary assessment of experiments. K.U.L, A.F., I.Y, V.N., A.B., G.M., and A.J. performed statistical analysis and created illustrations. The paper was drafted and finalized by T.R.H., A.J., K.U.L., A.F., I.Y., V.N., A.B., V.J.M., P.B., and I.V.K. All authors contributed to data analysis and reviewed the manuscript.

## DECLARATION OF COMPETING INTERESTS

O.F. is employed by Roche.

---

**Extended Data Figure 1.**
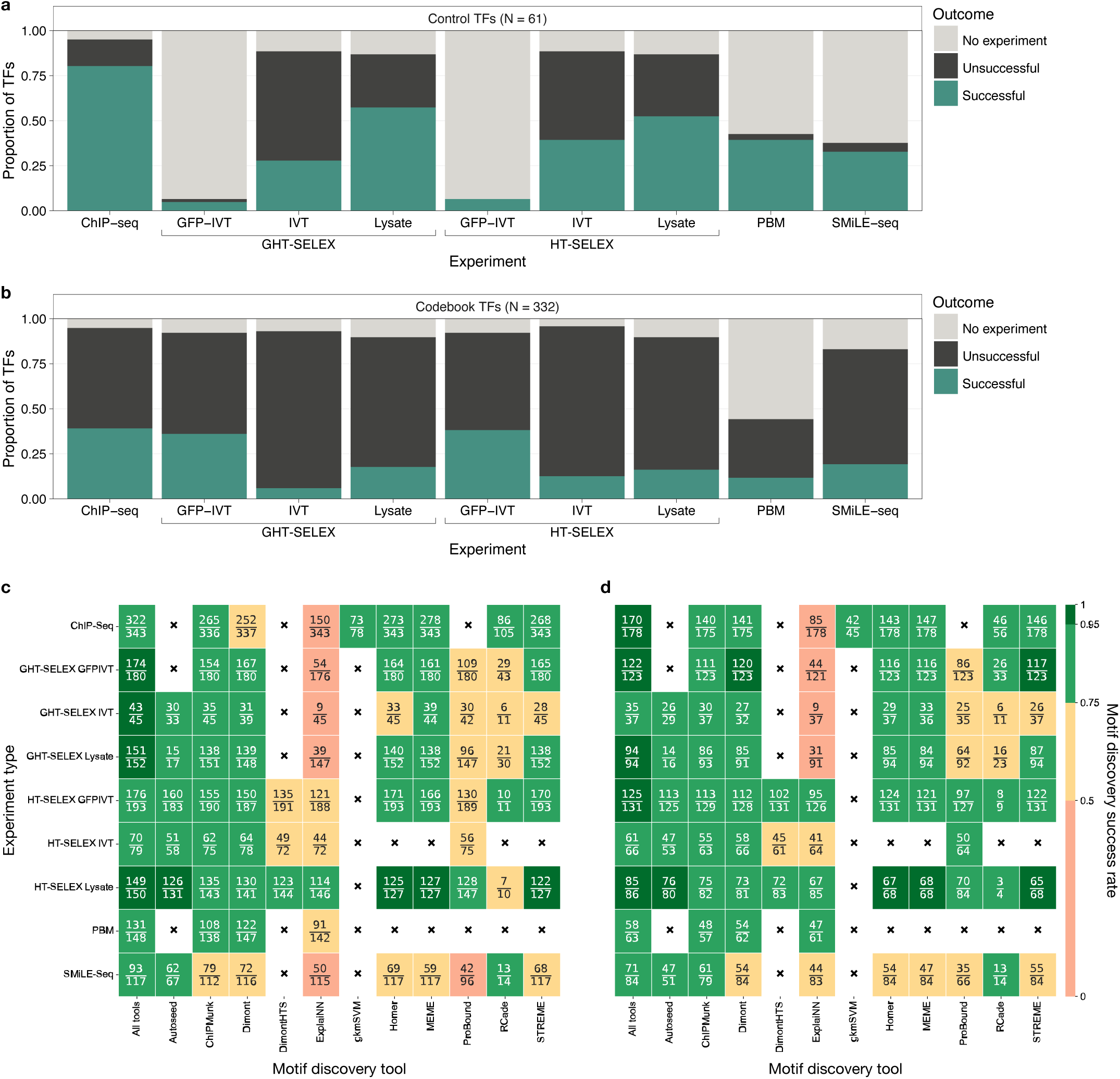
Motif discovery success rates. **a**, **b**, Proportion of control (**a**) and Codebook (**b**) TFs for which a motif was successfully identified across experiment types. GHT-SELEX and HT-SELEX experiments are separated according to protein expression system. The proportion of untested TFs is also indicated. **c, d**, Rate of successful motif discovery across experiment types and motif discovery tools. Numerator in each cell represents the number of successes (cases of correctly identified motifs), and denominator represents the number of experiments (**c**) or TFs (**d**) for a particular combination of a tool and an assay. The first column in **c** and **d** displays the joint estimates across all tools. IVT: in vitro transcription, TF: transcription factor

**Extended Data Figure 2.**
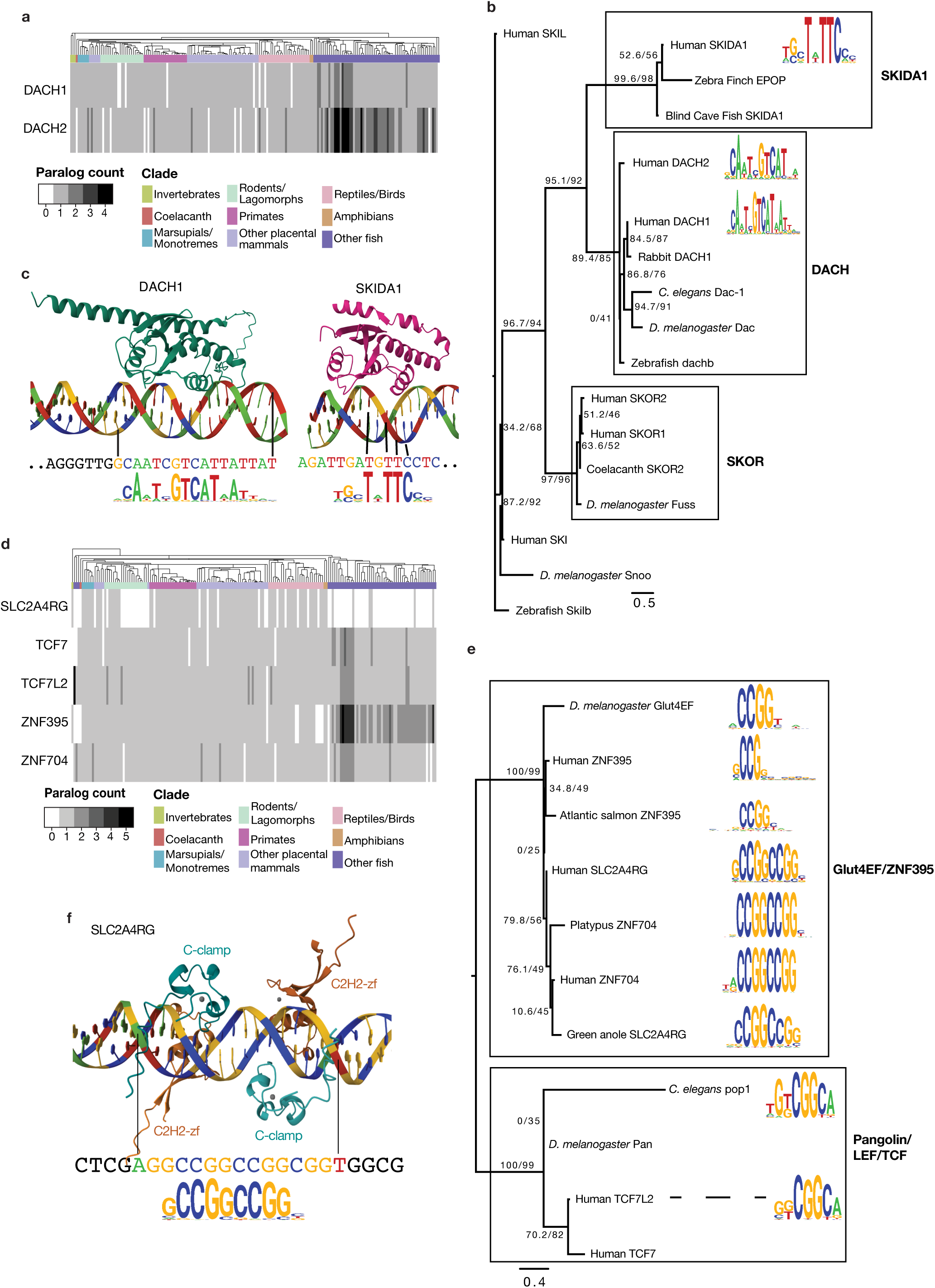
Undercharacterized DNA-binding domains. Overview of new motifs for undercharacterized TF families. **a,** Number of DACH1 and DACH2 orthologs (union of one-to-one and one-to-many) across Ensembl v111 vertebrates and selected invertebrates. Species order reflects the Ensembl species tree. **b**, Sequence logos (from the main Codebook project, and follow-up study described in the **Supplementary Discussion**) and sequence relationships of human and representative animal SKI/SNO/DAC domains. All motifs from successful HT-SELEX assays are shown, and a representative set of the DBDs tested is included in the phylogenetic tree. Tree is an unrooted maximum-likelihood phylogram from IQTree^100^ using parameters --alrt 1000 -B 1000 (1000 replicates to compute SH-aLRT (a branch support value from 0-100%), 1000 replicates for ultrafast bootstrap), and automatic best-fit model selection with ModelFinder. Branch values are SH-aLRT/ultrafast bootstrap. **c,** AlphaFold3-predicted structure of the DACH1 SKI/SNO/DAC region (residues 130 – 390) bound to an HT-SELEX ligand sequence with a high-scoring PWM match. **d,** Number of orthologs (same counting process as **a**) for human C-clamp proteins. **e**, Sequence logos (from the main Codebook project and follow-up study) and sequence relationships of human and representative animal C-Clamp domains (*ZNF704 motif from ^101^). The tree was produced identically to the tree in **b**. **f,** AlphaFold3-predicted structure of two full-length SLC2A4RG proteins bound to a CTOP sequence with flanking sequences (chr17:48,048,369-48,048,401), and four Zn**^2+^** ions (grey). The remainder of the proteins (beyond the C-clamp and C2H2-zf domains) are hidden, for visual simplicity. DBD: DNA-binding domain

**Extended Data Figure 3.**
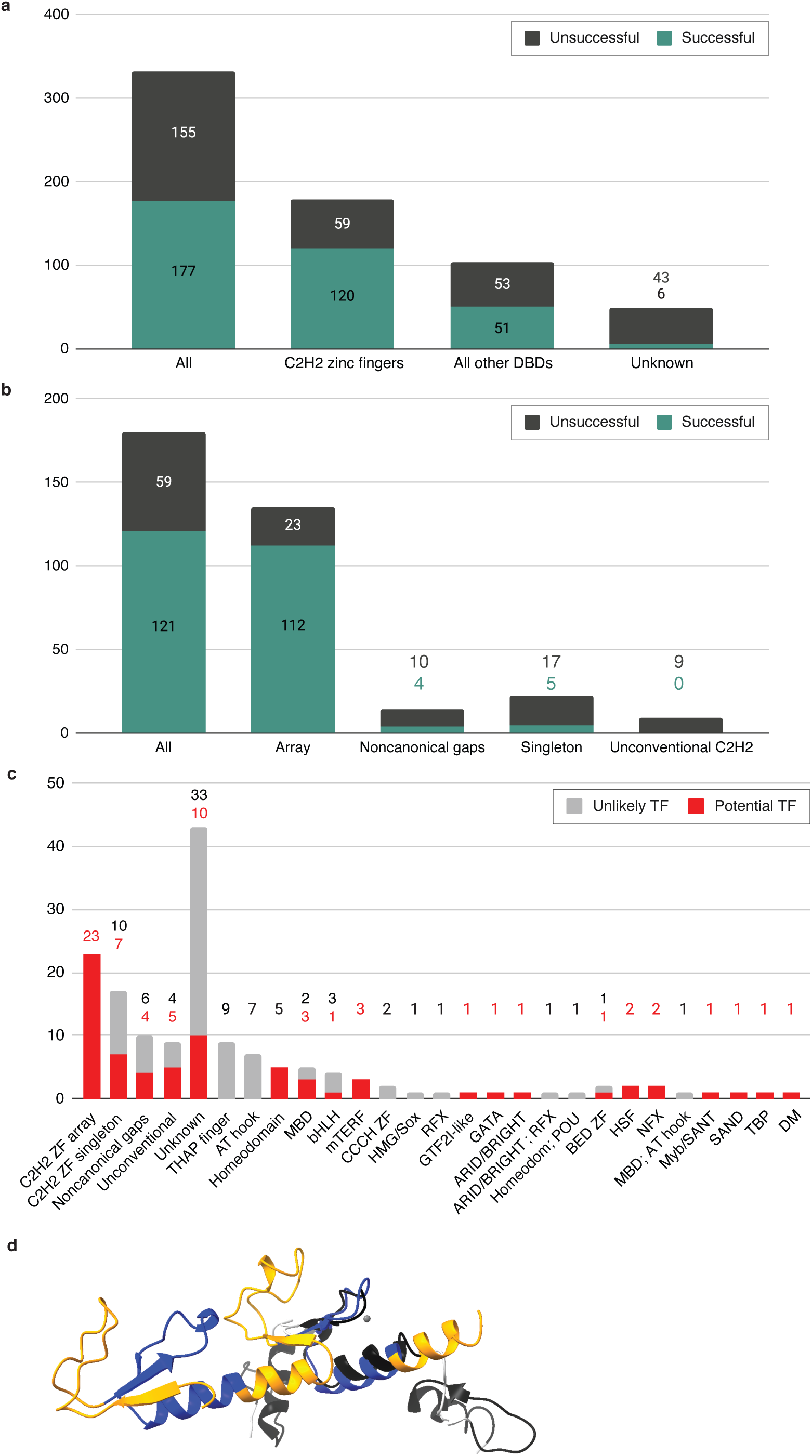
Description and re-assessment of unsuccessful proteins. **a**, Number of successful and unsuccessful Codebook proteins by category. **b,** Number of successful and unsuccessful C2H2-zf proteins categorized based on: whether a protein has an array of at least two DBDs separated by 3-10 AA; has multiple C2H2-zf domains but with other kinds of spacings; has only a single domain; or if it has a potential C2H2-zf domain based on literature but none are predicted by our standard HMM based prediction. **c,** Summary of re-assessment (see **Supplementary Table 6** and **Supplementary Document 1** for details) of unsuccessful proteins by structural class or category. Colors indicate if the protein is still assessed as a potential TF (red) and is therefore a potential false negative, or if it is now assessed as an unlikely TF (gray). **d**, shows a comparison of the CASZ1 “para-ZNF” domain^102^ zinc fingers 1 and 2 against an example of canonical C2H2-zf domain of (GLI2, PDB:2GLI), with the predicted C2H2-zf region CASZ1 DBD2 structurally aligned to DBD4 of GLI2. TF: transcription factor, DBD: DNA-binding domain, HMM: hidden Markov model

**Extended Data Figure 4.**
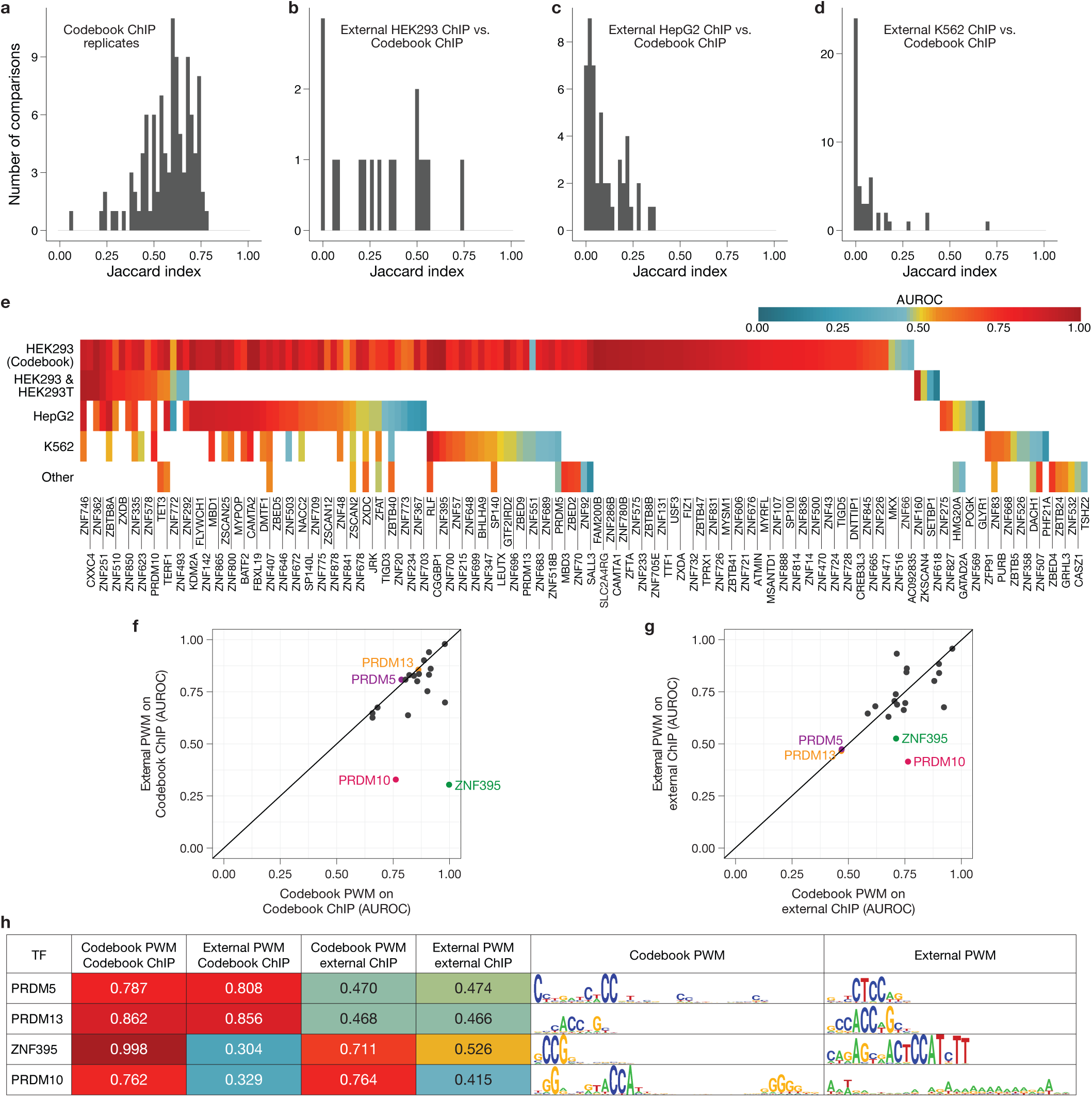
Comparison of Codebook ChIP-seq datasets and PWMs to external ChIP-seq datasets and PWMs. a-d, Histograms of optimized Jaccard indices measuring the overlap between ChIP-seq peak sets for the same TF: **a,** Codebook ChIP-seq replicates; **b, c, d**: Codebook ChIP-seq vs. external ChIP-seq performed in HEK293 cells (**b**), HepG2 cells (**c**), or K562 cells (**d**). Control TFs are not included. **e,** AUROC values for expert-curated Codebook PWMs (columns), discriminating ChIP-seq peaks vs. random genomic background loci. Rows show different cell types. **f, g,** comparison of Codebook and external PWMs at the task of discriminating ChIP-seq peak sets from random sequences (as in **e**), for the 20 TFs that have Codebook ChIP-seq data, a Codebook PWM, external ChIP-seq data, and an external PWM. **f**, Performance of Codebook PWM vs. the best performing external PWM on Codebook ChIP-seq data. **g**, Performance of Codebook PWM vs. the best performing external PWM on external ChIP-seq data. The four TFs with an AUROC of < 0.5 on either axis of either plot are highlighted. **h,** Sequence logos for the four TFs highlighted in **f** and **g**. All expert-curated Codebook PWMs displayed are supported by ChIP-seq, GHT-SELEX, HT-SELEX, and SMiLE-seq data. PWM: position weight matrix, TF: transcription factor, AUROC: area under the receiver operating characteristic

**Extended Data Figure 5.**
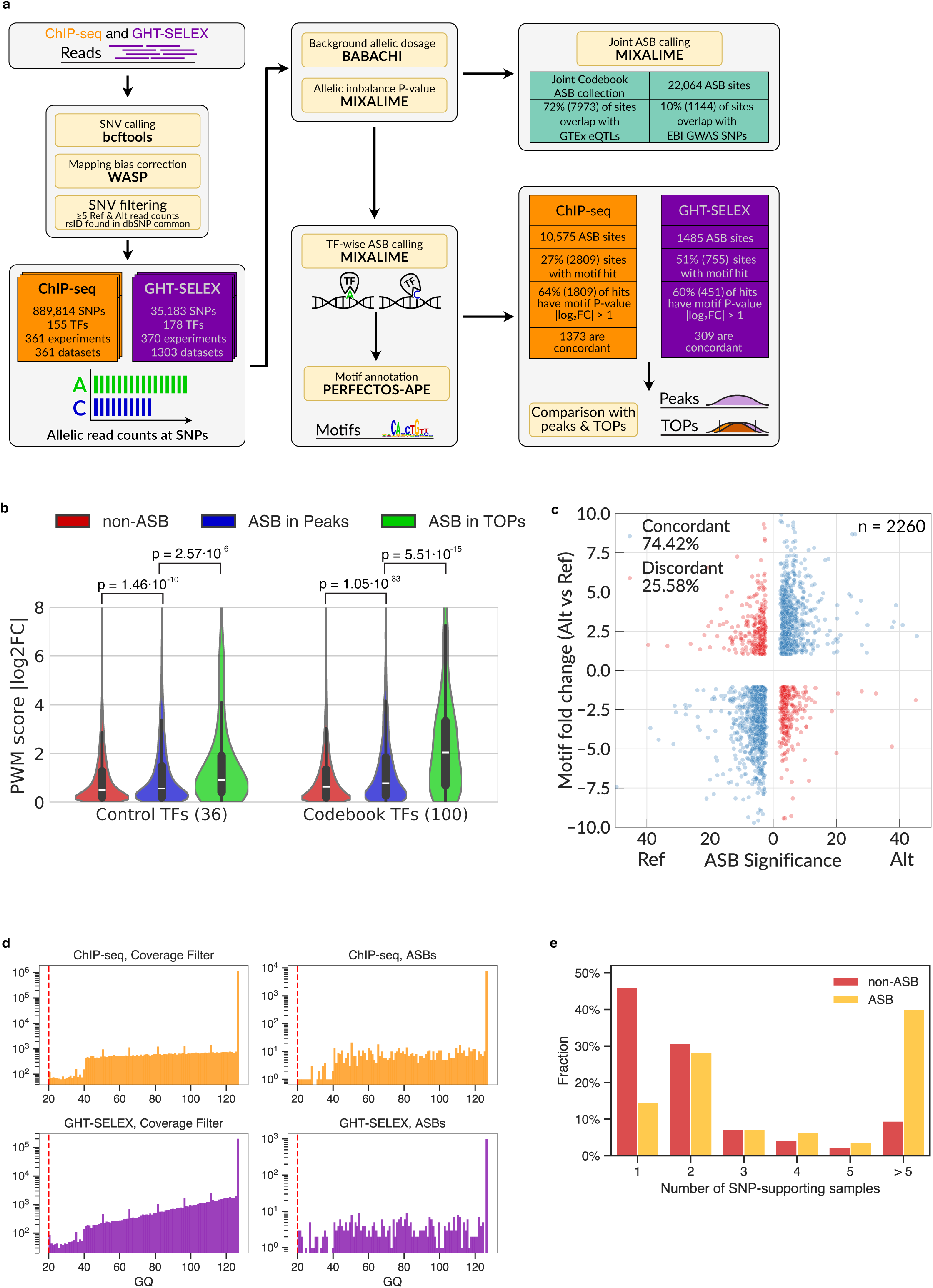
Identifying allele-specific TF binding sites in Codebook ChIP-seq and GHT-SELEX data. a, Principal scheme of the allele-specific analysis. **b,** Distribution of PWM score P-value ratio (fold change), |log_2_ FC|, between alleles for non-ASBs (FDR ≥ 5%), ASBs (FDR < 5%) in peaks, and ASBs in TOPs (FDR < 5%). *Left*, 36 positive control TFs; *Right*, 100 Codebook TFs. P-values: Mann-Whitney U test. **c,** Motif concordance of Codebook ASBs. X-axis: ASB allelic preference and significance (-log_10_ FDR), *left*: preference for the reference allele (Ref), *right*: preference for the alternative allele (Alt). Y-axis: PWM score P-value ratio (log_2_ FC) between Alt and Ref. The plot shows only ASBs with |log_2_FC| > 1. **d,** Genotype quality (GQ) of SNPs called from ChIP-Seq (top) and GHT-SELEX (bottom) data. X-axis: GQ, Y-axis: number of SNPs or ASBs. Red dashed line at 20 is the default GQ threshold used to filter SNP calls. The majority of candidate SNPs and ASBs have GQ much higher than the default threshold**. e,** The number of SNP-supporting ChIP-Seq or GHT-SELEX datasets for ASB and non-ASB SNPs. The majority of ASB SNPs are supported by two or more datasets. ASB: allele-specific binding site, FC: fold change, SNP: single-nucleotide polymorphism, TOP: triple-overlap

**Extended Data Figure 6.**
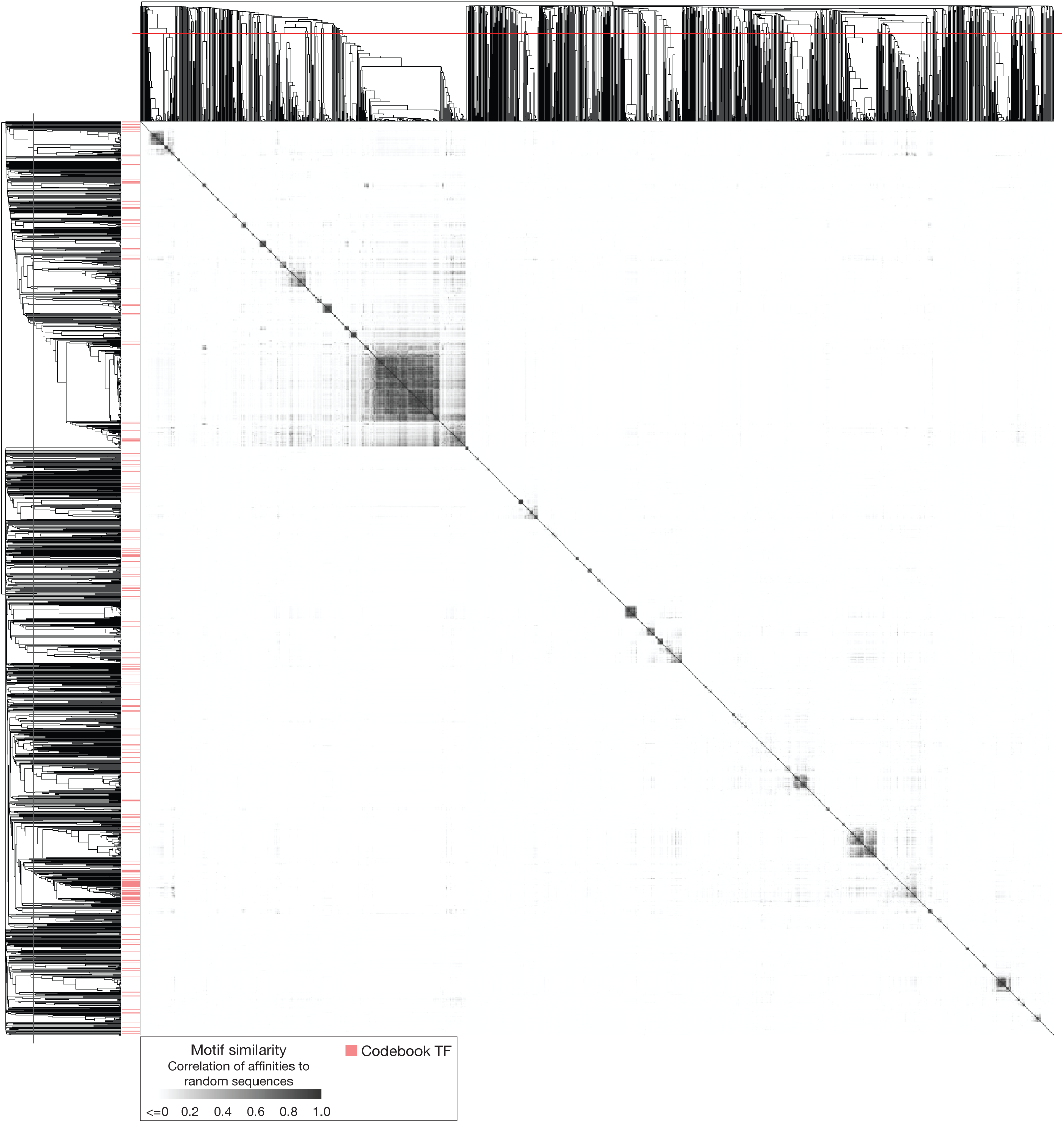
Similarity of TF motifs. Symmetric heatmap displays the similarity between 1,582 PWMs clustered by Pearson correlation with average linkage. The PWM set is composed of the 177 representative PWMs for Codebook TFs, and 1,405 PWMs for 1,211 TFs with previously identified binding specificities. The set of PWMs for non-Codebook TFs was retrieved from Lambert et al. 2018^1^. PWM similarity is the correlation between pairwise affinities to 150,000 random sequences of length 100 bp, as calculated by MoSBAT^31^. The red line across the dendrogram indicates the clustering threshold (see Fig. 3, **Methods**) determined by the optimal silhouette value^84^. PWMs and cluster membership are in **Supplementary Table 10**. PWM: position weight matrix, TF: transcription factor

**Extended Data Figure 7.**
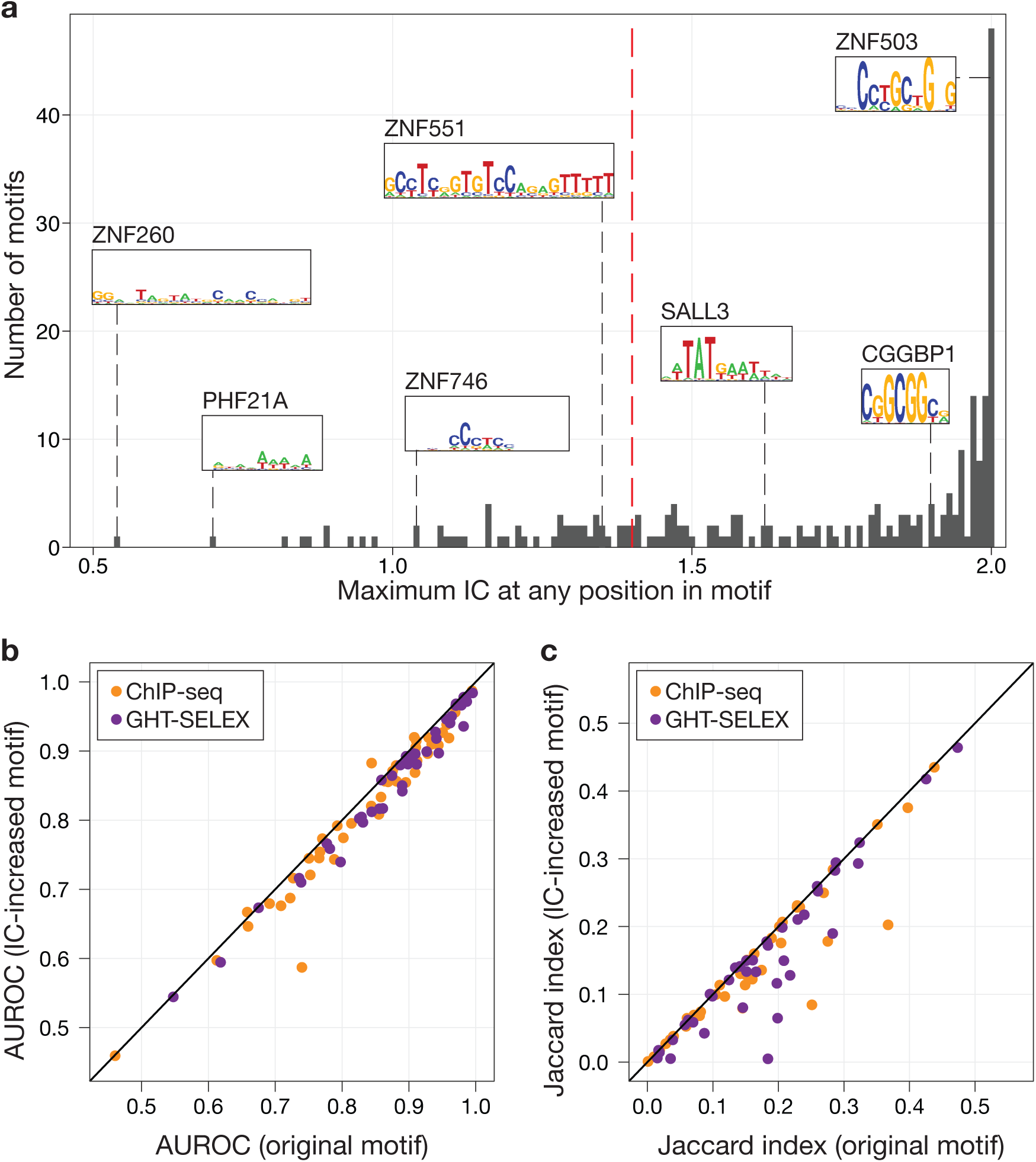
Motif degeneracy analysis. a, Histogram displays the maximum IC for any position within the representative PWM for all Codebook and control TFs. Logos are shown for TFs at various maximum positional IC values, for illustration. Red dashed line indicates an IC of 1.4. **b,** and **c,** comparison of original PWMs to IC-increased PWMs for the 52 TF PWMs for which no base position exceeded an IC of 1.4. **b,** AUROC values for original vs. IC-increased PWMs, discriminating ChIP-seq or GHT-SELEX peaks vs. random genomic background loci. **c,** Maximum Jaccard index for ChIP-seq or GHT-SELEX peak sets; using the approach described for optimized TOPs in **Methods**, for original vs. IC-increased PWMs. PWM: position weight matrix, IC: information content, TOP: triple-overlap regions, AUROC: area under the receiver operating characteristic

**Extended Data Figure 8.**
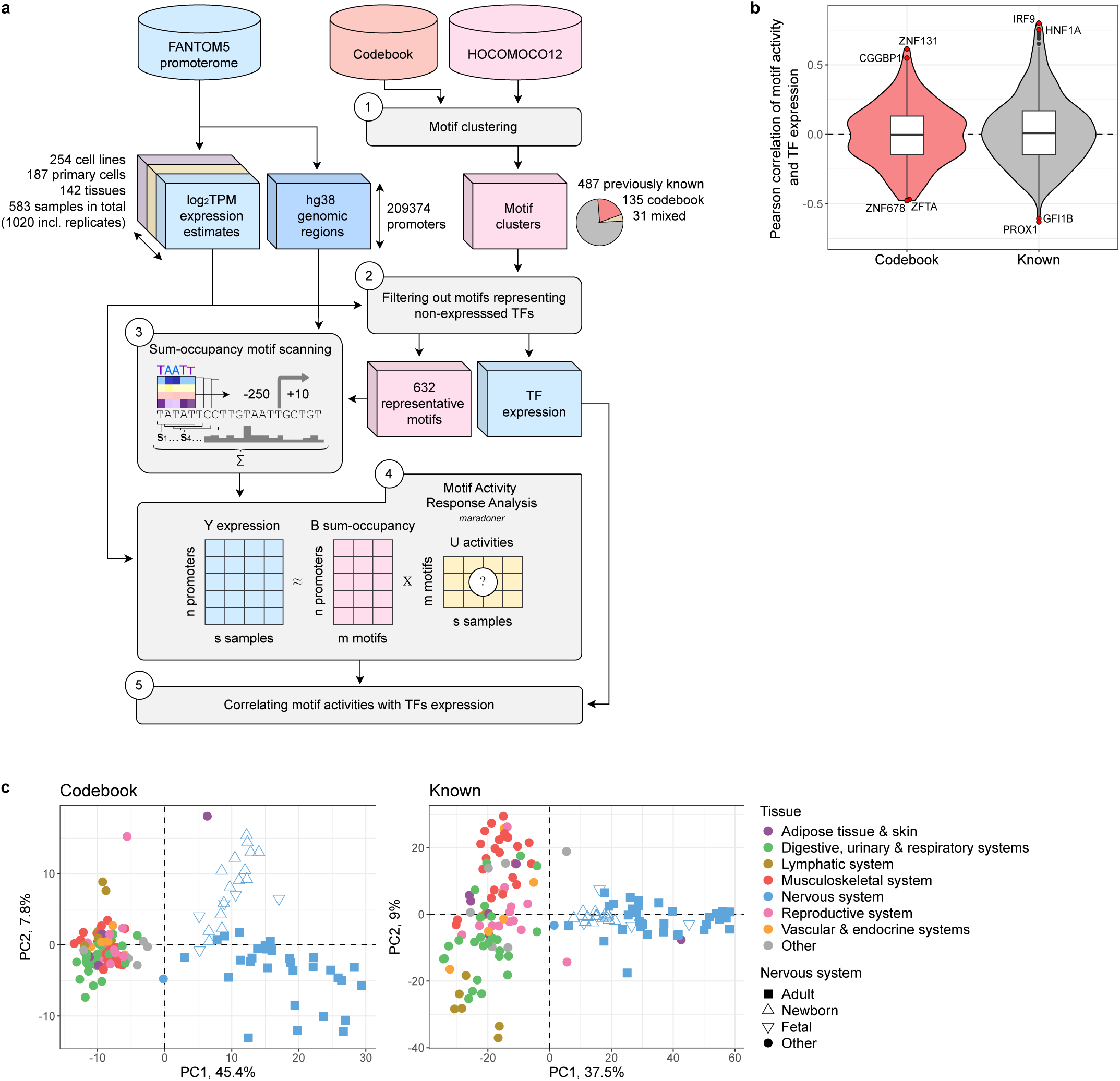
Motif activity response analysis highlights the relevance of Codebook motifs. a, MARA was conducted in five consecutive steps. (1) TF binding motifs from Codebook and HOCOMOCO^38^ (v12) were clustered, yielding 653 clusters, 135 of which consisted solely of Codebook motifs. (2) Motif clusters were filtered based on the total log_2_TPM of member TFs, resulting in 632 clusters of expressed TF motifs. (3) Representative motifs for each cluster were used to scan 209,374 genomic regions of 260 bp (250 bp upstream and 10 bp downstream relative to the representative TSS position of each promoter). (4) The sum-occupancy scores were used to perform the motif activity response analysis with the MARADONER Python package; (5) Pearson correlation coefficients were calculated for each representative motif by comparing the total TF expression against motif activities from MARA. **b,** Correlation between the TF expression (log_2_ TPM) and the activity of Codebook-specific (left) and known (right) motifs, the top 2 activators and inhibitors are labeled. Center line: median; box limits: upper and lower quartiles; whiskers: minimum and maximum values within the 1.5x interquartile range; dots: values outside the 1.5x interquartile range. **c,** Principal component analysis of FANTOM5 data for 142 human tissues using motif activities of 130 Codebook (left) and 471 known (right) representative motifs. X- and Y-axes: the first two principal components. Marker color: tissue groups, marker shape: nervous tissue development stage. TF: transcription factor, TPM: CAGE tags per million, CAGE: cap analysis of gene expression, TSS: transcription start site

**Extended Data Figure 9.**
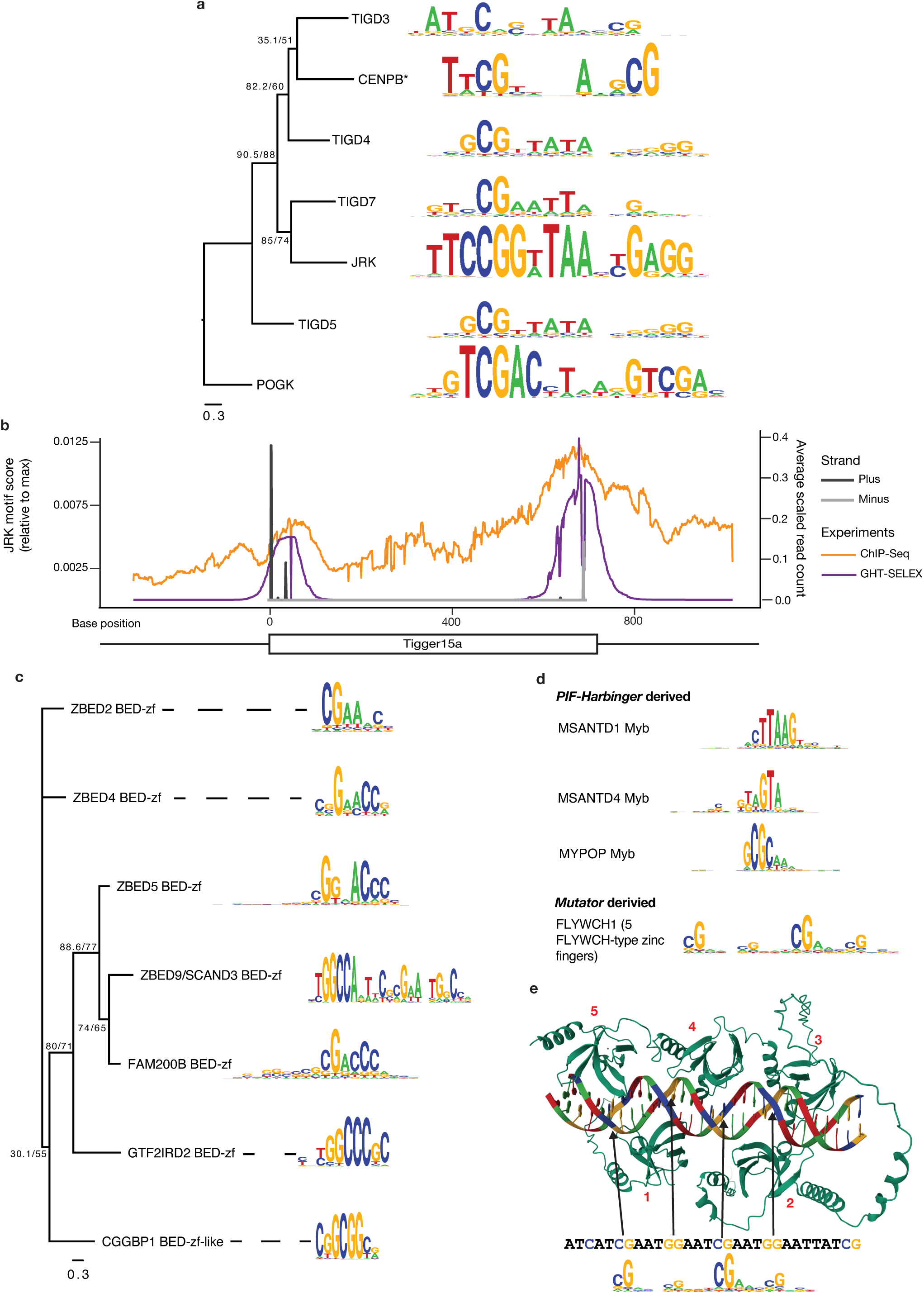
Transcription factors co-opted from DNA transposons. a, Motif logos of human TFs that are derived from the domestication of *Tigger* and *Pogo* DNA transposons and have previously known DNA binding motifs. Phylogenetic tree was derived from DBD sequence alignments produced with MAFFT L-INS-I^103^, and produced using the same method described in the **Extended Data** Fig. 2e legend. The tree is rooted on POGK, which is derived from a more distantly related family of *Pogo/Tigger*-like elements relative to the other proteins^104,105^. Sequence logos are Codebook-derived, except for CENPB, from the liturature^106^. **b**, average per-base read count over Tigger15a TOPs in the human genome, for JRK ChIP-seq (orange) and GHT-SELEX (purple), with peak sequences aligned to the Tigger15a consensus sequence. JRK PWM scores at each base of the Tigger15a consensus sequence are shown in black (plus strand) and grey (minus strand). **c,** Sequence logos of Codebook human TFs that are derived from the domestication of DBDs from *hAT* superfamily DNA transposons. Phylogenetic tree was derived from DBD sequence alignments produced with MAFFT L-INS-I^103^, although for ZBED4 the third of its four BED-zf domains was used. The tree was produced using the same method as described in the **Extended Data** Fig. 2e legend. **d,** Sequence logos of TFs derived from domestication of *Mutator* and *PIF-Harbinger* DNA transposons. **e**, Alphafold3 structure of the five FLYWCH-type zinc-fingers of FLYWCH1, and a TOP sequence from chr10:42167256-42167276 with 5bp flanks. Zinc finger domains are numbered according to their order in FLYWCH1 from the N-terminus to the C-terminus. Long linkers and terminal extensions are hidden for simplicity. DBD: DNA-binding domain, PWM: position weight matrix.

**Extended Data Figure 10.**
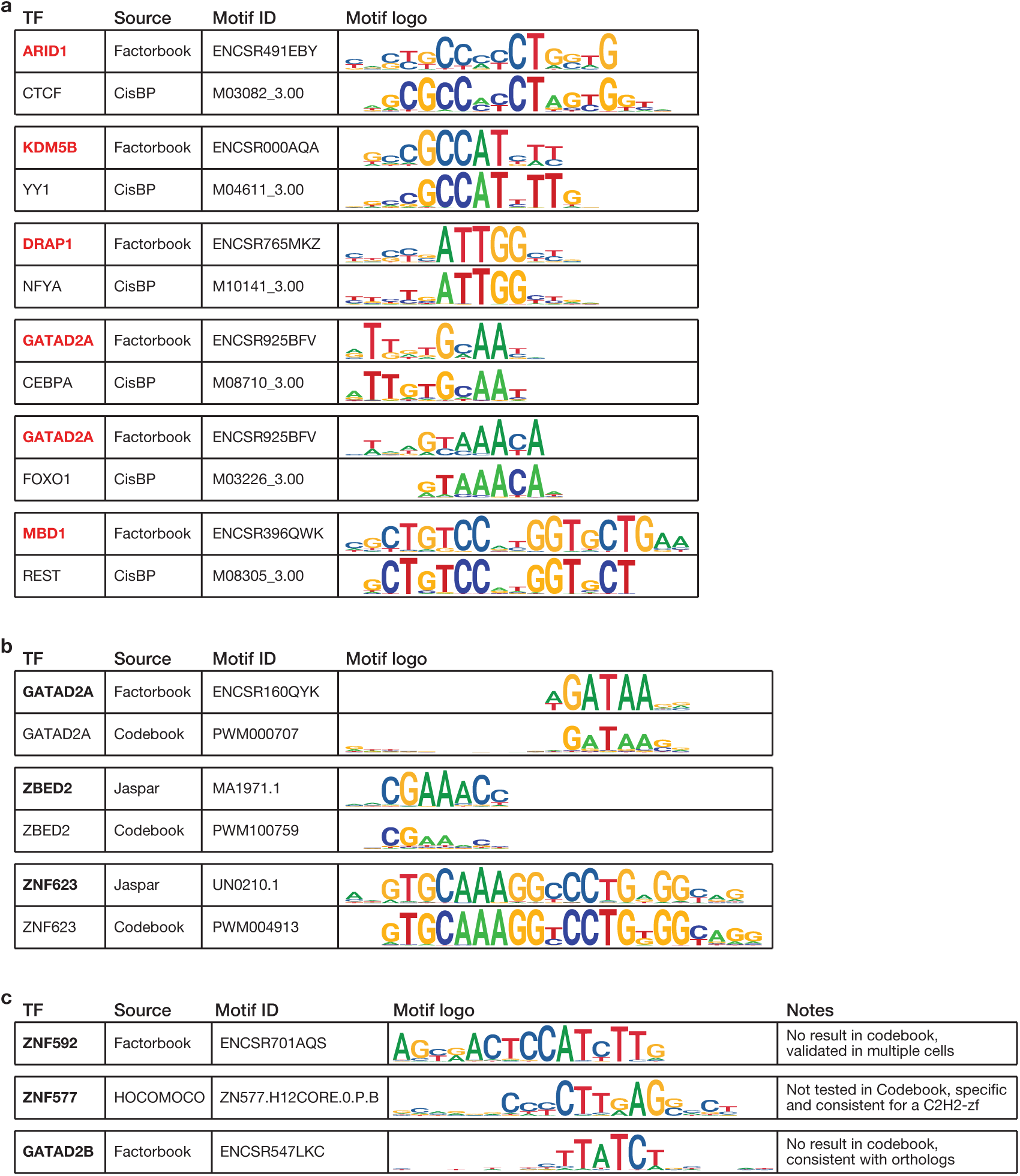
Examples of evaluation of external PWMs. a, Cases in which the external PWM matches the PWM of a well-studied TF that is a frequent “contaminant” motif in ChIP-seq^107^. In each example, the top sequence logo represents the external PWM, and the bottom sequence logo represents a highly similar CisBP PWM for a “contaminant” motif. **b,** Cases in which the external PWM (top in each example) is consistent with the Codebook PWM for the same TF (bottom in each example). **c,** External PWM sequence logos that cannot be explained as known contaminants or artifacts, some of which are supported by multiple lines of evidence and thus appear accurate. PWM: position weight matrix

## SUPPLEMENTARY INFORMATION

**Supplementary Table 1. List of tested proteins and assay successes.** Table lists the Codebook proteins and positive control TFs that were analyzed in the Codebook studies and provides metadata and information on whether they showed sequence-specific DNA binding activities in different types of experiments, together with the ID of the representative PWM selected in this study, if any.

**Supplementary Table 2. Inserts used in this study.** Table provides the amino acid sequence and type (full-length or DBD) for the 716 inserts used in the Codebook studies.

**Supplementary Table 3. Plasmids used in this study.** Table lists the plasmid backbone and insert for each of the 1,387 plasmids used in the Codebook studies.

**Supplementary Table 4. Experiments performed in this study.** Table lists the 4,805 experiments performed on Codebook and control TFs, along with 20 additional GFP control experiments, as well as 43 experiments from Schmitges et al. 2016^10^ and 25 experiments from Isakova et al. 2017^9^ that were reanalyzed. The experiment ID, experiment type, TF assayed, expert curation result, and plasmid ID are listed for each experiment. Each experiment is mapped to its ID in an accompanying manuscript^14^, and 9 additional experiments used only in an accompanying manuscript^14^ are explicitly marked.

**Supplementary Table 5. Representative PWMs.** Table displays logo representations for the PWMs that were manually selected as the representative for each of the TFs (i.e. the expert-curated motifs).

**Supplementary Table 6. Re-assessment of unsuccessful proteins.** Table summarizes the re-assessment outcomes for the unsuccessful proteins. A detailed description can be found in **Supplementary Document 1.**

**Supplementary Table 7. External ChIP-seq datasets**. Table lists external ChIP-seq datasets obtained from the GTRD database and ENCODE consortium for Codebook and control TFs as well as other putative TFs that were not tested in Codebook.

**Supplementary Table 8. TF-wise allele-specific binding sites.** This table contains transcription factor allele-specific binding sites (ASBs) identified using Codebook data at 5% FDR. Each entry represents an allele specific interaction between a TF and its DNA binding site in the vicinity of or directly encompassing the single-nucleotide variant, for which the allelic imbalance of read counts was observed between the reference and the alternative alleles.

**Supplementary Table 9. Aggregated allele-specific binding sites.** This table contains the joint set of allele-specific binding sites (FDR 5%) aggregated across all Codebook data (all transcription factors and experiment types), eQTL associated phenotypes (EBI GWAS Catalog), and known ASB annotations of the respective single-nucleotide variants. Each entry corresponds to the allele-specific binding of one or more TFs. The aggregated statistics combine evidence from multiple TFs and experiments (ChIP-Seq and multiple cycles of GHT-SELEX).

**Supplementary Table 10. Representative motifs for all human TFs.** Table lists representative motifs for Codebook TFs, TFs with previously known specificities, and motifs for 33 uncharacterized TFs curated from other publications (see “**Curation of new external motifs for previously uncharacterized TFs**”). Motif cluster membership is also indicated (**Extended Data Fig. 6**). The PWMs can be found in **Supplementary Data 1**.

**Supplementary Table 11. MARA motif clusters and representative motifs.** Details for the 653 motif clusters generated for the motif activity response analysis using FANTOM5 data.

**Supplementary Table 12. MARA results.** Table contains the MARA-predicted motif activity estimates across FANTOM5 samples for each motif cluster, their Pearson correlations with the expression of corresponding TFs in 583 samples used for the MARA analysis, ANOVA statistics, and significance estimates.

**Supplementary Table 13. External PWMs**. Table lists PWM identifiers, manual curation and other metadata for external motifs available from the JASPAR, HOCOMOCO and Factorbook databases.

**Supplementary Table 14. Updated census of human transcription factors and their motif coverage**. Table is modified from Lambert et al. 2018^1^ to display an updated motif coverage of human TFs and indicates the IDs of the representative motifs (see **Supplementary Table 10**).

**Supplementary Table 15. Motif derivation tools.** The table describes which methods were used for analyzing the main experiment types and summarizes the numbers of TFs and experiments that the methods were applied to, as well as the outcomes of motif analyses.

**Supplementary Table 16. Common artifact motifs.** Table contains PWMs that represent artifact or false positive signals that were commonly observed in the Codebook experiments.

**Supplementary Data 1. Representative motifs for all Human TFs.** The archive contains representative PWMs for all 1,421 TFs with a known binding specificity. Motif annotation is provided in **Supplementary Table 10**.

**Supplementary Document 1. Case-by-case re-assessment of unsuccessful Codebook proteins.** This document contains detailed case-by-case manual assessments for the proteins that did not yield successful experiments. The proteins are organized by DNA binding domain composition and the result of the re-assessment. The objective of the re-assessment was to determine whether manual examination of existing knowledge, and in some cases the Codebook data itself, suggest that they are more likely than not to be potential sequence specific TFs (i.e., false negatives in the scope of the Codebook project), or whether the available evidence instead supports them being true negatives..

**Supplementary Discussion.** Additional discussion of undercharacterized DNA-binding domains (particularly the SKI/SNO/DAC and C-Clamp domains), potential reasons for the failure of experiments for some proteins to yield motifs, and transposon-derived TFs.

**Supplementary Methods.** Expanded methods sections related to the allele-specific binding (ASB) analysis and Motif Activity Response Analysis (MARA).

## Notes

https://codebook.ccbr.utoronto.ca/

### Competing Interest Statement

Oriol Fornes is employed by Roche.

### Summary of Updates

Author list updated; Text significantly revised; Supplemental files updated and new supplemental files added; New figure (Figure 2) shows motif enrichment around ChIP-seq peaks for the 179 TFs with successful Codebook ChIP-seq experiments; New analysis shows the number of unique human TF motifs and the contribution of Codebook motifs to the motif repertoire; New section ("Codebook TF motifs predict gene expression levels across tissues and cell types") and figure (Figure 5) highlight apparent regulatory functions of novel TF motifs in regulating gene expression in specific biological contexts; New supplement (Supplementary Document 1) outlines reassessment of the 175 putative TFs that did not produce a motif, and identification of 83 proteins that are not likely to be TFs.

