## Supplementary Table 5 for "Codebook: sequence specificity and genomic binding of poorly-characterized human transcription factors"

| TF | Type | DBD type(s) | Motif source | Motif derivation algorithm | Motif |
|---|---|---|---|---|---|
| YY1 | Control | C2H2 ZF | ChIP-seq | MEME | 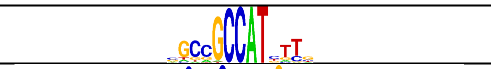 |
| CTCF | Control | C2H2 ZF | GHT-SELEX (Lysate) | Homer | 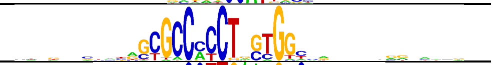 |
| GLI4 | Control | C2H2 ZF | GHT-SELEX (Lysate) | ExplaiNN | 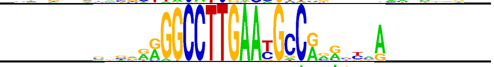 |
| ZFP3 | Control | C2H2 ZF | ChIP-seq | RCade | 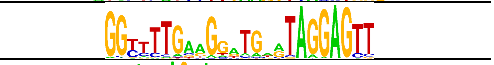 |
| ZIM3 | Control | C2H2 ZF | ChIP-seq | RCade | 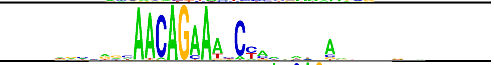 |
| ZNF134 | Control | C2H2 ZF | GHT-SELEX (Lysate) | RCade | 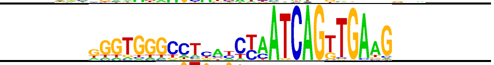 |
| ZNF18 | Control | C2H2 ZF | HT-SELEX (Lysate) | Autoseed | 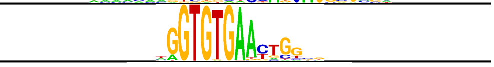 |
| ZNF260 | Control | C2H2 ZF | GHT-SELEX (Lysate) | Dimont | 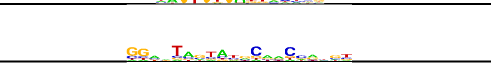 |
| ZNF322 | Control | C2H2 ZF | ChIP-seq | ChIPMunk | 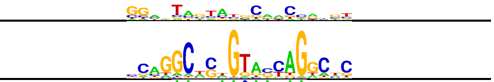 |
| ZNF35 | Control | C2H2 ZF | GHT-SELEX (Lysate) | RCade | 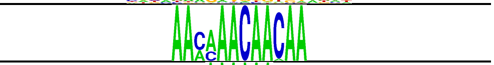 |
| ZNF384 | Control | C2H2 ZF | ChIP-seq | RCade | 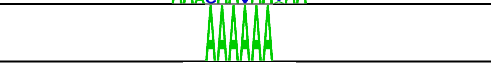 |
| ZNF770 | Control | C2H2 ZF | HT-SELEX (Lysate) | Dimont | 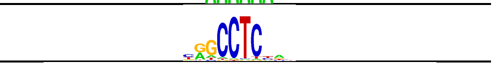 |
| ZNF250 | Control | C2H2 ZF | ChIP-seq | GkmSVM | 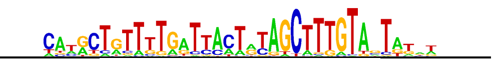 |
| ZNF264 | Control | C2H2 ZF | GHT-SELEX (Lysate) | ChIPMunk | 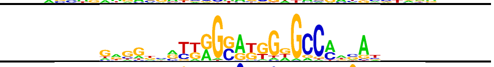 |
| ZNF436 | Control | C2H2 ZF | GHT-SELEX (Lysate) | RCade | 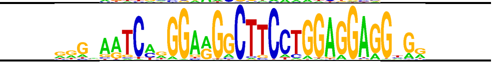 |
| ZNF596 | Control | C2H2 ZF | ChIP-seq | ChIPMunk | 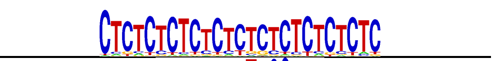 |
| ZNF8 | Control | C2H2 ZF | ChIP-seq | MEME | 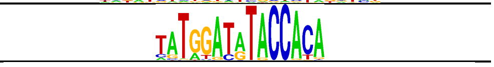 |
| ZFP28 | Control | C2H2 ZF | ChIP-seq | RCade | 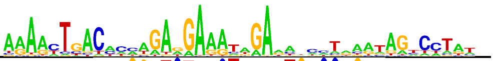 |
| ZNF121 | Control | C2H2 ZF | ChIP-seq | ExplaiNN | 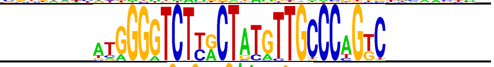 |
| ZNF140 | Control | C2H2 ZF | ChIP-seq | Streme | 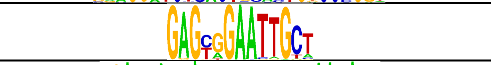 |
| ZNF146 | Control | C2H2 ZF | ChIP-seq | ChIPMunk | 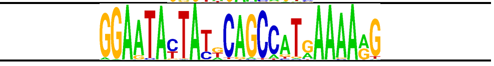 |
| ZNF30 | Control | C2H2 ZF | ChIP-seq | ChIPMunk | 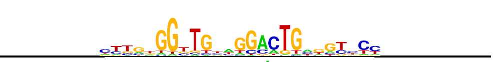 |
| ZNF317 | Control | C2H2 ZF | ChIP-seq | RCade | 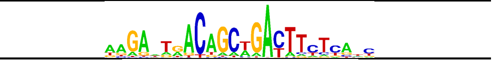 |
| ZNF382 | Control | C2H2 ZF | ChIP-seq | ChIPMunk | 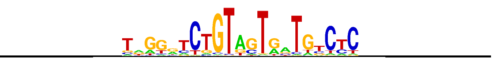 |
| ZNF454 | Control | C2H2 ZF | ChIP-seq | ExplaiNN | 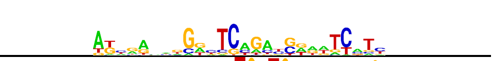 |
| ZNF490 | Control | C2H2 ZF | ChIP-seq | ChIPMunk | 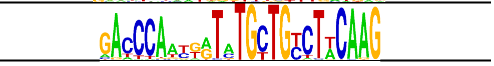 |
| ZNF582 | Control | C2H2 ZF | ChIP-seq | ChIPMunk | 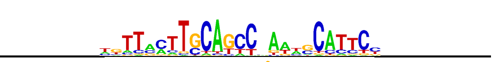 |
| ZNF708 | Control | C2H2 ZF | ChIP-seq | RCade | 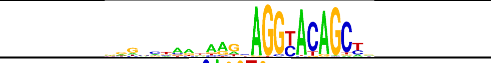 |
| SNAI1 | Control | C2H2 ZF | GHT-SELEX (IVT) | MEME | 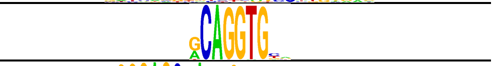 |
| ZNF16 | Control | C2H2 ZF | GHT-SELEX (Lysate) | ChIPMunk | 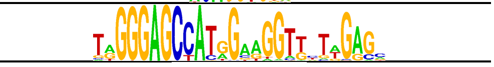 |
| ZSCAN4 | Control | C2H2 ZF | GHT-SELEX (IVT) | ProBound | 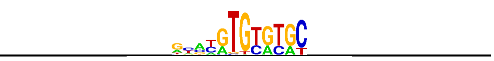 |
| MAX | Control | bHLH | GHT-SELEX (Lysate) | Dimont | 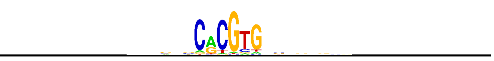 |

| TF | Type | DBD type(s) | Motif source | Motif derivation algorithm | Motif |
|---|---|---|---|---|---|
| FOSL2 | Control | bZIP | ChIP-seq | ChIPMunk | 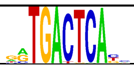 |
| GCM1 | Control | GCM | GHT-SELEX (IVT) | Dimont | 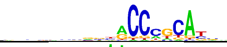 |
| LEF1 | Control | HMG/Sox | HT-SELEX (Lysate) | Autoseed | 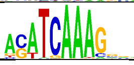 |
| SOX2 | Control | HMG/Sox | GHT-SELEX (Lysate) | Dimont | 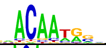 |
| SRY | Control | HMG/Sox | HT-SELEX (Lysate) | Autoseed | 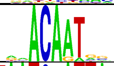 |
| PAX7 | Control | Homeodomain; Paired box | HT-SELEX (Lysate) | Streme | 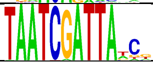 |
| RORB | Control | Nuclear receptor | GHT-SELEX (Lysate) | Dimont | 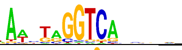 |
| VDR | Control | Nuclear receptor | HT-SELEX (IVT) | Autoseed | 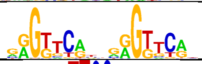 |
| ELF3 | Control | Ets; AT hook | ChIP-seq | MEME | 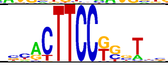 |
| NFKB1 | Control | Rel | GHT-SELEX (Lysate) | Dimont | 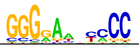 |
| MYF6 | Control | bHLH | GHT-SELEX (Lysate) | Dimont | 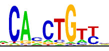 |
| FLI1 | Control | Ets | GHT-SELEX (IVT) | Dimont | 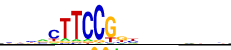 |
| GABPA | Control | Ets | HT-SELEX (Lysate) | Autoseed | 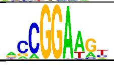 |
| SOX15 | Control | HMG/Sox | GHT-SELEX (Lysate) | Dimont | 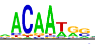 |
| NR1H4 | Control | Nuclear receptor | HT-SELEX (Lysate) | Autoseed | 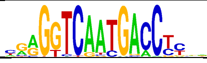 |
| RFX5 | Control | RFX | ChIP-seq | ChIPMunk | 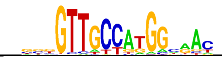 |
| MGA | Control | T | GHT-SELEX (Lysate) | Dimont | 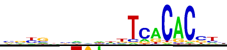 |
| JUN | Control | bZIP | ChIP-seq | Streme | 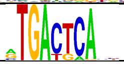 |
| LHX6 | Control | Homeodomain | ChIP-seq | Dimont | 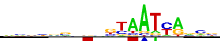 |
| JUNB | Control | bZIP | ChIP-seq | MEME | 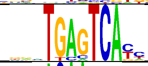 |
| SOX5 | Control | HMG/Sox | HT-SELEX (IVT) | ChIPMunk | 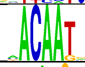 |
| NR4A2 | Control | Nuclear receptor | HT-SELEX (IVT) | ChIPMunk | 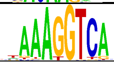 |
| HMGA1 | Control | AT hook | HT-SELEX (GFPIVT) | Dimont | 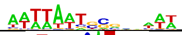 |
| POU5F1 | Control | Homeodomain; POU | GHT-SELEX (GFPIVT) | MEME | 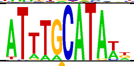 |
| RARA | Control | Nuclear receptor | HT-SELEX (IVT) | ProBound | 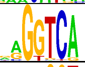 |
| RXRA | Control | Nuclear receptor | HT-SELEX (Lysate) | MEME | 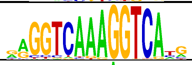 |
| FIZ1 | Codebook TF | C2H2 ZF | HT-SELEX (Lysate) | Autoseed | 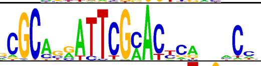 |
| PRDM10 | Codebook TF | C2H2 ZF | HT-SELEX (GFPIVT) | Autoseed | 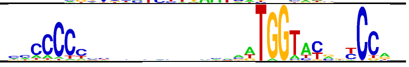 |
| PRDM13 | Codebook TF | C2H2 ZF | GHT-SELEX (Lysate) | Dimont | 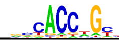 |
| PRDM5 | Codebook TF | C2H2 ZF | HT-SELEX (GFPIVT) | Homer | 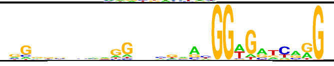 |
| RLF | Codebook TF | C2H2 ZF | ChIP-seq | Dimont | 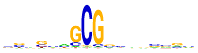 |
| ZBTB8A | Codebook TF | C2H2 ZF | HT-SELEX (Lysate) | MEME | 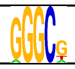 |

| TF | Type | DBD type(s) | Motif source | Motif derivation algorithm | Motif |
|---|---|---|---|---|---|
| ZBTB8B | Codebook TF | C2H2 ZF | HT-SELEX (Lysate) | Homer | 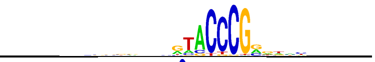 |
| ZFAT | Codebook TF | C2H2 ZF | GHT-SELEX (GFPIVT) | ChIPMunk | 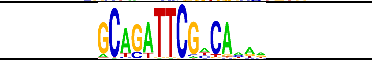 |
| ZNF107 | Codebook TF | C2H2 ZF | ChIP-seq | ChIPMunk | 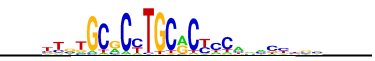 |
| ZNF131 | Codebook TF | C2H2 ZF | GHT-SELEX (Lysate) | Dimont | 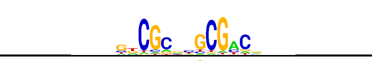 |
| ZNF142 | Codebook TF | C2H2 ZF | ChIP-seq | ChIPMunk | 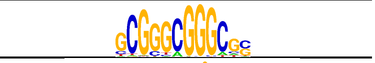 |
| ZNF226 | Codebook TF | C2H2 ZF | HT-SELEX (GFPIVT) | ChIPMunk | 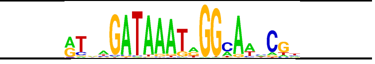 |
| ZNF251 | Codebook TF | C2H2 ZF | ChIP-seq | Dimont | 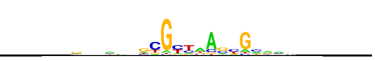 |
| ZNF347 | Codebook TF | C2H2 ZF | ChIP-seq | ChIPMunk | 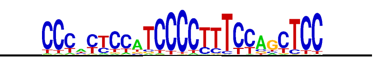 |
| ZNF362 | Codebook TF | C2H2 ZF | SMiLE-seq | Dimont | 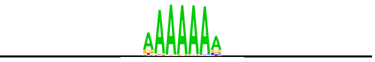 |
| ZNF367 | Codebook TF | C2H2 ZF | GHT-SELEX (GFPIVT) | Dimont | 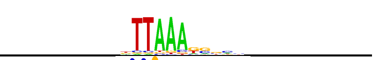 |
| ZNF395 | Codebook TF | C2H2 ZF | GHT-SELEX (IVT) | ProBound | 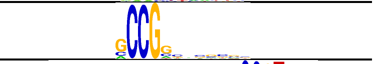 |
| ZNF407 | Codebook TF | C2H2 ZF | GHT-SELEX (Lysate) | ChIPMunk | 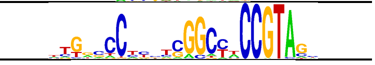 |
| ZNF43 | Codebook TF | C2H2 ZF | ChIP-seq | Dimont | 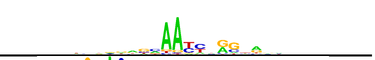 |
| ZNF493 | Codebook TF | C2H2 ZF | HT-SELEX (GFPIVT) | ChIPMunk | 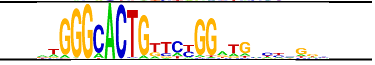 |
| ZNF500 | Codebook TF | C2H2 ZF | GHT-SELEX (GFPIVT) | Dimont | 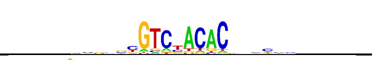 |
| ZNF648 | Codebook TF | C2H2 ZF | GHT-SELEX (GFPIVT) | Homer | 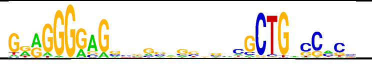 |
| ZNF726 | Codebook TF | C2H2 ZF | ChIP-seq | Dimont | 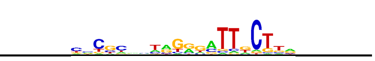 |
| ZNF780B | Codebook TF | C2H2 ZF | GHT-SELEX (GFPIVT) | Dimont | 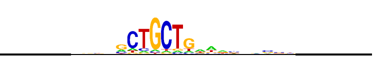 |
| ZNF831 | Codebook TF | C2H2 ZF | GHT-SELEX (GFPIVT) | Dimont | 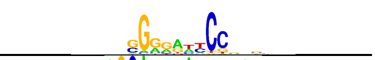 |
| ZNF841 | Codebook TF | C2H2 ZF | GHT-SELEX (GFPIVT) | Streme | 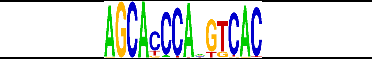 |
| ZNF518B | Codebook TF | C2H2 ZF | GHT-SELEX (GFPIVT) | Dimont | 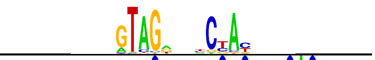 |
| ZNF865 | Codebook TF | C2H2 ZF | ChIP-seq | ExplaiNN | 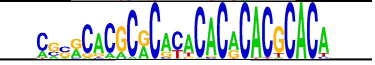 |
| SALL3 | Codebook TF | C2H2 ZF | HT-SELEX (Lysate) | ProBound | 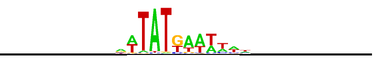 |
| SLC2A4RG | Codebook TF | C2H2 ZF | GHT-SELEX (IVT) | Autoseed | 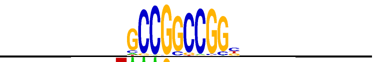 |
| ZBTB40 | Codebook TF | C2H2 ZF | HT-SELEX (GFPIVT) | Autoseed | 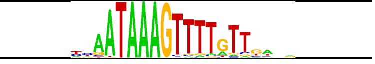 |
| ZBTB41 | Codebook TF | C2H2 ZF | GHT-SELEX (GFPIVT) | Dimont | 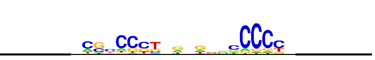 |
| ZBTB47 | Codebook TF | C2H2 ZF | GHT-SELEX (GFPIVT) | Dimont | 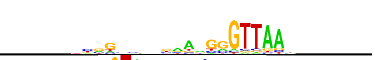 |
| ZNF20 | Codebook TF | C2H2 ZF | GHT-SELEX (GFPIVT) | Streme | 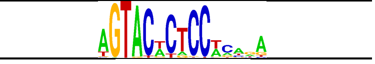 |
| ZNF215 | Codebook TF | C2H2 ZF | HT-SELEX (GFPIVT) | ProBound | 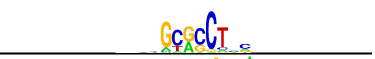 |
| ZNF234 | Codebook TF | C2H2 ZF | HT-SELEX (GFPIVT) | Autoseed | 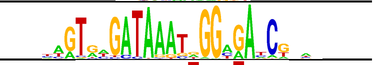 |
| ZNF286B | Codebook TF | C2H2 ZF | HT-SELEX (GFPIVT) | Autoseed | 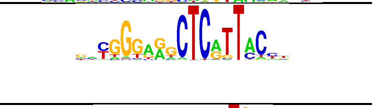 |
| ZNF292 | Codebook TF | C2H2 ZF | HT-SELEX (GFPIVT) | MEME | 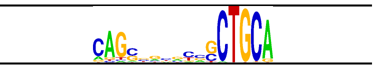 |

| TF | Type | DBD type(s) | Motif source | Motif derivation algorithm | Motif |
|---|---|---|---|---|---|
| ZNF335 | Codebook TF | C2H2 ZF | HT-SELEX (GFPIVT) | Autoseed | 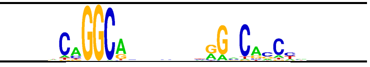 |
| ZNF470 | Codebook TF | C2H2 ZF | ChIP-seq | Dimont | 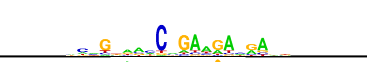 |
| ZNF471 | Codebook TF | C2H2 ZF | GHT-SELEX (GFPIVT) | MEME | 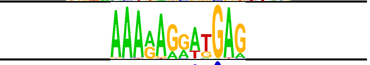 |
| ZNF48 | Codebook TF | C2H2 ZF | GHT-SELEX (GFPIVT) | ProBound | 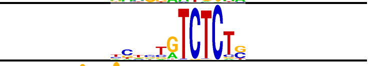 |
| ZNF510 | Codebook TF | C2H2 ZF | HT-SELEX (GFPIVT) | Autoseed | 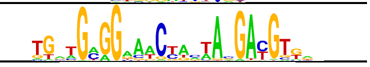 |
| ZNF551 | Codebook TF | C2H2 ZF | ChIP-seq | ChIPMunk | 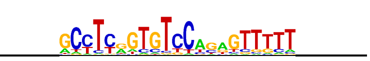 |
| ZNF57 | Codebook TF | C2H2 ZF | ChIP-seq | ChIPMunk | 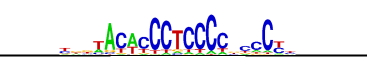 |
| ZNF575 | Codebook TF | C2H2 ZF | GHT-SELEX (GFPIVT) | ProBound | 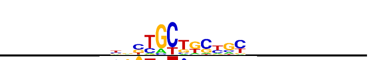 |
| ZNF606 | Codebook TF | C2H2 ZF | GHT-SELEX (GFPIVT) | Streme | 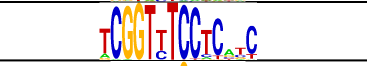 |
| ZNF646 | Codebook TF | C2H2 ZF | GHT-SELEX (IVT) | ProBound | 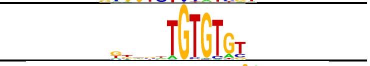 |
| ZNF665 | Codebook TF | C2H2 ZF | HT-SELEX (GFPIVT) | Autoseed | 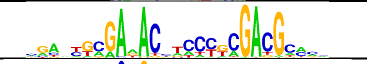 |
| ZNF672 | Codebook TF | C2H2 ZF | HT-SELEX (GFPIVT) | Autoseed | 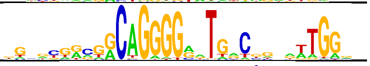 |
| ZNF676 | Codebook TF | C2H2 ZF | HT-SELEX (GFPIVT) | Autoseed | 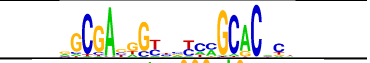 |
| ZNF678 | Codebook TF | C2H2 ZF | GHT-SELEX (GFPIVT) | ExplaiNN | 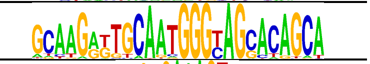 |
| ZNF683 | Codebook TF | C2H2 ZF | HT-SELEX (GFPIVT) | Streme | 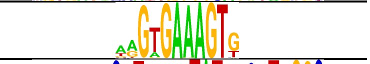 |
| ZNF689 | Codebook TF | C2H2 ZF | GHT-SELEX (GFPIVT) | ExplaiNN | 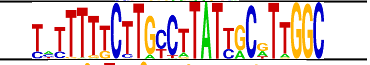 |
| ZNF696 | Codebook TF | C2H2 ZF | HT-SELEX (GFPIVT) | ChIPMunk | 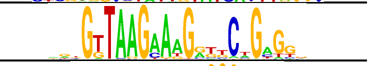 |
| ZNF699 | Codebook TF | C2H2 ZF | HT-SELEX (GFPIVT) | Autoseed | 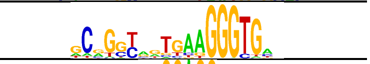 |
| ZNF70 | Codebook TF | C2H2 ZF | HT-SELEX (GFPIVT) | MEME | 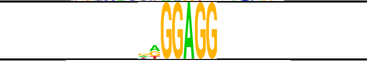 |
| ZNF700 | Codebook TF | C2H2 ZF | ChIP-seq | Dimont | 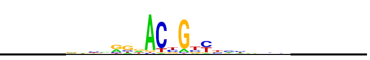 |
| ZNF721 | Codebook TF | C2H2 ZF | GHT-SELEX (GFPIVT) | Dimont | 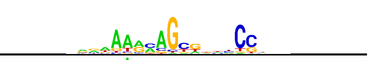 |
| ZNF724 | Codebook TF | C2H2 ZF | HT-SELEX (GFPIVT) | Autoseed | 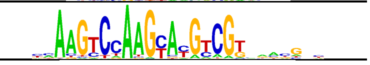 |
| ZNF728 | Codebook TF | C2H2 ZF | GHT-SELEX (GFPIVT) | ChIPMunk | 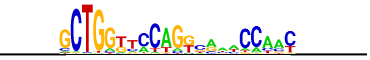 |
| ZNF732 | Codebook TF | C2H2 ZF | GHT-SELEX (GFPIVT) | Dimont | 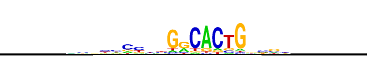 |
| ZNF746 | Codebook TF | C2H2 ZF | HT-SELEX (GFPIVT) | Dimont | 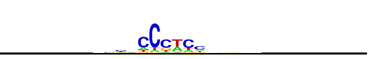 |
| ZNF775 | Codebook TF | C2H2 ZF | GHT-SELEX (GFPIVT) | Dimont | 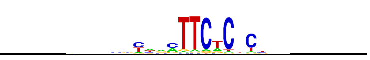 |
| ZNF800 | Codebook TF | C2H2 ZF | GHT-SELEX (GFPIVT) | Dimont | 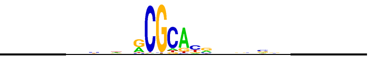 |
| ZNF814 | Codebook TF | C2H2 ZF | GHT-SELEX (GFPIVT) | Dimont | 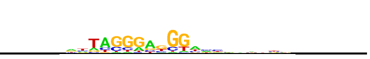 |
| ZNF836 | Codebook TF | C2H2 ZF | ChIP-seq | ChIPMunk | 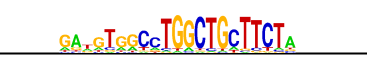 |
| ZNF845 | Codebook TF | C2H2 ZF | ChIP-seq | Dimont | 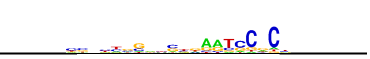 |
| ZNF850 | Codebook TF | C2H2 ZF | ChIP-seq | ChIPMunk | 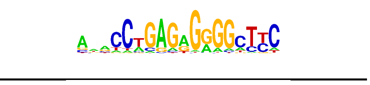 |
| ZSCAN25 | Codebook TF | C2H2 ZF | GHT-SELEX (Lysate) | Dimont | 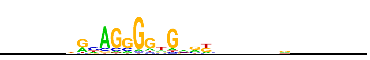 |

| TF | Type | DBD type(s) | Motif source | Motif derivation algorithm | Motif |
|---|---|---|---|---|---|
| ZNF233 | Codebook TF | C2H2 ZF | ChIP-seq | Dimont | 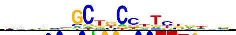 |
| ZNF623 | Codebook TF | C2H2 ZF | GHT-SELEX (GFPIVT) | ChIPMunk | 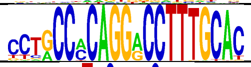 |
| ZNF66 | Codebook TF | C2H2 ZF | GHT-SELEX (GFPIVT) | Homer | 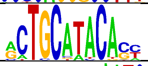 |
| ZNF772 | Codebook TF | C2H2 ZF | GHT-SELEX (GFPIVT) | ChIPMunk | 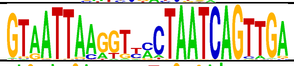 |
| ZNF773 | Codebook TF | C2H2 ZF | GHT-SELEX (GFPIVT) | ExplaiNN | 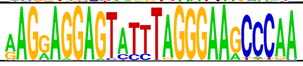 |
| ZSCAN2 | Codebook TF | C2H2 ZF | ChIP-seq | RCade | 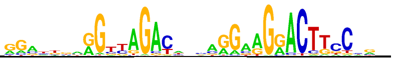 |
| ZXDA | Codebook TF | C2H2 ZF | ChIP-seq | ChIPMunk | 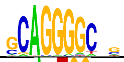 |
| ZNF516 | Codebook TF | C2H2 ZF | HT-SELEX (GFPIVT) | MEME | 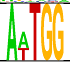 |
| ZNF878 | Codebook TF | C2H2 ZF | ChIP-seq | Dimont | 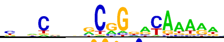 |
| ZXDB | Codebook TF | C2H2 ZF | ChIP-seq | RCade | 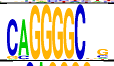 |
| ZXDC | Codebook TF | C2H2 ZF | ChIP-seq | Homer | 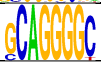 |
| ZNF14 | Codebook TF | C2H2 ZF | ChIP-seq | ChIPMunk | 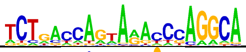 |
| ZNF503 | Codebook TF | C2H2 ZF | ChIP-seq | MEME | 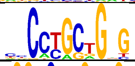 |
| ZNF578 | Codebook TF | C2H2 ZF | ChIP-seq | Homer | 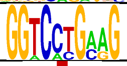 |
| ZNF703 | Codebook TF | C2H2 ZF | ChIP-seq | ChIPMunk | 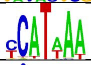 |
| ZNF705E | Codebook TF | C2H2 ZF | ChIP-seq | ChIPMunk | 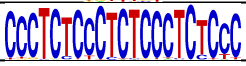 |
| ZNF709 | Codebook TF | C2H2 ZF | ChIP-seq | ChIPMunk | 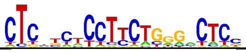 |
| ZNF888 | Codebook TF | C2H2 ZF | ChIP-seq | Dimont | 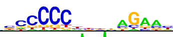 |
| ZNF92 | Codebook TF | C2H2 ZF | ChIP-seq | ChIPMunk | 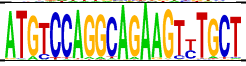 |
| ZSCAN12 | Codebook TF | C2H2 ZF | ChIP-seq | ChIPMunk | 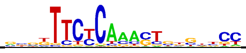 |
| AC092835 | Codebook TF | C2H2 ZF | ChIP-seq | ChIPMunk | 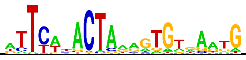 |
| ATMIN | Codebook TF | C2H2 ZF | ChIP-seq | ExplaiNN | 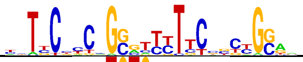 |
| ZNF208 | Codebook TF | C2H2 ZF | GHT-SELEX (GFPIVT) | Streme | 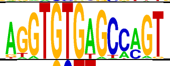 |
| ZNF853 | Codebook TF | C2H2 ZF | GHT-SELEX (GFPIVT) | ChIPMunk | 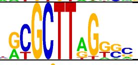 |
| ZNF729 | Codebook TF | C2H2 ZF | GHT-SELEX (GFPIVT) | ChIPMunk | 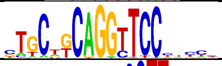 |
| ZNF813 | Codebook TF | C2H2 ZF | HT-SELEX (GFPIVT) | MEME | 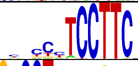 |
| ZBTB24 | Codebook TF | C2H2 ZF; AT hook | HT-SELEX (GFPIVT) | ChIPMunk | 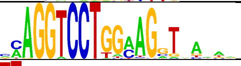 |
| CASZ1 | Codebook TF | C2H2 ZF | HT-SELEX (Lysate) | ChIPMunk | 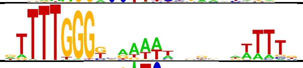 |
| ZBTB5 | Codebook TF | C2H2 ZF | SMiLE-seq | ChIPMunk | 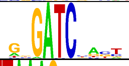 |
| ZFP91 | Codebook TF | C2H2 ZF | GHT-SELEX (GFPIVT) | Streme | 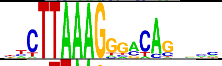 |
| ZNF536 | Codebook TF | C2H2 ZF | HT-SELEX (GFPIVT) | ChIPMunk | 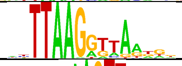 |
| MYT1 | Codebook TF | C2H2 ZF | HT-SELEX (Lysate) | MEME | 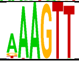 |

| TF | Type | DBD type(s) | Motif source | Motif derivation algorithm | Motif |
|---|---|---|---|---|---|
| TSHZ2 | Codebook TF | C2H2 ZF | HT-SELEX (IVT) | Dimont | 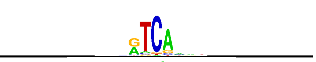 |
| ZKSCAN4 | Codebook TF | C2H2 ZF | GHT-SELEX (GFPIVT) | ChIPMunk | 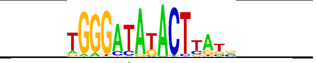 |
| ZNF160 | Codebook TF | C2H2 ZF | GHT-SELEX (GFPIVT) | ChIPMunk | 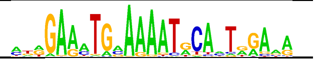 |
| ZNF229 | Codebook TF | C2H2 ZF | HT-SELEX (GFPIVT) | Autoseed | 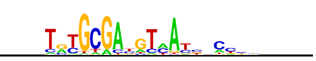 |
| ZNF275 | Codebook TF | C2H2 ZF | HT-SELEX (GFPIVT) | Autoseed | 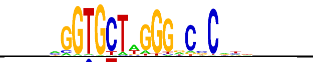 |
| ZNF358 | Codebook TF | C2H2 ZF | HT-SELEX (GFPIVT) | ChIPMunk | 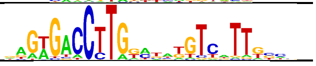 |
| ZNF497 | Codebook TF | C2H2 ZF | GHT-SELEX (GFPIVT) | ChIPMunk | 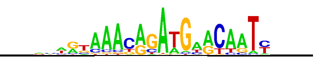 |
| ZNF532 | Codebook TF | C2H2 ZF | HT-SELEX (GFPIVT) | Dimont | 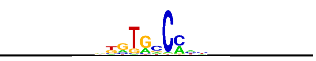 |
| ZNF568 | Codebook TF | C2H2 ZF | GHT-SELEX (GFPIVT) | Dimont | 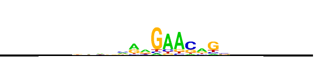 |
| ZNF569 | Codebook TF | C2H2 ZF | GHT-SELEX (GFPIVT) | Dimont | 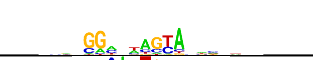 |
| ZNF587B | Codebook TF | C2H2 ZF | HT-SELEX (GFPIVT) | ChIPMunk | 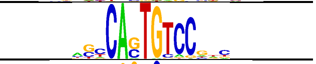 |
| ZNF668 | Codebook TF | C2H2 ZF | GHT-SELEX (GFPIVT) | ChIPMunk | 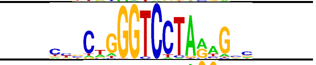 |
| ZNF827 | Codebook TF | C2H2 ZF | HT-SELEX (GFPIVT) | ProBound | 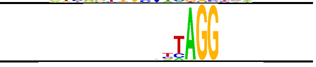 |
| ZNF618 | Codebook TF | C2H2 ZF | GHT-SELEX (Lysate) | ChIPMunk | 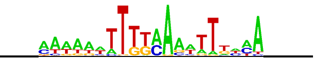 |
| ZNF83 | Codebook TF | C2H2 ZF | GHT-SELEX (GFPIVT) | Dimont | 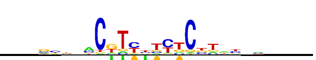 |
| ZNF526 | Codebook TF | C2H2 ZF | HT-SELEX (GFPIVT) | ChIPMunk | 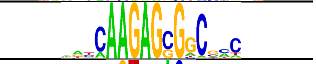 |
| ZNF788P | Codebook TF | C2H2 ZF | HT-SELEX (GFPIVT) | Streme | 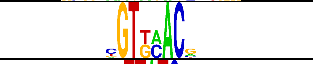 |
| ZFPM1 | Codebook TF | C2H2 ZF | SMiLE-seq | ChIPMunk | 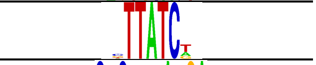 |
| ZNF507 | Codebook TF | C2H2 ZF | SMiLE-seq | Streme | 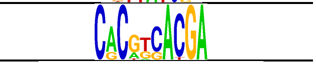 |
| FAM200B | Codebook TF | BED ZF | ChIP-seq | Dimont | 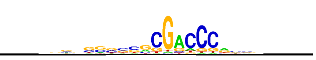 |
| ZBED2 | Codebook TF | BED ZF | GHT-SELEX (Lysate) | Dimont | 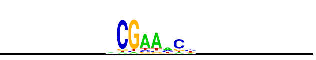 |
| ZBED5 | Codebook TF | BED ZF | GHT-SELEX (GFPIVT) | Dimont | 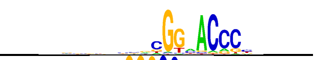 |
| ZFTA | Codebook TF | BED ZF | ChIP-seq | MEME | 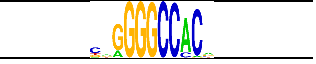 |
| USF3 | Codebook TF | bHLH | GHT-SELEX (Lysate) | Dimont | 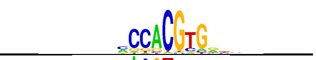 |
| CREB3L3 | Codebook TF | bZIP | GHT-SELEX (Lysate) | ChIPMunk | 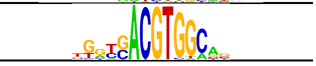 |
| TIGD3 | Codebook TF | CENPB | GHT-SELEX (Lysate) | Dimont | 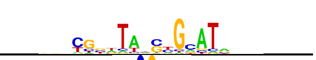 |
| CXXC4 | Codebook TF | CxxC | HT-SELEX (IVT) | Autoseed | 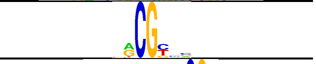 |
| FBXL19 | Codebook TF | CxxC | GHT-SELEX (Lysate) | ProBound | 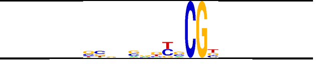 |
| TET3 | Codebook TF | CxxC | HT-SELEX (IVT) | Dimont | 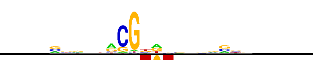 |
| MKX | Codebook TF | Homeodomain | HT-SELEX (GFPIVT) | MEME | 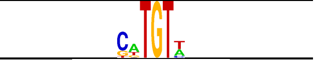 |
| TPRX1 | Codebook TF | Homeodomain | HT-SELEX (GFPIVT) | Dimont | 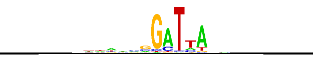 |
| MSANTD1 | Codebook TF | MADF | GHT-SELEX (Lysate) | Dimont | 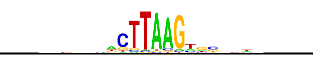 |

| TF | Type | DBD type(s) | Motif source | Motif derivation algorithm | Motif |
|---|---|---|---|---|---|
| DMTF1 | Codebook TF | Myb/SANT | GHT-SELEX (GFPIVT) | Dimont | 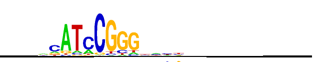 |
| SP140 | Codebook TF | SAND | PBM | Dimont | 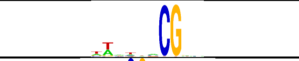 |
| SP140L | Codebook TF | SAND | SMiLE-seq | ProBound | 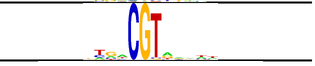 |
| TIGD4 | Codebook TF | CENPB | GHT-SELEX (Lysate) | Dimont | 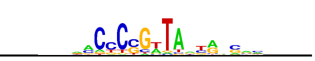 |
| CAMTA1 | Codebook TF | CG | GHT-SELEX (GFPIVT) | Dimont | 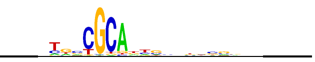 |
| CAMTA2 | Codebook TF | CG | GHT-SELEX (Lysate) | Dimont | 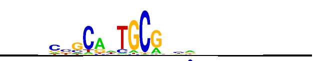 |
| SP100 | Codebook TF | SAND | GHT-SELEX (GFPIVT) | ProBound | 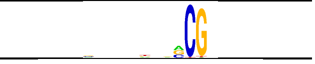 |
| MYPOP | Codebook TF | Myb/SANT | GHT-SELEX (Lysate) | Dimont | 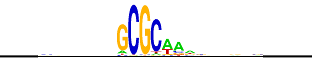 |
| TERF1 | Codebook TF | Myb/SANT | GHT-SELEX (Lysate) | Dimont | 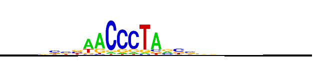 |
| DNTTIP1 | Codebook TF | AT hook | HT-SELEX (Lysate) | Dimont | 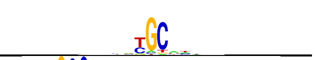 |
| ZBED9 | Codebook TF | BED ZF | HT-SELEX (Lysate) | ChIPMunk | 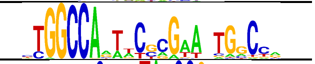 |
| JRK | Codebook TF | CENPB | HT-SELEX (GFPIVT) | ChIPMunk | 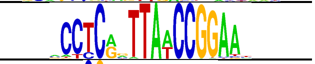 |
| KDM2A | Codebook TF | CxxC | GHT-SELEX (Lysate) | ProBound | 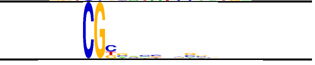 |
| FLYWCH1 | Codebook TF | FLYWCH | ChIP-seq | Dimont | 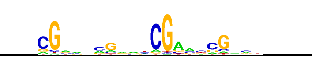 |
| LEUTX | Codebook TF | Homeodomain | GHT-SELEX (GFPIVT) | Dimont | 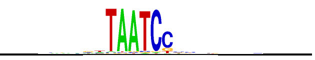 |
| BATF2 | Codebook TF | bZIP | HT-SELEX (Lysate) | ProBound | 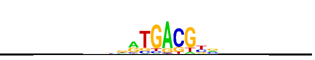 |
| MYRFL | Codebook TF | Ndt80/PhoG | ChIP-seq | ChIPMunk | 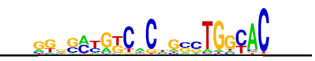 |
| MBD1 | Codebook TF | MBD; CxxC ZF | ChIP-seq | Dimont | 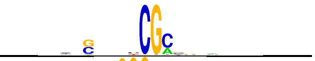 |
| GTF2IRD2 | Codebook TF | GTF2I | HT-SELEX (Lysate) | ChIPMunk | 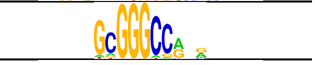 |
| BHLHA9 | Codebook TF | bHLH | ChIP-seq | Dimont | 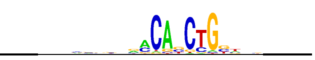 |
| MBD3 | Codebook TF | MBD | ChIP-seq | Dimont | 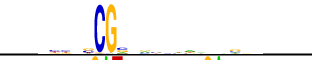 |
| MYSM1 | Codebook TF | Myb/SANT | ChIP-seq | Streme | 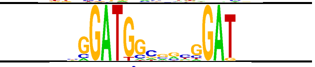 |
| TTF1 | Codebook TF | Myb/SANT | ChIP-seq | ChIPMunk | 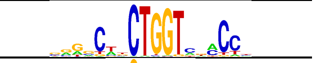 |
| ZBED4 | Codebook TF | BED ZF | SMiLE-seq | Homer | 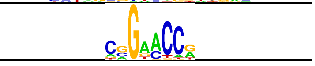 |
| TIGD7 | Codebook TF | CENPB | GHT-SELEX (GFPIVT) | Dimont | 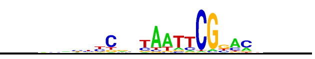 |
| MSANTD4 | Codebook TF | Myb/SANT | GHT-SELEX (Lysate) | Dimont | 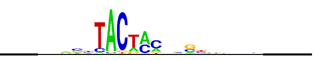 |
| TIGD5 | Codebook TF | CENPB | GHT-SELEX (GFPIVT) | Dimont | 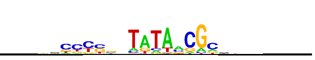 |
| ZGLP1 | Codebook TF | GATA | HT-SELEX (Lysate) | Homer | 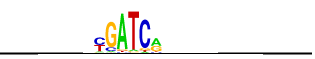 |
| GATAD2A | Codebook TF | GATA | GHT-SELEX (GFPIVT) | Dimont | 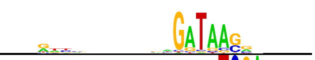 |
| POGK | Codebook TF | Brinker | HT-SELEX (IVT) | ChIPMunk | 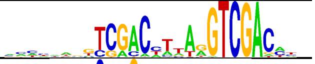 |
| GRHL3 | Codebook TF | Grainyhead | HT-SELEX (IVT) | ChIPMunk | 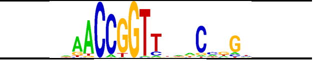 |
| HMG20A | Codebook TF | HMG/Sox | HT-SELEX (GFPIVT) | Streme | 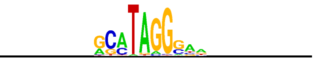 |

| TF | Type | DBD type(s) | Motif source | Motif derivation algorithm | Motif |
|---|---|---|---|---|---|
| AHCTF1 | Codebook TF | AT hook | PBM | Dimont | 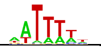 |
| GLYR1 | Codebook TF | AT hook | PBM | Dimont | 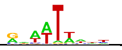 |
| PHF21A | Codebook TF | AT hook | PBM | Dimont | 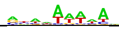 |
| SETBP1 | Codebook TF | AT hook | PBM | Dimont | 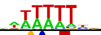 |
| MBNL2 | Codebook TF | CCCH ZF | PBM | ChIPMunk | 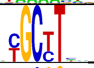 |
| CGGBP1 | Codebook TF | Unknown | HT-SELEX (Lysate) | MEME | 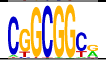 |
| NACC2 | Codebook TF | Unknown | GHT-SELEX (GFPIVT) | Dimont | 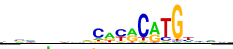 |
| DACH1 | Codebook TF | Unknown | HT-SELEX (GFPIVT) | ChIPMunk | 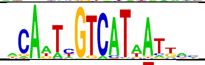 |
| DACH2 | Codebook TF | Unknown | HT-SELEX (IVT) | Autoseed | 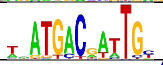 |
| PURB | Codebook TF | Unknown | PBM | ExplaiNN | 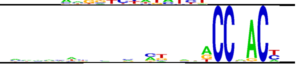 |
| TCF20 | Codebook TF | Unknown | PBM | Dimont | 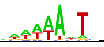 |
