## Supplementary Document 1 for "Codebook: sequence specificity and genomic binding of poorly-characterized human transcription factors"

### Description

This document contains case-by-case manual assessment of the proteins that did not yield successful experiments in the Codebook Consortium project. They are organized by DNA-binding domain (DBD) composition and result of the re-assessment.

Assessment objective is to see whether manual examination of existing knowledge, and in some cases the Codebook data itself, suggest that they are more likely than not to be potential sequence specific TFs (false negatives), or whether the available evidence instead supports them being true negatives.

In many cases of the unsuccessful proteins that we assessed as unlikely TFs, the main evidence for sequence specific DNA-binding activity is an early individual study without any further evidence. In most of these studies, the DNA binding was observed with Electrophoretic Mobility Shift Assay (EMSA) or SELEX experiments analyzed with low read count sequencing. There are many mechanisms that can lead to false positives in these types of analyzes: **1)** Binding can be indirect, in many cases the proteins have now further evidence for being transcriptional cofactors that either bind DNA indirectly or show only nonspecific binding, such as contacts with DNA backbone; **2)** Proteins can be DNA-binding, but with practically no specificity; **3)** The proteins have been binding to fully or partially single-stranded DNA, which can exist in reactions due to stoichiometric differences of annealed oligomers, or in case of SELEX, if DNA has been amplified with too many PCR cycles and denatured strands have then annealed to wrong counterparts, leading to products with correctly hybridized constant regions and partially single stranded randomized regions – this mechanism can even cause enrichment of aptameric DNA molecules; **4)** SELEX-ligand pool can have lost too much diversity or reflect a PCR-artifact and; **5)** Binding conditions may be too permissible, i.e. capturing DNA-protein interactions with too small affinity to be relevant in physiological conditions.

Based on these factors we decided that in the absence of any other definitive information, isolated sources, such as EMSA, SELEX, or single-locus reporter and ChIP-seq assays do not conclusively show that a protein has sequence specific DNA-binding activity.

All of the SELEX-associated artifacts were observed within Codebook SELEX, but they are likely rare amongst our curated experiments, because we: 1) cross-validated the results from multiple analyses; 2) implemented experimental strategies to reduce their effect (such as an additional double-stranding step carried out with additional primers after each PCR amplification); and 3) because the hundreds of parallel experiments made it possible to detect recurrent artifacts that would have seemed like a real signals if we analyzed only an individual experiment.

For each protein, we show an assessment of previous evidence, a brief summary of the Codebook experiments, a table of assays that it was analyzed with, and a link to the MEX website[1], that can be used to examine PWMs generated for the protein and their scores across the datatypes.

### C2H2 zinc finger proteins that are unlikely to be sequence specific TFs

Note that “Unconventional C2H2 domain” is defined as a domain that is described as a C2H2 zinc finger (C2H2-zf) domain in some of the databases or literature, but did not meet the recommended HMMer detection threshold of Per-sequence Eval < 0.01 and Per-domain conditional Eval < 0.01, that is used in CisBP (Pfam model: PF00096).

**FAM170A (Unconventional C2H2) is not likely to be a TF.** This protein was included based on it containing an unconventional C2H2-zf domain. A single study predicted that it contains an atypical C2H2-zf domain, localizes to the nucleus and overexpression upregulates other genes[2]. However, based on AlphaFold3 (AF3), the protein does not contain any C2H2-zf domains.
**Codebook experiment summary**: No PBM or ChIP-seq experiments passed quality control. All other experiment types were performed but none of them yielded specific PWMs. FAM170A

| FAM170A | ChIP | GHT | HT | SMS | PBM | Lysate | eGFP-IVT | IVT |
|---|---|---|---|---|---|---|---|---|
| Tested? | TRUE | TRUE | TRUE | TRUE | FALSE | TRUE | TRUE | TRUE |

**KIN (Unconventional C2H2) is not likely to be a TF.** This protein was included based on it containing an unconventional C2H2-zf domain. It is also known as "KIN17 DNA and RNA binding protein". Based on previous EMSA data the protein binds curved DNA associated with erroneous recombination[3], whereas recent evidence point towards it regulating transcription of ribosomal RNA encoding genes[4]. The C2H2-like domain is a specific “KIN17” C2H2 (IPR056767). There are multiple lines of evidence that this protein binds nucleic acids in the context of DNA metabolism, but no indications pointing to sequence specificity or regulation of transcription[5]. AF3 predicts that the single C2H2 folds properly, but also that it interacts with other regions of the protein, which could interfere with the DNA-binding activity.
**Codebook experiment summary**: No PBM. All other experiment types were performed but none of them yielded specific PWMs. KIN

| KIN | ChIP | GHT | HT | SMS | PBM | Lysate | eGFP-IVT | IVT |
|---|---|---|---|---|---|---|---|---|
| Tested? | TRUE | TRUE | TRUE | TRUE | FALSE | TRUE | TRUE | TRUE |

**TRAFD1 (Unconventional C2H2) is not likely to be a TF.** This protein was included based on it containing an unconventional C2H2-zf domain. It is described as negatively regulating IRF3 and NFKB, but there is no evidence for DNA-binding, and it appears to operate upstream in the signalling cascade[6]. The C2H2-like domain is IPR049439 (TRAFD1/XAF1, zinc finger). AF3 predicts that the protein has 4 canonically folded

C2H2-like domains, but they are located immediately adjacent to each other, which is likely incompatible for binding to DNA as a C2H2-zf array.
**Codebook experiment summary**: No PBM or ChIP-seq experiments passed quality control. All experiment types were performed but none of them yielded specific PWMs. TRAFD1

| TRAFD1 | ChIP | GHT | HT | SMS | PBM | Lysate | eGFP-IVT | IVT |
|---|---|---|---|---|---|---|---|---|
| Tested? | TRUE | TRUE | TRUE | TRUE | FALSE | TRUE | TRUE | TRUE |

**ZNF326 (Unconventional C2H2) is likely an RNA binding protein.** This protein was included based on it containing an unconventional C2H2-zf domain. ZNF326 is a member of the DBIRD complex that regulates splicing by binding to PolII in an RNA-independent manner[7]. The protein domain is of "AKAP95" C2H2 subtype as in AKAP8 proteins. In the AF3 prediction, the C2H2-zf domain is followed immediately by an extra alpha-helix.
**Codebook experiment summary**: No Lysate SELEX or PBM. ChIP-seq did not pass quality control. All experiment types were performed but none of them yielded specific PWMs. ZNF326

| ZNF326 | ChIP | GHT | HT | SMS | PBM | Lysate | eGFP-IVT | IVT |
|---|---|---|---|---|---|---|---|---|
| Tested? | TRUE | TRUE | TRUE | TRUE | FALSE | FALSE | TRUE | TRUE |

**ZNF474 (Unconventional C2H2) is not likely to be a TF.** This protein contains a single C2H2-zf domain based on our standard threshold and multiple "C2HC/C3H-type" domains (IPR049899) according to pfam predictions. AF3 predicts that the most C-terminal domain is folded canonically, whereas three other domains have noncanonical features, including a much longer recognition helix and folded domains adjacent to both sides of the C2H2-zf domain. This structure appears incompatible with the established C2H2 DNA binding mode.
**Codebook experiment summary**: No PBM or ChIP-seq experiments passed quality control. None of the other experiments enriched specific PWMs. ZNF474

| ZNF474 | ChIP | GHT | HT | SMS | PBM | Lysate | eGFP-IVT | IVT |
|---|---|---|---|---|---|---|---|---|
| Tested? | TRUE | TRUE | TRUE | TRUE | FALSE | TRUE | TRUE | TRUE |

**ANKZF1 (Single C2H2) is not likely to be a TF.** This protein was included based on it containing a potential C2H2-zf. AF3 predicts the correct fold for the putative C2H2-zf, but the protein is known to be an enzyme that cleaves polypeptidyl-tRNAs.
**Codebook experiment summary**: All experiment types were performed, but none of them yielded specific PWMs. ANKZF1

| ANKZF1 | ChIP | GHT | HT | SMS | PBM | Lysate | eGFP-IVT | IVT |
|---|---|---|---|---|---|---|---|---|
| Tested? | TRUE | TRUE | TRUE | TRUE | TRUE | TRUE | TRUE | TRUE |

**CPXCR1 (Single C2H2) is not likely to be a TF.** This protein was included based on it containing a potential C2H2-zf. Very little is known about this protein except that it is associated with cleft palate. Based on STRING database the protein it interacts with endo- and lysosomal proteins VTA1 (Vesicle Trafficking 1) and CHMP3 (Charged Multivesicular Body Protein 3). AF3 does not predict correct fold for the putative C2H2-zf sequence.
**Codebook experiment summary**: All experiment types were performed but none of them yielded specific PWMs. CPXCR1

| CPXCR1 | ChIP | GHT | HT | SMS | PBM | Lysate | eGFP-IVT | IVT |
|---|---|---|---|---|---|---|---|---|
| Tested? | TRUE | TRUE | TRUE | TRUE | TRUE | TRUE | TRUE | TRUE |

**DZIP1 (Single C2H2) is not likely to be a TF.** The C2H2-zf domain is folded properly based on AF3 prediction, but this protein is cytosolic, and has a known role in organization of cilia[8].
**Codebook experiment summary**: All experiment types were performed but none of them yielded specific PWMs. DZIP1

| DZIP1 | ChIP | GHT | HT | SMS | PBM | Lysate | eGFP-IVT | IVT |
|---|---|---|---|---|---|---|---|---|
| Tested? | TRUE | TRUE | TRUE | TRUE | TRUE | TRUE | TRUE | TRUE |

**EEA1 (Single C2H2) is not likely to be a TF.** This protein was included based on it containing a potential C2H2-zf domain. The C2H2-zf domain is folded properly based on AF3 prediction. It is known to be a cytosolic and endosomal protein, however, that associates with RAB5A and membranes, and binds phospholipid vesicles[9].
**Codebook experiment summary**: No ChIP-seq or Lysate SELEX experiments. None of the other experiments enriched specific PWMs. EEA1

| EEA1 | ChIP | GHT | HT | SMS | PBM | Lysate | eGFP-IVT | IVT |
|---|---|---|---|---|---|---|---|---|
| Tested? | FALSE | TRUE | TRUE | TRUE | TRUE | FALSE | TRUE | TRUE |

**L3MBTL4 (Single C2H2) is likely a transcriptional cofactor.** The putative C2H2-zf domain corresponding sequence does not fold into a C2H2-zf domain based on AF3, but is instead a part of globular domain. Its closest paralog L3MBTL3 interacts with histone demethylase KDM1A and RBPJ to repress enhancers[10]. Consistently with its

paralog L3MBTL3, it makes protein–protein interactions with chromatin proteins, including RBPJ. An RBPJ-like PWM is enriched in our ChIP-seq data.
**Codebook experiment summary**: No PBM experiment. ChIP-seq experiments show enrichment of RBPJ-like PWMs with “RNTCCCRG” consensus. There is no clear PWM enrichment in other data. L3MBTL4

| L3MBTL4 | ChIP | GHT | HT | SMS | PBM | Lysate | eGFP-IVT | IVT |
|---|---|---|---|---|---|---|---|---|
| Tested? | TRUE | TRUE | TRUE | TRUE | FALSE | TRUE | TRUE | TRUE |

**PRMT3 (Single C2H2) is likely a transcriptional cofactor.** PRMT3 is a protein arginine methyltransferase that is known to modify many proteins, including histone H4[11]. Based on the NMR structure of the mouse homolog, PDB:1WIR[12], the predicted C2H2-zf sequence is a part of larger folded protein domain and not a real C2H2-zf domain.
**Codebook experiment summary**: No PBM or ChIP-seq experiments passed quality control. None of the other experiments enriched specific PWMs. PRMT3

| PRMT3 | ChIP | GHT | HT | SMS | PBM | Lysate | eGFP-IVT | IVT |
|---|---|---|---|---|---|---|---|---|
| Tested? | TRUE | TRUE | TRUE | TRUE | FALSE | TRUE | TRUE | TRUE |

**RBSN (Single C2H2) is not likely to be a TF.** This protein was included based on it containing a potential C2H2-zf. The C2H2-like sequence folds properly based on AF3. It is also known as rabenosyn, and it has a well-established role in membrane trafficking and recycling of endosomes. The Codebook ChIP-seq data produced peaks, suggesting that some proportion of the protein is localized in nucleus and that it may have chromatin associated roles.
**Codebook experiment summary**: No PBM experiment. ChIP-seq reads are enriched in CG-rich sequences, and lysate-based SELEX reads are enriched in NRF1 sites. None of the other experiments enriched specific PWMs. RBSN

| RBSN | ChIP | GHT | HT | SMS | PBM | Lysate | eGFP-IVT | IVT |
|---|---|---|---|---|---|---|---|---|
| Tested? | TRUE | TRUE | TRUE | TRUE | FALSE | TRUE | TRUE | TRUE |

**ZNF385A and ZNF385B (Single C2H2) are potential RNA-binding proteins**. Many studies have shown that these paralogs function as RNA-binding proteins[13,14]. The SMART database lists three Matrin U1-like C2H2-zf domains for these proteins, and these are supported by AF3 prediction that shows three C2H2-like folded domains, but with an additional alpha-helix following the C2H2-zf domain, and the domains are separated by long gaps, which would be incompatible with the established C2H2-zf DNA-binding mode.

**Codebook experiment summary**: None of the experiments enriched specific PWMs. ZNF385A, ZNF385B

| ZNF385A | ChIP | GHT | HT | SMS | PBM | Lysate | eGFP-IVT | IVT |
|---|---|---|---|---|---|---|---|---|
| Tested? | TRUE | TRUE | TRUE | FALSE | FALSE | TRUE | FALSE | TRUE |
| ZNF385B | ChIP | GHT | HT | SMS | PBM | Lysate | eGFP-IVT | IVT |
| Tested? | TRUE | TRUE | TRUE | TRUE | FALSE | TRUE | TRUE | TRUE |

**ZNF804A (Single C2H2) is not likely to be a TF.** This protein was included based on it containing a potential C2H2-zf. Protein-protein interactions show that it interacts with many alternative splicing associated RNA-binding proteins[15]. The protein has been also shown to bind to STAT2 and it potentially affects its transport from cytoplasm to nucleus as a part of interferon response[16].
**Codebook experiment summary**: No PBM or ChIP-seq experiments passed quality control. None of the other experiments enriched specific PWMs. ZNF804A

| ZNF804A | ChIP | GHT | HT | SMS | PBM | Lysate | eGFP-IVT | IVT |
|---|---|---|---|---|---|---|---|---|
| Tested? | TRUE | TRUE | TRUE | TRUE | FALSE | TRUE | TRUE | TRUE |

**ZMAT4 (C2H2 with noncanonical gaps) is not likely to be a TF.** This protein has been annotated to have four (4) potential C2H2-zf domains. The C2H2-zf domains are all “Matrin U1” domains (IPR003604), however, which are associated with RNA binding, and have an extra alpha helix adjacent to the C2H2-zf domain in AF3 predictions. ZMAT4 interacts with RNA-binding- and ubiquitinoylation-associated proteins, according to the STRING database.
**Codebook experiment summary**: No Lysate SELEX or PBM experiments. ChIP-seq did not pass quality control. None of the other experiments enriched specific PWMs. ZMAT4

| ZMAT4 | ChIP | GHT | HT | SMS | PBM | Lysate | eGFP-IVT | IVT |
|---|---|---|---|---|---|---|---|---|
| Tested? | TRUE | TRUE | TRUE | TRUE | FALSE | FALSE | TRUE | TRUE |

**ZNF207 (C2H2 with noncanonical gaps) is not likely to be a TF.** This protein has been annotated to have two (2) potential (and overlapping) C2H2-zf domains. AF3 predicts that the overlapping C2H2-zfs do not fold into a C2H2-zf domain, but instead into a completely different protein fold. ZNF207 also has also a very well characterized role as a kinetochore- and microtubule-binding protein[17,18].
**Codebook experiment summary**: No eGFP-IVT SELEX or PBM experiments. All other types of experiments were performed and none enriched specific PWMs. ZNF207

| ZNF207 | ChIP | GHT | HT | SMS | PBM | Lysate | eGFP-IVT | IVT |
|---|---|---|---|---|---|---|---|---|

| | | | | | | | | |
|---|---|---|---|---|---|---|---|---|
| Tested? | TRUE | TRUE | TRUE | TRUE | FALSE | TRUE | FALSE | TRUE |

**ZNF318 (C2H2 with noncanonical gaps) is not likely to be a TF.** This protein has been annotated to have two (2) potential C2H2-zf domains, which are also Matrin U1-type zinc finger domains[19] (IPR003604), typically associated with RNA binding. Based on AF3, these domains do not fold into individual C2H2-zf domains, but instead, fold together with additional structural features.
**Codebook experiment summary**: No PBM experiment. None of the other experiments enriched specific PWMs. ZNF318

| ZNF318 | ChIP | GHT | HT | SMS | PBM | Lysate | eGFP-IVT | IVT |
|---|---|---|---|---|---|---|---|---|
| Tested? | TRUE | TRUE | TRUE | TRUE | FALSE | TRUE | TRUE | TRUE |

**ZNF385C (C2H2 with noncanonical gaps) is not likely to be a TF.** This protein has been annotated to have two (2) potential C2H2-zf domains. The SMART-database lists it as having four Matrin U1 subtype C2H2-zf domains (IPR003604), however, which are associated with RNA binding. The C2H2-like domains do fold into the expected structure, but they have an extra alpha helix on AF3 predictions, which would be inconsistent with the typical C2H2-zf DNA-binding mode.
**Codebook experiment summary**: No PBM or ChIP-seq experiments passed quality control. None of the other experiments enriched specific PWMs. ZNF385C

| ZNF385C | ChIP | GHT | HT | SMS | PBM | Lysate | eGFP-IVT | IVT |
|---|---|---|---|---|---|---|---|---|
| Tested? | TRUE | TRUE | TRUE | TRUE | FALSE | TRUE | TRUE | TRUE |

**ZNF598 (C2H2 with noncanonical gaps) is not likely to be a TF.** This protein has been annotated as having five (5) potential C2H2-zf domains. A subset fold in the expected manner on AF3. This protein is a known ubiquitin ligase, however, which functions in translation quality control.
**Codebook experiment summary**: No PBM or ChIP-seq experiments passed quality control. None of the other experiments enriched specific PWMs. ZNF598

| ZNF598 | ChIP | GHT | HT | SMS | PBM | Lysate | eGFP-IVT | IVT |
|---|---|---|---|---|---|---|---|---|
| Tested? | TRUE | TRUE | TRUE | TRUE | FALSE | TRUE | TRUE | TRUE |

**ZUP1 (C2H2 with noncanonical gaps) is a potential RNA-binding protein**. This protein has two potential C2H2-zf domains, which fold as expected on AF3. It has been reported to use its zinc fingers as a viral RNA-binding protein that functions as a part of

the MAVS complex, which activates IRF3 and NFKB TFs[20]. The protein also has also well-established role as a de-ubiquitinase functioning in DNA damage response[20].
**Codebook experiment summary**: No eGFP-IVT SELEX or PBM experiments. ChIP-seq did not pass quality control. None of the other experiments enriched specific PWMs.
ZUP1

| ZUP1 | ChIP | GHT | HT | SMS | PBM | Lysate | eGFP-IVT | IVT |
|---|---|---|---|---|---|---|---|---|
| Tested? | TRUE | TRUE | TRUE | TRUE | FALSE | TRUE | FALSE | TRUE |

### C2H2 zinc finger proteins that are potential sequence specific TFs

**AKAP8 and AKAP8L (Unconventional C2H2) are potential TFs.** These proteins were included based on containing an unconventional C2H2-zf domain. AKAP8 is also known as AKAP95 and is associated with many different roles. In a previous study, AKAP8/AKAP/95 bound to G/C-rich DNA according to SELEX[21] and associates with rDNA promoters according to Interpro. Their C2H2-zf domain is of type “AKAP95-type 2 domain” that is associated with RNA binding. Other lines of evidence, however point towards them being RNA-binding proteins are that they are associated with alternative splicing[22] or participating in subcellular targeting of cyclic AMP-dependent protein kinase A[23].
**Codebook experiment summary**: All experiment types were performed but none of them yielded specific PWMs. AKAP8, AKAP8L

| AKAP8 | ChIP | GHT | HT | SMS | PBM | Lysate | eGFP-IVT | IVT |
|---|---|---|---|---|---|---|---|---|
| Tested? | TRUE | TRUE | TRUE | TRUE | TRUE | TRUE | TRUE | TRUE |
| AKAP8L | ChIP | GHT | HT | SMS | PBM | Lysate | eGFP-IVT | IVT |
| Tested? | TRUE | TRUE | TRUE | TRUE | TRUE | TRUE | TRUE | TRUE |

**ZNF706 (Unconventional C2H2) is a potential TF.** This protein was included based on containing an unconventional C2H2-zf domain. This tiny protein (76AA) works in controlling stem cell state via regulating KLF4 expression[24]. The study establishing this role did not decipher the mechanism, however.
**Codebook experiment summary**: No PBM or ChIP-seq experiment. Other experiment trypes were performed but without specific PWM enrichment. ZNF706

| ZNF706 | ChIP | GHT | HT | SMS | PBM | Lysate | eGFP-IVT | IVT |
|---|---|---|---|---|---|---|---|---|
| Tested? | FALSE | TRUE | TRUE | TRUE | FALSE | TRUE | TRUE | TRUE |

**ZNF750 (Unconventional C2H2) is likely a transcriptional cofactor and potentially a TF.** The main source of evidence for the role of ZNF750 as a TF is that it has a potential C2H2-zf domain, which is classified as a specific subtype “Zinc finger protein 750-like” (IPR039064). AF3 does not predict any C2H2-zf folded regions within the protein, however. A ChIP-seq study concluded that it binds “NNSCYNNRGGSW” motif, which is indistinguishable from target specificities of TFAP2 proteins. The same study has also mass-spectrometry data that shows that this protein interacts strongly with TFAP2A and thus the binding is likely indirect[25]. The same study also indicates that it interacts with KLF4, and consistent with this observation, an earlier study shows that it may Influence KLF4 expression[26].

**Codebook experiment summary**: No PBM experiment. All experiment types were performed but none of them yielded specific PWMs. ZNF750

| ZNF750 | ChIP | GHT | HT | SMS | PBM | Lysate | eGFP-IVT | IVT |
|---|---|---|---|---|---|---|---|---|
| Tested? | TRUE | TRUE | TRUE | TRUE | FALSE | TRUE | TRUE | TRUE |

**CCDC17 (Single C2H2) is likely a transcriptional cofactor and a potential TF.** This protein was included based on it containing a potential C2H2-zf domain. Very little is known about this protein except that it makes strong protein–protein interactions with RNA PolII and cohesin components. The C2H2-zf domain is folded properly based on AF3 prediction.
**Codebook experiment summary**: No ChIP-seq experiment. Other experiments were performed but without specific PWM enrichment. CCDC17

| CCDC17 | ChIP | GHT | HT | SMS | PBM | Lysate | eGFP-IVT | IVT |
|---|---|---|---|---|---|---|---|---|
| Tested? | FALSE | TRUE | TRUE | TRUE | TRUE | TRUE | TRUE | TRUE |

**ZMAT1 (Single C2H2) is a potential TF.** This protein was included based on it containing a potential C2H2-zf. Little is known about this protein; It interacts mostly with RBPs and has been suggested to function either as RBP or TF. On AF3, the N-terminal sequence folds into a C2H2-zf fold that is followed directly by a long alpha-helix.
**Codebook experiment summary**: No SMS or PBM experiments. ChIP-seq did not pass quality control. All other experiment types were performed but none of them yielded specific PWMs. ZMAT1

| ZMAT1 | ChIP | GHT | HT | SMS | PBM | Lysate | eGFP-IVT | IVT |
|---|---|---|---|---|---|---|---|---|
| Tested? | TRUE | TRUE | TRUE | FALSE | FALSE | TRUE | TRUE | TRUE |

**ZNF365 (Single C2H2) is a potential TF.** The C2H2-zf domain in this protein mediates interaction between it and a complex that contains PARP1 and MRE11[27]. ZNF365 contains a Ku C terminal like domain that is thought to be involved in DNA-damage repair pathways. AF3 predicts that the predicted C2H2-zf domain folds properly.
**Codebook experiment summary**: No ChIP-seq or PBM experiments. Other experiments were performed but without specific PWM enrichment. ZNF365

| ZNF365 | ChIP | GHT | HT | SMS | PBM | Lysate | eGFP-IVT | IVT |
|---|---|---|---|---|---|---|---|---|
| Tested? | FALSE | TRUE | TRUE | TRUE | FALSE | TRUE | TRUE | TRUE |

**ZNF428 (Single C2H2) is a potential TF.** This protein was included based on it containing a potential C2H2-zf. Little is known about this protein, but AF3 predicts that the C2H2-zf is folded properly, and that it is the only folded domain in the protein.
**Codebook experiment summary**: No PBM experiment. eGFP-IVT based HT-SELEX enriched specific "CCGTA" motifs but the enrichment was weak and none of the other experiments supported the result. ZNF428

| ZNF428 | ChIP | GHT | HT | SMS | PBM | Lysate | eGFP-IVT | IVT |
|---|---|---|---|---|---|---|---|---|
| Tested? | TRUE | TRUE | TRUE | TRUE | FALSE | TRUE | TRUE | TRUE |

**ZNF608 (Single C2H2) is a potential TF.** This protein was included based on it containing a potential C2H2-zf. Little is known about this protein. AF3 predicts that the C2H2-zf is folded properly.
**Codebook experiment summary**: No PBM experiments. All other experiment types were performed but without specific PWM enrichment. ZNF608

| ZNF608 | ChIP | GHT | HT | SMS | PBM | Lysate | eGFP-IVT | IVT |
|---|---|---|---|---|---|---|---|---|
| Tested? | TRUE | TRUE | TRUE | TRUE | FALSE | TRUE | TRUE | TRUE |

**ZNF609 (Single C2H2) is a potential TF.** This protein was included based on it containing a potential C2H2-zf. Little is known about this protein. AF3 predicts that the C2H2-zf is folded properly. The corresponding gene makes a circ-RNA that has recently been studied extensively.
**Codebook experiment summary**: No PBM or ChIP-seq experiments passed quality control. All experiment types were performed but none of them yielded specific PWMs. ZNF609

| ZNF609 | ChIP | GHT | HT | SMS | PBM | Lysate | eGFP-IVT | IVT |
|---|---|---|---|---|---|---|---|---|
| Tested? | TRUE | TRUE | TRUE | TRUE | FALSE | TRUE | TRUE | TRUE |

**KAT7 (Single C2H2) is a transcriptional cofactor and potentially a TF**. This protein was included based on it containing a potential C2H2-zf domain. However, the protein is a well-established H3K14 acetyltransferase. AF3 also predicts that its putative C2H2-zf sequence does not fold into a C2H2-zf domain. There is an external ChIP-seq-derived CG-rich PWM, which is potentially a false positive derived from indirect binding.
**Codebook experiment summary**: No PBM experiment. ChIP-seq enriches weakly CG-rich PWMs but there is no PWM enrichment in other datasets. KAT7

| KAT7 | ChIP | GHT | HT | SMS | PBM | Lysate | eGFP-IVT | IVT |
|---|---|---|---|---|---|---|---|---|
| Tested? | TRUE | TRUE | TRUE | TRUE | FALSE | TRUE | TRUE | TRUE |

**PEG3 (C2H2 with noncanonical gaps) is a potential transcriptional factor.** This protein has been annotated to have 11 potential C2H2-zf domains. Previous studies have analyzed the target specificity of PEG3 with ChIP-seq and EMSA[28], and ChIP-qPCR and EMSA[29]. PWM construction in these studies involved non-standard data analysis methods, and the resulting PWMs are in disagreement. AF3 predicts that some of the putative 13 C2H2-zf domains fold correctly.
**Codebook experiment summary**: No PBM experiment. All other experiment types were performed but without specific PWM enrichment. PEG3

| PEG3 | ChIP | GHT | HT | SMS | PBM | Lysate | eGFP-IVT | IVT |
|---|---|---|---|---|---|---|---|---|
| Tested? | TRUE | TRUE | TRUE | TRUE | FALSE | TRUE | TRUE | TRUE |

**PRDM8 (C2H2 with noncanonical gaps) is a transcriptional cofactor and potentially a TF.** This protein has been annotated to have 2 potential C2H2-zf domains. The same study that established PRDM8 as a histone methyltransferase (H3K9me3) included ChIP-seq, which did not result in a specific PWMs[30]. AF3 predicts that both C2H2-zf domains fold correctly, but the first one has a very long recognition helix and the second one is in the extreme C-terminus of the protein.
**Codebook experiment summary**: No PBM experiment. All other experiment types were performed but without specific PWM enrichment. PRDM8

| PRDM8 | ChIP | GHT | HT | SMS | PBM | Lysate | eGFP-IVT | IVT |
|---|---|---|---|---|---|---|---|---|
| Tested? | TRUE | TRUE | TRUE | TRUE | FALSE | TRUE | TRUE | TRUE |

**ZFPM2 (C2H2 with noncanonical gaps) is a transcriptional cofactor and potentially a TF.** This protein has been annotated to have 3 potential C2H2-zf domains. ZFPM2 is also known as Friend Of GATA 2 (FOG2), and FOGs are known to use some of their zinc fingers to interact with the GATA proteins and other TFs[31]. It is still unclear whether these proteins have intrinsic target specificity. Its paralog ZFPM1 enriched a specific GATA-consensus PWM. The protein used in that experiment was produced in wheat germ extract system and it could be co-precipitating a wheat GATA protein. AF3 predicts that the protein has more than three domains with C2H2-like fold.
**Codebook experiment summary**: No PBM experiment. All other experiment types were performed but without specific PWM enrichment. ZFPM2

| ZFPM2 | ChIP | GHT | HT | SMS | PBM | Lysate | eGFP-IVT | IVT |
|---|---|---|---|---|---|---|---|---|
| Tested? | TRUE | TRUE | TRUE | TRUE | FALSE | TRUE | TRUE | TRUE |

**ZNF644 (C2H2 with noncanonical gaps) is a potential TF.** This protein has been annotated to have 2 potential C2H2-zf domains. The C2H2-zf domains fold correctly in AF3 predictions. Very little is known about the protein.

**Codebook experiment summary**: No PBM experiment. ChIP-seq reads are enriched in YY1 sites. None of the other experiment types yielded specific PWMs. ZNF644

| ZNF644 | ChIP | GHT | HT | SMS | PBM | Lysate | eGFP-IVT | IVT |
|---|---|---|---|---|---|---|---|---|
| Tested? | TRUE | TRUE | TRUE | TRUE | FALSE | TRUE | TRUE | TRUE |

### Unsuccessful proteins with canonical C2H2 arrays (potential TFs)

The following proteins contain standard C2H2-zf arrays. We did not annotate them on a case-by-case basis. They are all potential TFs.

| TF name | #C2H2 | ChIP | GHT | HT | SMS | PBM | Lysate | eGFP-IVT | IVT |
|---|---|---|---|---|---|---|---|---|---|
| AC008770 | 13 | TRUE | TRUE | TRUE | TRUE | TRUE | FALSE | TRUE | TRUE |
| CHAMP1 | 5 | TRUE | TRUE | TRUE | TRUE | TRUE | TRUE | TRUE | TRUE |
| JAZF1 | 2 | TRUE | TRUE | TRUE | TRUE | FALSE | TRUE | TRUE | TRUE |
| PRDM2 | 5 | TRUE | TRUE | TRUE | TRUE | FALSE | TRUE | TRUE | TRUE |
| ZBTB46 | 3 | TRUE | TRUE | TRUE | TRUE | FALSE | TRUE | TRUE | TRUE |
| ZNF230 | 8 | TRUE | TRUE | TRUE | TRUE | FALSE | TRUE | TRUE | TRUE |
| ZNF280B | 6 | TRUE | TRUE | TRUE | TRUE | FALSE | TRUE | TRUE | TRUE |
| ZNF280D | 9 | TRUE | TRUE | TRUE | TRUE | FALSE | TRUE | TRUE | TRUE |
| ZNF446 | 3 | TRUE | TRUE | TRUE | TRUE | FALSE | TRUE | TRUE | TRUE |
| ZNF469 | 3 | TRUE | TRUE | TRUE | TRUE | FALSE | FALSE | TRUE | TRUE |
| ZNF579 | 9 | TRUE | TRUE | TRUE | TRUE | FALSE | FALSE | FALSE | TRUE |
| ZNF592 | 5 | TRUE | TRUE | TRUE | TRUE | FALSE | TRUE | TRUE | TRUE |
| ZNF630 | 13 | TRUE | TRUE | TRUE | TRUE | FALSE | TRUE | TRUE | TRUE |
| ZNF639 | 8 | TRUE | TRUE | TRUE | TRUE | FALSE | TRUE | TRUE | TRUE |
| ZNF654 | 3 | TRUE | TRUE | TRUE | TRUE | FALSE | TRUE | TRUE | TRUE |
| ZNF687 | 10 | TRUE | TRUE | TRUE | TRUE | FALSE | TRUE | TRUE | TRUE |
| ZNF688 | 2 | TRUE | TRUE | TRUE | TRUE | FALSE | TRUE | TRUE | TRUE |
| ZNF717 | 21 | TRUE | TRUE | TRUE | TRUE | FALSE | TRUE | TRUE | TRUE |
| ZNF763 | 6 | TRUE | TRUE | TRUE | TRUE | FALSE | TRUE | TRUE | TRUE |
| ZNF781 | 4 | TRUE | TRUE | TRUE | TRUE | FALSE | TRUE | TRUE | TRUE |
| ZNF844 | 7 | TRUE | TRUE | TRUE | TRUE | FALSE | TRUE | TRUE | TRUE |
| ZNF91 | 33 | FALSE | TRUE | TRUE | TRUE | FALSE | FALSE | FALSE | TRUE |
| ZSCAN18 | 2 | TRUE | TRUE | TRUE | TRUE | FALSE | TRUE | FALSE | TRUE |

### Other known-DBD-containing proteins that are unlikely to be sequence specific TFs

**ARID2 (ARID/BRIGHT; RFX) is likely a low specificity transcriptional cofactor**. This protein has an ARID/BRIGHT DBD. It participates as a component in some forms of the SWI/SNF remodeler[32]. In the structures existing for this protein[33], it either does not contact DNA (PDB:7VDV) but instead binds the histone core, or makes very minor backbone contacts (PDB:7Y8R). The structures contain only a part of this protein, and there is no clear evidence that would rule out sequence specific DNA binding. Among ARID family proteins, however, only ARID3 and ARID5 subgroups are known to possess strong sequence specificity.
**Codebook experiment summary**: No SMS experiment. The protein showed strong enrichment for CTCF sites in ChIP-seq. All other experiment types were performed but without specific PWM enrichment. ARID2

| ARID2 | ChIP | GHT | HT | SMS | PBM | Lysate | eGFP-IVT | IVT |
|---|---|---|---|---|---|---|---|---|
| Tested? | TRUE | TRUE | TRUE | FALSE | TRUE | TRUE | TRUE | TRUE |

**AHDC1 (AT hook) is likely a low specificity transcriptional cofactor**. This protein has an AT-hook DNA-binding peptide motif, which would suggest interaction with DNA, but on the basis of the fact that it has many protein-protein interactions with other TFs (including dozens of C2H2-zf proteins), it could be a cofactor.
**Codebook experiment summary**: Only ChIP-seq and Lysate-based SELEX experiments. ChIP-seq yielded motif, but is not supported by other experiments. AHDC1

| AHDC1 | ChIP | GHT | HT | SMS | PBM | Lysate | eGFP-IVT | IVT |
|---|---|---|---|---|---|---|---|---|
| Tested? | TRUE | TRUE | TRUE | FALSE | FALSE | TRUE | FALSE | FALSE |

**AKNA (AT hook) is not likely to be a TF**. This protein has an AT-hook DNA-binding peptide motif. Has a well-established role in centrosome formation. It was included in the study because it has been suggested to moonlight as TF through binding AT-rich promoters of CD40 and CD40L and coordinate their expression[34].
**Codebook experiment summary**: All experiment types were performed but none of them yielded specific PWMs. AKNA

| AKNA | ChIP | GHT | HT | SMS | PBM | Lysate | eGFP-IVT | IVT |
|---|---|---|---|---|---|---|---|---|
| Tested? | TRUE | TRUE | TRUE | TRUE | TRUE | TRUE | TRUE | TRUE |

**CBX2 (AT hook) is likely a low specificity transcriptional cofactor.** This protein has an AT-hook DNA-binding peptide motif. CBX2 functions in the polycomb repressive

complex, binding trimethylated H3K27 with its chromodomain, and the AT-hook may contribute into this interaction[35].
**Codebook experiment summary**: No IVT SELEX or PBM experiments. Other experiment types were performed but without specific PWM enrichment. CBX2

| CBX2 | ChIP | GHT | HT | SMS | PBM | Lysate | eGFP-IVT | IVT |
|---|---|---|---|---|---|---|---|---|
| Tested? | TRUE | TRUE | TRUE | TRUE | FALSE | TRUE | TRUE | FALSE |

**DOT1L (AT hook) is likely a low specificity transcriptional cofactor**. This protein has an AT-hook DNA-binding peptide motif**.** It is known to be a histone methyltransferase. The putative AT-hook may contribute to DNA interactions but is unlikely to define target locations. The protein binds the histone core in PDB:6J99 and makes only very minor contacts with DNA backbone[36].
**Codebook experiment summary**: Only ChIP-seq and Lysate SELEX experiments. Other experiment types were performed but without specific PWM enrichment. DOT1L

| DOT1L | ChIP | GHT | HT | SMS | PBM | Lysate | eGFP-IVT | IVT |
|---|---|---|---|---|---|---|---|---|
| Tested? | TRUE | TRUE | TRUE | FALSE | FALSE | TRUE | FALSE | FALSE |

**PHF20 (AT hook) is likely a low specificity transcriptional cofactor**. This protein has an AT-hook DNA-binding peptide motif. PHF20 is known to bind di-methylated H3K4me2 and to function as a part of "nonspecific lethal" protein complex that regulates gene expression through histone acetyltransferase activity[37].
**Codebook experiment summary**: It has only ChIP-seq data, with no specific enrichment. PHF20

| PHF20 | ChIP | GHT | HT | SMS | PBM | Lysate | eGFP-IVT | IVT |
|---|---|---|---|---|---|---|---|---|
| Tested? | TRUE | FALSE | FALSE | FALSE | FALSE | FALSE | FALSE | FALSE |

**PRR12 (AT hook) is likely a low specificity transcriptional cofactor**. This protein has an AT-hook DNA-binding peptide motif. It has been recently shown to interact with cohesin complex through NIPBL[38]. Based on protein–protein interactions, it also associates with TFs and cofactors, consisted with a role as a transcriptional cofactor.
**Codebook experiment summary**: Only ChIP-seq and Lysate SELEX experiments. ChIP-seq reads are enriched posterior homeodomain sites. No clear enrichment in Lysate SELEX experiments. PRR12

| PRR12 | ChIP | GHT | HT | SMS | PBM | Lysate | eGFP-IVT | IVT |
|---|---|---|---|---|---|---|---|---|
| Tested? | TRUE | TRUE | TRUE | FALSE | FALSE | TRUE | FALSE | FALSE |

**SCML4 (AT hook) is likely a low specificity transcriptional cofactor**. This protein has an AT-hook DNA-binding peptide motif. Very little is known about the protein. It associates with TFs and cofactors based on protein–protein interactions.
**Codebook experiment summary**: No SMS experiment. All other types of experiments were tested, with no clear motifs. SCML4

| SCML4 | ChIP | GHT | HT | SMS | PBM | Lysate | eGFP-IVT | IVT |
|---|---|---|---|---|---|---|---|---|
| Tested? | TRUE | TRUE | TRUE | FALSE | TRUE | TRUE | TRUE | TRUE |

**SGSM2 (BED ZF) is unlikely to be a TF**. The protein was included because it was predicted to contain a ZBED domain. However, AF3 predicts that the associated sequence does not fold into a ZBED domain but is instead a part of  larger folded protein domain. So, this protein is not a ZBED. The protein has GTPase activity and is thought to function in membrane trafficking.
**Codebook experiment summary**: There is an enrichment for CTCF sites in ChIP-seq. Other experiment types were performed but without specific PWM enrichment. SGSM2

| SGSM2 | ChIP | GHT | HT | SMS | PBM | Lysate | eGFP-IVT | IVT |
|---|---|---|---|---|---|---|---|---|
| Tested? | TRUE | TRUE | TRUE | TRUE | TRUE | TRUE | TRUE | TRUE |

**NCOA1, NCOA2 and NCOA3 (bHLH) are likely non-DNA-binding transcriptional cofactors.** The main source of evidence for their role as TFs is that they have an apparent bHLH domain. There is no previous evidence of them interacting with DNA directly, however; They are best known as coactivators (for which they are named). The HLH domain is known to function as a protein-protein interaction interphase for other TFs.
**Codebook experiment summary**: The NCOA family proteins were systematically unsuccessful in Codebook experiments. Protein showed enrichment for CG-rich sites in ChIP-seq. Other experiment types were performed but without specific PWM enrichment. NCOA1, NCOA2, NCOA3

| NCOA1 | ChIP | GHT | HT | SMS | PBM | Lysate | eGFP-IVT | IVT |
|---|---|---|---|---|---|---|---|---|
| Tested? | TRUE | TRUE | TRUE | FALSE | TRUE | TRUE | TRUE | TRUE |
| NCOA2 | ChIP | GHT | HT | SMS | PBM | Lysate | eGFP-IVT | IVT |
| Tested? | TRUE | TRUE | TRUE | TRUE | TRUE | TRUE | TRUE | TRUE |
| NCOA3 | ChIP | GHT | HT | SMS | PBM | Lysate | eGFP-IVT | IVT |
| Tested? | TRUE | TRUE | TRUE | FALSE | TRUE | TRUE | TRUE | TRUE |

**ZC3H8 (CCCH ZF) is an RNA-binding protein.** A recombinant protein. EMSA experiments on the protein's mouse ortholog has indicated that it binds DNA specifically[39]. However, the protein has been shown to be a sequence specific RNA-binding protein that binds "GCUUGY" consensus based on HTR-SELEX[40], and overall, the CCCH-zf domains are known to be RNA-binding proteins. RNA-binding proteins can often also bind single-stranded DNA.
**Codebook experiment summary**: No ChIP-seq experiment. All other experiment types were performed but without specific PWM enrichment. ZC3H8

| ZC3H8 | ChIP | GHT | HT | SMS | PBM | Lysate | eGFP-IVT | IVT |
|---|---|---|---|---|---|---|---|---|
| Tested? | FALSE | TRUE | TRUE | TRUE | TRUE | TRUE | TRUE | TRUE |

**ZGPAT (CCCH ZF) is likely a transcriptional cofactor and a potential RNA-binding protein**. This protein has a CCCH-zf domain, which is best known for binding RNA. In published analyses, it binds "RRRGGAGGAGA" consensus based on SELEX performed with purified protein, and cell extract-based EMSA. As with ZC3H8 above, we note that RNA-binding proteins can often bind single-stranded DNA. The same study also shows mass spectrometry evidence that it binds NuRD complex[41].
**Codebook experiment summary**: All experiment types were performed but none of them yielded specific PWMs. ZGPAT

| ZGPAT | ChIP | GHT | HT | SMS | PBM | Lysate | eGFP-IVT | IVT |
|---|---|---|---|---|---|---|---|---|
| Tested? | TRUE | TRUE | TRUE | TRUE | TRUE | TRUE | TRUE | TRUE |

**HMGN3 (HMG/Sox) is likely a low specificity DNA-binding cofactor**. This protein has an HMG DBD of a type that is typically thought to be non-specific DNA-binding domain. HMGN3 binds nucleosomal DNA[42].
**Codebook experiment summary**: ChIP-seq did not pass quality control. Other experiment types were performed but without specific PWM enrichment. HMGN3

| HMGN3 | ChIP | GHT | HT | SMS | PBM | Lysate | eGFP-IVT | IVT |
|---|---|---|---|---|---|---|---|---|
| Tested? | TRUE | TRUE | TRUE | TRUE | TRUE | TRUE | TRUE | TRUE |

**POU5F2 (Homeodomain; POU) is not likely to be a TF.** All of the Codebook experiments were unsuccessful, whereas its closest paralog POU5F1 (that was run as a positive control) was successful in all methods except for ChIP-seq. Furthermore, there is no clear evidence of sequence specific DNA-binding activity even though its paralog, POU5F1, is one of the best characterized human TFs.
When compared against the structural data for POU5F1, all base contacting residues in POU5F2 appear to be intact. The protein has less basic intrinsically disordered spacer between the POU and the homeodomain, and the homeodomain has a shorter DNA

recognition helix that misses several K and R residues that make electrostatic interactions with DNA backbone in the POU5F1 structure. Taken together, the evidence suggests that POU5F2 has lost its sequence specific DNA-binding capability.
**Codebook experiment summary**: This protein showed strong enrichment for posterior homeodomain sites in ChIP-seq, which could be due to indirect binding, but there was no enrichment for expected POU class target sites. None of the other experiment types enriched specific PWMs. POU5F2

| POU5F2 | ChIP | GHT | HT | SMS | PBM | Lysate | eGFP-IVT | IVT |
|---|---|---|---|---|---|---|---|---|
| Tested? | TRUE | TRUE | TRUE | TRUE | TRUE | TRUE | TRUE | TRUE |

**BAZ2A (MBD) is likely a low specificity DNA-binding cofactor**. This protein encodes a subtype of the MBD domain, called a TAM domain, that can bind unmethylated DNA in a non-specific manner, as well as a potential AT-hook DNA-binding peptide motif. It binds DNA backbone (PDB:7FHJ, 7MWL, and 7MWH)[43] and also contacts acetylated histones.
**Codebook experiment summary**: Only ChIP-seq and Lysate SELEX experiments. ChIP-seq did not pass quality control. None of the other experiment types yielded specific PWMs. BAZ2A

| BAZ2A | ChIP | GHT | HT | SMS | PBM | Lysate | eGFP-IVT | IVT |
|---|---|---|---|---|---|---|---|---|
| Tested? | TRUE | TRUE | TRUE | FALSE | FALSE | TRUE | FALSE | FALSE |

**BAZ2B (MBD; AT hook) is likely a low specificity DNA-binding cofactor**. This is a close paralog of BAZ2A (above).
**Codebook experiment summary**: Only ChIP-seq and Lysate SELEX experiments, with no specific PWM enrichment. BAZ2B

| BAZ2B | ChIP | GHT | HT | SMS | PBM | Lysate | eGFP-IVT | IVT |
|---|---|---|---|---|---|---|---|---|
| Tested? | TRUE | TRUE | TRUE | FALSE | FALSE | TRUE | FALSE | FALSE |

**PIN1 (MBD) is not likely to be a TF**. A specific isoform of this protein contains a putative MBD domain; however, the isoform is not found in other animals and may be spurious. In addition, the protein is a peptidyl-prolyl cis/trans isomerase that is not associated with chromatin.
**Codebook experiment summary**: No SMS experiment. All other experiment types were tested without a clear PWM enrichment. None of the *in vitro* experiments were performed with methylated DNA. PIN1

| PIN1 | ChIP | GHT | HT | SMS | PBM | Lysate | eGFP-IVT | IVT |
|---|---|---|---|---|---|---|---|---|

| | | | | | | | | |
|---|---|---|---|---|---|---|---|---|
| Tested? | TRUE | TRUE | TRUE | FALSE | TRUE | TRUE | TRUE | TRUE |

**RFX8 (RFX) is not likely to be a TF.** This protein encodes an RFX domain, however based on structural data for paralogs, the RFX domain is truncated – it is missing a region that makes backbone contacts with DNA, and thus has likely lost its DNA-binding functionality.
**Codebook experiment summary:** ChIP-seq did not pass quality control. All experiment types were performed but none of them yielded specific PWMs. RFX8

| RFX8 | ChIP | GHT | HT | SMS | PBM | Lysate | eGFP-IVT | IVT |
|---|---|---|---|---|---|---|---|---|
| Tested? | TRUE | TRUE | TRUE | TRUE | TRUE | TRUE | TRUE | TRUE |

**THAP (THAP finger) proteins appear to be relatively low-affinity DNA binding TFs (and possibly low-specificity).** The nine THAP-proteins tested in Codebook yielded clear promoter-located peaks in the ChIP-seq experiments. The ChIP-seq data also enriched PWMs for a variety of well-known promoter TFs but did not show enrichment of specific motifs in any other experiments. We examined existing protein–DNA structures data for THAP proteins: There is an X-ray crystallography structure for *D. melanogaster* DmTHAP (**PDB: 3KDE**[44]) and an NMR structure of human THAP1 (**PDB:2KO0**[45]). The THAPs are clearly DNA-binding proteins that bind DNA with their C2CH zinc finger domains, and at least in these two examples, form also direct contacts with DNA bases. Based on previous studies, however, both dmTHAP and THAP1 bind their specific target sites with low affinities of 160nM and >480nM, respectively. We propose that DNA binding of these proteins may be dependent on the presence of other promoter-associated chromatin components.
THAP10, THAP11, THAP2, THAP4, THAP5, THAP6, THAP7, THAP8, THAP9

| THAP10 | ChIP | GHT | HT | SMS | PBM | Lysate | eGFP-IVT | IVT |
|---|---|---|---|---|---|---|---|---|
| Tested? | TRUE | TRUE | TRUE | TRUE | TRUE | TRUE | TRUE | TRUE |
| THAP11 | ChIP | GHT | HT | SMS | PBM | Lysate | eGFP-IVT | IVT |
| Tested? | TRUE | TRUE | TRUE | TRUE | TRUE | TRUE | TRUE | TRUE |
| THAP2 | ChIP | GHT | HT | SMS | PBM | Lysate | eGFP-IVT | IVT |
| Tested? | TRUE | TRUE | TRUE | TRUE | TRUE | TRUE | TRUE | TRUE |
| THAP4 | ChIP | GHT | HT | SMS | PBM | Lysate | eGFP-IVT | IVT |
| Tested? | TRUE | TRUE | TRUE | FALSE | TRUE | TRUE | TRUE | TRUE |
| THAP5 | ChIP | GHT | HT | SMS | PBM | Lysate | eGFP-IVT | IVT |
| Tested? | TRUE | TRUE | TRUE | TRUE | TRUE | TRUE | TRUE | TRUE |

| THAP6 | ChIP | GHT | HT | SMS | PBM | Lysate | eGFP-IVT | IVT |
|---|---|---|---|---|---|---|---|---|
| Tested? | TRUE | TRUE | TRUE | TRUE | TRUE | TRUE | TRUE | TRUE |
| THAP7 | ChIP | GHT | HT | SMS | PBM | Lysate | eGFP-IVT | IVT |
| Tested? | TRUE | TRUE | TRUE | TRUE | TRUE | TRUE | TRUE | TRUE |
| THAP8 | ChIP | GHT | HT | SMS | PBM | Lysate | eGFP-IVT | IVT |
| Tested? | TRUE | TRUE | TRUE | TRUE | TRUE | FALSE | TRUE | TRUE |
| THAP9 | ChIP | GHT | HT | SMS | PBM | Lysate | eGFP-IVT | IVT |
| Tested? | TRUE | TRUE | TRUE | TRUE | TRUE | FALSE | TRUE | TRUE |

### Other known-DBD-containing proteins that are potential sequence specific TFs

**KDM5B (ARID/BRIGHT) is a potential TF.** The main source of evidence for the role of this protein as a TF is its ARID-BRIGHT domain, and an external experiment that is enriched with a credible motif (**Supplementary Data 1**)
**Codebook experiment summary**: No SMS experiment. All other experiment types were performed but none of them yielded specific PWMs. KDM5B

| KDM5B | ChIP | GHT | HT | SMS | PBM | Lysate | eGFP-IVT | IVT |
|---|---|---|---|---|---|---|---|---|
| Tested? | TRUE | TRUE | TRUE | FALSE | TRUE | TRUE | TRUE | TRUE |

**ZBED3 (BED ZF) a potential TF**. This protein has a ZBED DBD. There are no indications of it not being sequence specific (See **Supplementary Information** for more discussion on this structural class).
**Codebook experiment summary**: ChIP-seq did not pass quality control. All other experiment types were performed but without specific PWM enrichment. ZBED3

| ZBED3 | ChIP | GHT | HT | SMS | PBM | Lysate | eGFP-IVT | IVT |
|---|---|---|---|---|---|---|---|---|
| Tested? | TRUE | TRUE | TRUE | TRUE | TRUE | TRUE | TRUE | TRUE |

**SOHLH1 (bHLH) is a Potential TF.** This protein has a well-established bHLH DBD. Indirect evidence suggests that it is a sequence specific TF that potentially binds DNA as a heterodimer with SOHLH2[46].
**Codebook experiment summary**: ChIP-seq did not pass quality control. All other experiment types were performed but none of them yielded specific PWMs. SOHLH1

| SOHLH1 | ChIP | GHT | HT | SMS | PBM | Lysate | eGFP-IVT | IVT |
|---|---|---|---|---|---|---|---|---|
| Tested? | TRUE | TRUE | TRUE | TRUE | TRUE | TRUE | TRUE | TRUE |

**DMRTB1 (DM) is a likely TF**. This protein has a DM domain. Some of the DM-domain-containing proteins are well-established TFs and an external experiment is enriched with a credible motif (**Supplementary Data 1**).
**Codebook experiment summary**: All experiment types were performed but none of them yielded specific PWMs. DMRTB1

| DMRTB1 | ChIP | GHT | HT | SMS | PBM | Lysate | eGFP-IVT | IVT |
|---|---|---|---|---|---|---|---|---|
| Tested? | TRUE | TRUE | TRUE | TRUE | TRUE | TRUE | TRUE | TRUE |

**GATAD2B (GATA) is a likely TF**. This protein has a GATA domain. Some of the GATA-domain proteins are well-established TFs and an external experiment is enriched with a credible motif (**Data S1**)
**Codebook experiment summary**: All experiment types were performed but none of them yielded specific PWMs. GATAD2B

| GATAD2B | ChIP | GHT | HT | SMS | PBM | Lysate | eGFP-IVT | IVT |
|---|---|---|---|---|---|---|---|---|
| Tested? | TRUE | TRUE | TRUE | TRUE | TRUE | TRUE | TRUE | TRUE |

**GTF2IRD2B (GTF2I-like and potentially also a BED-zf like domain) is likely a sequence specific TF**. We tested this protein and its close paralog GTF2IRD2, the latter of which was successful and enriched a clear motif in ChIP-seq and lysate-based SELEX experiments. Both of these proteins have a GTF2I-like DBD that binds DNA specifically based on SELEX data from an individual study that characterized GTF2I[47]. Additionally, they have a C-terminal domain belonging to IPR012337 "Ribonuclease H-like superfamily" that is related to BED-zfs and could also be the DBD that binds the observed PWM (See **Supplementary Information,** for further discussion of the BED-zf domains).
Old SELEX data for GTF2I is discordant with the PWM enriched in our GTF2IRD2 experiments. GTF2I does not have the BED-zf-like domain, and the discrepancy could be due to this. Examination of the literature experiment for GTF2I shows that it has lost diversity during the selection cycles (as many of the reads are identical) and thus, the older comparison PWM may be erroneous even if the GTF2IRD2 site is enriched by GTF2I.
**Codebook experiment summary**: ChIP-seq did not pass quality control. All other experiment types were performed but none of them were enriched in any motifs. GTF2IRD2B

| GTF2IRD2B | ChIP | GHT | HT | SMS | PBM | Lysate | eGFP-IVT | IVT |
|---|---|---|---|---|---|---|---|---|
| Tested? | TRUE | TRUE | TRUE | TRUE | TRUE | TRUE | TRUE | TRUE |

**ADNP and ADNP2 (Homeodomain) are transcriptional cofactors with potential intrinsic specificity.** The main source of evidence for their role as a TFs is that they each possess a homeodomain DBD. In addition, they are predicted to have many loosely spaced noncanonical C2H2-zf domains (predicted only by SMART). Based on AF3, some of the predicted C2H2-zf domains in ADNP fold correctly but are directly adjacent to other kind of domains or even interact with them, whereas in ADNP2, one of the C2H2-zf domains is within unstructured context. Recent research has shown that these proteins are components of ChAHP and ChAHP2 chromatin remodeling complexes, respectively, which also contain CHD4 and HP1, and function to silence repeat elements. It is conceivable that their target sites are defined mostly by H3K9 trimethylation[48], but it is also possible that they encode sequence specificity that is not detectable by our assays. ADNP, ADNP2

**Codebook experiment summary**: ChIP-seq experiments displayed a strong enrichment for CTCF target sites, and additionally, ADNP enriched in an unknown motif. They were tested and unsuccessful with all other approaches, however.

| ADNP | ChIP | GHT | HT | SMS | PBM | Lysate | eGFP-IVT | IVT |
|---|---|---|---|---|---|---|---|---|
| Tested? | TRUE | TRUE | TRUE | TRUE | TRUE | TRUE | TRUE | TRUE |
| ADNP2 | ChIP | GHT | HT | SMS | PBM | Lysate | eGFP-IVT | IVT |
| Tested? | TRUE | TRUE | TRUE | TRUE | TRUE | TRUE | TRUE | TRUE |

**NANOGNB (Homeodomain) is a potential TF.** This protein has a homeodomain DBD. Very little is known about the protein. AF3 predicts that the DNA recognition helix is much longer than what is required for binding DNA.
**Codebook experiment summary**: All experiment types were performed but none of them yielded specific PWMs. NANOGNB

| NANOGNB | ChIP | GHT | HT | SMS | PBM | Lysate | eGFP-IVT | IVT |
|---|---|---|---|---|---|---|---|---|
| Tested? | TRUE | TRUE | TRUE | TRUE | TRUE | TRUE | TRUE | TRUE |

**ZHX2 and ZHX3 (Homeodomain) are transcriptional cofactors with potential intrinsic specificity.** ZHX1, ZHX2, and ZHX3 have all four homeodomain-like domains and potentially two C2H2-zf domains. AF3 predicts that both ZHX2 and ZHX3 have a single properly folded C2H2-zf domain, whereas the putative second C2H2-zf domain is folded into a larger domain. Furthermore, it predicts that the first C2H2-zf domain interacts with other parts of the protein, potentially inhibiting it from interacting with DNA. Relatively little is known about these proteins, but they likely form homo- and heterodimers with each other, and interact with the heterotrimeric NFY TF complex, leading to repression of transcription[49]. The only direct evidence of direct DNA binding by these proteins is that ZHX1 recognizes a low information content “ACG” site in previous PBM assays[50]. Interaction with NFY, combined with large number of homeodomains, yet showing very low information content target site in a single experiment, suggests that these proteins are likely cofactors, but they could also have inherent specificity that impacts the selection or function of loci to which they are recruited.
**Codebook experiment summary**: All experiment types were performed for both proteins but without specific PWM enrichment. ZHX2, ZHX3,

| ZHX2 | ChIP | GHT | HT | SMS | PBM | Lysate | eGFP-IVT | IVT |
|---|---|---|---|---|---|---|---|---|
| Tested? | TRUE | TRUE | TRUE | TRUE | TRUE | TRUE | TRUE | TRUE |
| ZHX3 | ChIP | GHT | HT | SMS | PBM | Lysate | eGFP-IVT | IVT |
| Tested? | TRUE | TRUE | TRUE | TRUE | TRUE | TRUE | TRUE | TRUE |

**HSFX1 and HSFX2 (HSF) are potential TFs.** These proteins have an HSF domain, which is the main source of evidence for a role as a TF. Little is known about both proteins, and there is no evidence besides negative results in our assays to indicate that they are not TFs.
**Codebook experiment summary**: ChIP-seq did not pass quality control. All other experiment types were performed but none of them yielded specific PWMs. HSFX1, HSFX2

| HSFX1 | ChIP | GHT | HT | SMS | PBM | Lysate | eGFP-IVT | IVT |
|---|---|---|---|---|---|---|---|---|
| Tested? | TRUE | TRUE | TRUE | TRUE | TRUE | TRUE | TRUE | TRUE |
| HSFX2 | ChIP | GHT | HT | SMS | PBM | Lysate | eGFP-IVT | IVT |
| Tested? | TRUE | TRUE | TRUE | TRUE | TRUE | TRUE | TRUE | TRUE |

**MBD4 and MBD6 (MBD) are likely sequence specific TFs.** These proteins have an MBD domain. MBD4 binds CpG hemi-methylated DNA (one of the chains is methylated and the second one is not) in the crystal structure PDB:2MOE[51]. Some sources suggest that MBD6 does not bind methylated DNA and the MBD domain instead facilitates protein-protein interactions[52].
**Codebook experiment summary**: No PWM enrichment in any assay; *in vitro* experiments were performed with unmethylated DNA. MBD4, MBD6

| MBD4 | ChIP | GHT | HT | SMS | PBM | Lysate | eGFP-IVT | IVT |
|---|---|---|---|---|---|---|---|---|
| Tested? | TRUE | TRUE | TRUE | TRUE | TRUE | TRUE | TRUE | TRUE |
| MBD6 | ChIP | GHT | HT | SMS | PBM | Lysate | eGFP-IVT | IVT |
| Tested? | TRUE | TRUE | TRUE | TRUE | TRUE | TRUE | TRUE | TRUE |

**SETDB2 (MBD) is likely a sequence specific TF.** This protein has a MBD domain, which is the main source of evidence for a role as a TF. At least some MBD domains bind CpG-methylated DNA.
**Codebook experiment summary**: ChIP-seq did not pass quality control. All experiment types were performed but none of them yielded specific PWMs. None of the *in vitro* experiments were performed with methylated DNA. SETDB2

| SETDB2 | ChIP | GHT | HT | SMS | PBM | Lysate | eGFP-IVT | IVT |
|---|---|---|---|---|---|---|---|---|
| Tested? | TRUE | TRUE | TRUE | TRUE | TRUE | TRUE | TRUE | TRUE |

**MTERF2, MTERF3, and MTERF4 (mTERF) are Potential TFs**. The main source of evidence for a role as a TF is that they have an mTERF DBD. DNA recognition of the

most well characterized paralog MTERF1 has been characterized with a crystal structure **PDB:3MVA**[53]. MTERF1 terminates mitochondrial transcription through binding DNA with a mechanism that leads to a large strain on the DNA shape and flips out a base pair from the double-helical context. The binding site is very long.
**Codebook experiment summary**: No SMS experiment. ChIP-seq did not pass quality control. All other experiment types were performed but none of them yielded specific PWMs. MTERF2, MTERF3, MTERF4

| MTERF2 | ChIP | GHT | HT | SMS | PBM | Lysate | eGFP-IVT | IVT |
|---|---|---|---|---|---|---|---|---|
| Tested? | TRUE | TRUE | TRUE | FALSE | TRUE | TRUE | TRUE | TRUE |
| MTERF3 | ChIP | GHT | HT | SMS | PBM | Lysate | eGFP-IVT | IVT |
| Tested? | FALSE | TRUE | TRUE | FALSE | TRUE | FALSE | TRUE | TRUE |
| MTERF4 | ChIP | GHT | HT | SMS | PBM | Lysate | eGFP-IVT | IVT |
| Tested? | TRUE | TRUE | TRUE | FALSE | TRUE | TRUE | TRUE | TRUE |

**TERB1 (Myb/SANT) is a potential TF.** This protein has a Myb/SANT domain, which is the main source of evidence for a role as a TF. It contains only a single Myb domain (with slightly less similarity to a SANT domain model). This domain has been shown to be involved in protein-protein interactions[54], but this may not be mutually exclusive with DNA-binding.
**Codebook experiment summary**: All experiment types were performed but none of them yielded specific PWMs. TERB1

| TERB1 | ChIP | GHT | HT | SMS | PBM | Lysate | eGFP-IVT | IVT |
|---|---|---|---|---|---|---|---|---|
| Tested? | TRUE | TRUE | TRUE | TRUE | TRUE | TRUE | TRUE | TRUE |

**NFX1 and NFXL1 (NFX) are potential TFs.** NFX1 has been indicated to be a TF, based on EMSA experiments performed with recombinant *E. coli*-expressed protein[55] and, for NFXL1, there are recent PWMs derived from ENCODE ChIP-seq (available in Factorbook database). One of the Factorbook motifs is selected in this study as a representative PWM for NFXL1 (**Supplementary Table 13** and **Supplementary Data 1**). The two datasets are discrepant, however. Based on the old EMSA study, NFX1 binds to DNA fragment with a sequence: 5' CCCTTCCCCTAGCAAGAGATG 3', (underscored part is the genomic site predicted by the authors), whereas some of the ChIP-seq-derived PWMs for NFXL1 are somewhat similar to PAX sites and others are low complexity G- and A-rich sequences. We selected one of the PAX-like PWMs as the representative PWM (Consensus: "CYRARGTCACASAGC").
Additional evidence supporting NFX domains as DBDs is an HT-SELEX experiment in *E. coli* that derived PWM for a fruit fly ortholog, *stc* (shuttlecraft), with very different consensus sequence "TATCAYAWKRTGATA"[56]. This discrepancy is expected,

however, as both the number of NFX domains (11 in NFXL1 and 7 in *stc*) and their sequences have diverged substantially.
On the other hand, there is evidence that links these proteins to RNA-associated functions[57], and both NFX1 and NFXL1 are associated very clearly with ribosomes based on protein-protein interactions listed in STRING database, casting some doubt on whether these proteins are TFs with inherent specificity.
**Codebook experiment summary**: All experiment types were performed for NFX1 and all but ChIP-seq for NFXL1. None of them yielded specific PWMs. NFX1, NFXL1

| NFX1 | ChIP | GHT | HT | SMS | PBM | Lysate | eGFP-IVT | IVT |
|---|---|---|---|---|---|---|---|---|
| Tested? | TRUE | TRUE | TRUE | TRUE | TRUE | TRUE | TRUE | TRUE |
| NFXL1 | ChIP | GHT | HT | SMS | PBM | Lysate | eGFP-IVT | IVT |
| Tested? | FALSE | TRUE | TRUE | TRUE | TRUE | FALSE | TRUE | TRUE |

**SP110 (SAND) is a potential TF**. This protein has a SAND domain. Many SAND domain TFs are supported by modern systematic datasets. Mouse Sp110 bound DNA in PBM experiments, but the human and mouse proteins are diverged.
**Codebook experiment summary**: ChIP-seq did not pass quality control, and none of the other experiments enriched in motifs. SP110

| SP110 | ChIP | GHT | HT | SMS | PBM | Lysate | eGFP-IVT | IVT |
|---|---|---|---|---|---|---|---|---|
| Tested? | TRUE | TRUE | TRUE | TRUE | TRUE | TRUE | TRUE | TRUE |

**TBPL1 (TBP) is a potential TF.** This protein has a TBP domain. The Codebook ChIP-seq was enriched in typical promoter motifs, suggesting that this protein functions in a similar fashion as its paralog TBP. It is a relatively distant paralog of TBP, and the only member of the TBP family that lacks the ability to bind the TATA box[58]. *In vitro*, purified TLF-TFIIA binds directly to the NF1 promoter[59], so it may be an obligate heteromer.
**Codebook experiment summary**: No SMS experiment. ChIP-seq is enriched in promoter-specific signals including YY1 and NRF1. None of the other assays enriched any motifs. TBPL1

| TBPL1 | ChIP | GHT | HT | SMS | PBM | Lysate | eGFP-IVT | IVT |
|---|---|---|---|---|---|---|---|---|
| Tested? | TRUE | TRUE | TRUE | FALSE | TRUE | TRUE | TRUE | TRUE |

### Proteins with unknown DBDs that are unlikely to be sequence specific TFs

**CENPA (Unknown) is not likely to be a TF.** The main source of evidence for a role as a TF was PDB:3AN2. This protein is a histone H3 variant protein that binds same way as H3 in a regular histone, based on crystal structure PDB:3AN2.
**Codebook experiment summary:** No SMS experiment. All other experiment types were performed but without specific PWM enrichment. CENPA

| CENPA | ChIP | GHT | HT | SMS | PBM | Lysate | eGFP-IVT | IVT |
|---|---|---|---|---|---|---|---|---|
| Tested? | TRUE | TRUE | TRUE | FALSE | TRUE | TRUE | TRUE | TRUE |

**AEBP1 (Unknown) is not likely to be a TF.** This protein was not annotated to have any likely DBDs. It has been suggested to moonlight as a TF, and to bind DNA based on EMSA[60] and ChIP[61]. It has a well characterized function as a protease (carboxypeptidase), however.
**Codebook experiment summary**: No SMS experiment. ChIP-seq did not pass quality control. All other experiment types were performed, but none of them yielded specific PWMs. AEBP1

| AEBP1 | ChIP | GHT | HT | SMS | PBM | Lysate | eGFP-IVT | IVT |
|---|---|---|---|---|---|---|---|---|
| Tested? | TRUE | TRUE | TRUE | FALSE | TRUE | TRUE | TRUE | TRUE |

**ARHGAP35 (Unknown) is not likely to be a TF**. This protein was not annotated to have any likely DBDs. It has been indicated to bind DNA specifically based on EMSA and nuclease protection assay data[62]. It has a well characterized function as A GTPase-activating protein (GAP) functioning in RhoGAP signaling, however[63]. Its interactions with Rho and Rac GTPases and other cytoskeleton components are also well supported by systematic analyses on protein-protein interaction data in STRING database.
**Codebook experiment summary**: No SMS experiment. ChIP-seq did not pass quality control. All other experiment types were performed, but none of them yielded specific PWMs. ARHGAP35

| ARHGAP35 | ChIP | GHT | HT | SMS | PBM | Lysate | eGFP-IVT | IVT |
|---|---|---|---|---|---|---|---|---|
| Tested? | TRUE | TRUE | TRUE | FALSE | TRUE | TRUE | TRUE | TRUE |

**CC2D1A (Unknown) is not likely to be a TF.** This protein was not annotated to have any likely DBDs. It has been shown to bind DNA specifically on recombinant protein based EMSA and nuclease protection assay data[64]. However, it makes strong interactions with endosomal transport associated proteins

**Codebook experiment summary**: No IVT SELEX, SMS or PBM experiments. ChIP-seq did not pass quality control. All other experiment types were performed but none of them yielded specific PWMs. CC2D1A

| CC2D1A | ChIP | GHT | HT | SMS | PBM | Lysate | eGFP-IVT | IVT |
|---|---|---|---|---|---|---|---|---|
| Tested? | TRUE | TRUE | TRUE | FALSE | FALSE | TRUE | TRUE | FALSE |

**CENPT (Unknown) is a centromere associated protein complex component and not likely to be a TF.** This protein was not annotated to have any likely sequence specific DNA-binding domains. It contains an archaeal histone-like fold (CBF_NFY), however, and has a known role in centromere assembly. It makes DNA backbone contacts and potentially has a minor groove inserting arginine in cryo-EM structure 7R5S[65]. In the structure it is intertwined together with CENPW and can likely bind DNA with it as a heterodimer, based on DNA contacts the binding is likely to have low specificity.
**Codebook experiment summary**: No SMS experiments. ChIP-seq did not pass quality control. All other experiment types were performed but none of them yielded specific PWMs. CENPT

| CENPT | ChIP | GHT | HT | SMS | PBM | Lysate | eGFP-IVT | IVT |
|---|---|---|---|---|---|---|---|---|
| Tested? | TRUE | TRUE | TRUE | FALSE | TRUE | TRUE | TRUE | TRUE |

**CHCHD3 (Unknown) is not likely to be a TF.** This protein was not annotated to have any likely DBDs. Sequence specific binding activity has been indicated by EMSA utilizing recombinant protein produced in *E. coli*, showing that it can specially bind the hBAG-1 promoter[66]. However, the protein is a known mitochondrial membrane scaffold protein.
**Codebook experiment summary**: No SMS experiment. ChIP-seq did not pass quality control. All other experiment types were performed but none of them yielded specific PWMs. CHCHD3

| CHCHD3 | ChIP | GHT | HT | SMS | PBM | Lysate | eGFP-IVT | IVT |
|---|---|---|---|---|---|---|---|---|
| Tested? | TRUE | TRUE | TRUE | FALSE | TRUE | TRUE | TRUE | TRUE |

**CSRNP1, CSRNP2, and CSRNP3 (Unknown) may be transcriptional cofactors.** These proteins were not annotated to have any likely DBDs. All three paralogs (CSRNP1, CSRNP2, and CSRNP3) were reported to bind an “AGAGTG” consensus in SELEX experiments performed with recombinant HEK293 expressed proteins[67]. However, the study has insufficient details about the SELEX experiments for evaluation of its results, including entire sequences of the 30-bp DNA fragments. Based on protein-protein interactions and mostly nuclear localization, these proteins could be cofactors.

**Codebook experiment summary**: All experiment types were performed but none of them yielded specific PWMs. CSRNP1, CSRNP2, CSRNP3

| CSRNP1 | ChIP | GHT | HT | SMS | PBM | Lysate | eGFP-IVT | IVT |
|---|---|---|---|---|---|---|---|---|
| Tested? | TRUE | TRUE | TRUE | TRUE | TRUE | TRUE | TRUE | TRUE |
| CSRNP2 | ChIP | GHT | HT | SMS | PBM | Lysate | eGFP-IVT | IVT |
| Tested? | TRUE | TRUE | TRUE | TRUE | TRUE | TRUE | TRUE | TRUE |
| CSRNP3 | ChIP | GHT | HT | SMS | PBM | Lysate | eGFP-IVT | IVT |
| Tested? | TRUE | TRUE | TRUE | TRUE | TRUE | TRUE | TRUE | TRUE |

**GLMP (Unknown) is not likely to be a TF or a cofactor.** This protein was not annotated to have any likely DBDs. Sequence specificity is indicated by an EMSA experiment with a recombinant protein, showing that it binds GATCCGCCCGCTTGTGGCCAACTGGCTCCAGTCAC[68]. It is a known lysosomal membrane protein, however.
**Codebook experiment summary**: No SMS experiment. ChIP-seq did not pass quality control. All other experiment types were performed but none of them yielded specific PWMs. GLMP

| GLMP | ChIP | GHT | HT | SMS | PBM | Lysate | eGFP-IVT | IVT |
|---|---|---|---|---|---|---|---|---|
| Tested? | TRUE | TRUE | TRUE | FALSE | TRUE | TRUE | TRUE | TRUE |

**GPBP1 and GPBP1L1 (Unknown) are unlikely to be TFs.** These proteins were not annotated to have any likely DBDs. The main source of evidence for a role of GPBP1 (also known as Vasculin) as a TF is a single study with a recombinant protein based EMSA showing that it binds DNA[69], however another study shows that it has an "extended AT-hook" DNA-binding peptide motif that has stronger affinity towards RNA than DNA[70]. GPBP1L1 is a close paralog of GPBP1, but very little is known of it. GPBP1 and GPBP1L1 form protein–protein interactions mostly with RNA processing connected proteins.
**Codebook experiment summary**: No SMS experiment. ChIP-seq did not pass quality control. All other experiment types were performed but none of them yielded specific PWMs. GPBP1, GPBP1L1

| GPBP1 | ChIP | GHT | HT | SMS | PBM | Lysate | eGFP-IVT | IVT |
|---|---|---|---|---|---|---|---|---|
| Tested? | TRUE | TRUE | TRUE | FALSE | TRUE | TRUE | TRUE | TRUE |
| GPBP1L1 | ChIP | GHT | HT | SMS | PBM | Lysate | eGFP-IVT | IVT |
| Tested? | TRUE | TRUE | TRUE | FALSE | TRUE | TRUE | TRUE | TRUE |

**KCNIP3 (Unknown) is not likely to be a TF**. This protein was not annotated to have any likely DBDs. Sequence specific DNA binding activity was shown by recombinant protein expression based EMSA experiment[71]. However, the protein has a well-established role in binding to and controlling specific types of potassium channels. This protein has been researched extensively but there is very little further evidence of sequence specific DNA binding activity.
**Codebook experiment summary**: No SMS experiments. All other experiment types were performed but none of them yielded specific PWMs. KCNIP3

| KCNIP3 | ChIP | GHT | HT | SMS | PBM | Lysate | eGFP-IVT | IVT |
|---|---|---|---|---|---|---|---|---|
| Tested? | TRUE | TRUE | TRUE | FALSE | TRUE | TRUE | TRUE | TRUE |

**NKRF (Unknown) is not likely to be a TF**. This protein was not annotated to have any likely DBDs. It is known to regulate NFKB activity, indicating a role in transcriptional regulation. EMSA experiments performed with induced nuclear extracts have indicated binding to AATTTCCTCTGA[72] and to AATTCCCTGA[73]. A recent study, however, indicates that NKRF binds NFKB directly to inhibit its activity[74]. Protein interaction data in STRINGdb suggest a role in RNA editing.
**Codebook experiment summary**: ChIP-seq did not pass quality control. All other experiment types were performed but none of them yielded specific PWMs. NKRF

| NKRF | ChIP | GHT | HT | SMS | PBM | Lysate | eGFP-IVT | IVT |
|---|---|---|---|---|---|---|---|---|
| Tested? | TRUE | TRUE | TRUE | TRUE | TRUE | TRUE | TRUE | TRUE |

**NME2 (Unknown) is not likely to be a TF**. This protein was not annotated to have any likely DBDs. One study indicated that it binds to specific sequences in nuclear extract based EMSA[75], and another that it associated with telomeres based on ChIP-seq[76]. The protein is an enzyme, however (nucleoside diphosphate kinase) and the DNA in the structure PDB:3BBB[77] is a single-stranded dinucleotide. Taken together, and with the negative Codebook results, the evidence suggests that it is not a sequence specific TF.
**Codebook experiment summary**: No ChIP-seq experiment. All other experiments performed but without specific PWM enrichment. NME2

| NME2 | ChIP | GHT | HT | SMS | PBM | Lysate | eGFP-IVT | IVT |
|---|---|---|---|---|---|---|---|---|
| Tested? | FALSE | TRUE | TRUE | TRUE | TRUE | TRUE | TRUE | TRUE |

**PA2G4 (Unknown) is likely a low specificity DNA-binding cofactor**. This protein was not annotated to have any likely DBDs. It binds CGGCAAAAAGG repeats, based on EMSA performed with recombinant protein[78]. Many other lines of evidence including cryo-EM structure **PDB: 6SXO** show that it is a double-stranded ribosomal RNA binding protein and a co-regulator[79,80].

**Codebook experiment summary**: No SMS experiment. ChIP-seq did not pass quality control. All other experiment types were performed but none of them yielded specific PWMs. PA2G4

| [81]G4 | ChIP | GHT | HT | SMS | PBM | Lysate | eGFP-IVT | IVT |
|---|---|---|---|---|---|---|---|---|
| Tested? | TRUE | TRUE | TRUE | FALSE | TRUE | TRUE | TRUE | TRUE |

**PCGF2 and PCGF6 (Unknown) are likely to be transcriptional cofactors.** These paralogs were not annotated to have any likely DBDs, as RING type zinc finger domain in them is a protein-protein interaction interphase used to contact other RING domains. A study from 1995 study showed that PCGF2 binds “GACTNGACT“ based on SELEX and EMSA performed with cell expressed recombinant proteins, and that binding can be achieved also in EMSA performed with *E. coli* expressed protein[81]. Substantial amount of research has since shown alternative recruitment strategies for these protein complexes[82]. The PCGF2 and PCGF6 proteins function in a PRC1.2 and PRC1.6 polycomb complexes, respectively. PRC1.2 has been shown to be recruited mainly through CBX protein mediated interactions where the chromodomain of the CBX contacts H3K27me3 to silence chromatin. PRC1.6, on the other hand, has been shown to be recruited by heterodimeric combinations of MAX+MNT (bHLH) and E2F6+DP1 (E2F), and concordantly, our ChIP-seq experiments show enrichement of “CACGTG”- and “GGCGGGAAA”-containing PWMs[82].
**Codebook experiment summary**: No SMS experiment for either. ChIP-seq experiments for PCGF2 did not pass quality control, whereas PCGF6 enriched in “CACGTG”-sequences bound by many bHLH TFs and CG-rich sequences. None of the other experiment types yielded specific PWMs. PCGF2, PCGF6

| PCGF2 | ChIP | GHT | HT | SMS | PBM | Lysate | eGFP-IVT | IVT |
|---|---|---|---|---|---|---|---|---|
| Tested? | TRUE | TRUE | TRUE | FALSE | TRUE | TRUE | TRUE | TRUE |
| PCGF6 | ChIP | GHT | HT | SMS | PBM | Lysate | eGFP-IVT | IVT |
| Tested? | TRUE | TRUE | TRUE | FALSE | TRUE | TRUE | TRUE | TRUE |

**PLSCR1 (Unknown) is not likely to be a TF.** This protein was not annotated to have any likely DBDs. Recombinant protein based EMSA evidence indicates that this enzyme can bind to a “GTAACCATGTGGA” sequence that is present in the IP3R1 promoter[83]. Protein has been researched very extensively[84], it is best characterized as membrane bound enzyme (phospholipid scramblase), that localizes mostly to plasma membrane but is likely actively transported to intracellular compartments as response to various stimuli. In nucleus it has been shown to associate with several transcription factors and through them to different promoters[84] Based on these lines of evidence, and that it did not enrich specific PWMs in any of our analyses it functions most likely as a cofactor.
**Codebook experiment summary**: No SMS experiment and ChIP-seq did not pass quality control. No other experiment types enriched specific PWMs. PLSCR1

| PLSCR1 | ChIP | GHT | HT | SMS | PBM | Lysate | eGFP-IVT | IVT |
|---|---|---|---|---|---|---|---|---|
| Tested? | TRUE | TRUE | TRUE | FALSE | TRUE | TRUE | TRUE | TRUE |

**PREB (Unknown) is not likely to be a TF.** This protein was not annotated to have any likely DBDs. EMSA experiments indicated that it binds DNA[85]; however, many other lines of evidence link it to functioning in ER-Golgi transport as a "guanine nucleotide exchange factor" (GEF) that regulates the assembly of the coat protein complex II/COPII in endoplasmic reticulum (ER) to Golgi vesicle-mediated transport.
**Codebook experiment summary**: No SMS experiment. ChIP-seq did not pass quality control. All other experiment types were performed but none of them yielded specific PWMs. PREB

| PREB | ChIP | GHT | HT | SMS | PBM | Lysate | eGFP-IVT | IVT |
|---|---|---|---|---|---|---|---|---|
| Tested? | TRUE | TRUE | TRUE | FALSE | TRUE | TRUE | TRUE | TRUE |

**RBCK1 (Unknown) is not likely to be a TF.** This protein was not annotated to have any likely DBDs, and it has a well-established ubiquitin ligase. EMSA and SELEX indicated that it bound TGG trinucleotide repeats *in vitro*[86]. To our knowledge, there have been no other reports regarding its sequence specificity or function as a TF since 1998. Thus, altogether, it has poorly defined DNA sequence/structure specificity and limited support as a TF.
**Codebook experiment summary**: No SMS or PBM experiments. ChIP-seq did not pass quality control, and other experiment types did not enrich specific PWMs. RBCK1

| RBCK1 | ChIP | GHT | HT | SMS | PBM | Lysate | eGFP-IVT | IVT |
|---|---|---|---|---|---|---|---|---|
| Tested? | TRUE | TRUE | TRUE | FALSE | FALSE | TRUE | TRUE | TRUE |

**REXO4 (Unknown) is not likely to be a TF.** This protein does not have any apparent DBDs, and it is a DNA/RNA exonuclease. An EMSA-based study indicated that it can enhance the ability of ESR2 to bind its targets[87].
**Codebook experiment summary**: Only IVT based SELEX experiment. No specific PWM enrichment. REXO4

| REXO4 | ChIP | GHT | HT | SMS | PBM | Lysate | eGFP-IVT | IVT |
|---|---|---|---|---|---|---|---|---|
| Tested? | FALSE | TRUE | TRUE | FALSE | FALSE | FALSE | FALSE | TRUE |

**SAFB and SAFB2 (Unknown) are not likely to be TFs.** These proteins do not have likely DBDs, and many studies have linked them to binding RNA. Existing evidence for them being TFs are filter binding assays[88].
**Codebook experiment summary**: No ChIP-seq, Lysate SELEX or SMS experiments. Other experiments were performed but without specific PWM enrichment. SAFB, SAFB2

| SAFB | ChIP | GHT | HT | SMS | PBM | Lysate | eGFP-IVT | IVT |
|---|---|---|---|---|---|---|---|---|
| Tested? | FALSE | TRUE | TRUE | FALSE | TRUE | FALSE | TRUE | TRUE |
| SAFB2 | ChIP | GHT | HT | SMS | PBM | Lysate | eGFP-IVT | IVT |
| Tested? | TRUE | TRUE | TRUE | TRUE | TRUE | TRUE | TRUE | TRUE |

**SCMH1 (Unknown) is likely a transcriptional cofactor and not a TF.** This protein does not have likely DBDs. It was included due to homology with SCML2, a polycomb repressive complex protein. It associates with polycomb complexes and is likely a cofactor.
**Codebook experiment summary**: No SMS experiment. ChIP-seq did not pass quality control. All other experiment types were performed but none of them yielded specific PWMs. SCMH1

| SCMH1 | ChIP | GHT | HT | SMS | PBM | Lysate | eGFP-IVT | IVT |
|---|---|---|---|---|---|---|---|---|
| Tested? | TRUE | TRUE | TRUE | FALSE | TRUE | TRUE | TRUE | TRUE |

**SMYD3 (Unknown) is likely a transcriptional cofactor, and not a TF.** This protein was not annotated to have any likely DBDs. It is a histone methyltransferase. One report showed that it binds GGAGGG elements, based on SELEX and EMSA performed using GST fusion protein[89].
**Codebook experiment summary**: ChIP-seq did not pass quality control. All other experiment types were performed but none of them yielded specific PWMs. SMYD3

| SMYD3 | ChIP | GHT | HT | SMS | PBM | Lysate | eGFP-IVT | IVT |
|---|---|---|---|---|---|---|---|---|
| Tested? | TRUE | TRUE | TRUE | TRUE | TRUE | TRUE | TRUE | TRUE |

**SNAPC2 and SNAPC5 (Unknown) are unlikely to be TFs.** These proteins have been not annotated to have any likely DBDs and. SNAPC2 was included due to cellular extract based EMSA and antibody supershift evidence, which show that it is a part of initiation complex, but there is no evidence suggesting that it binds DNA on its own[90]. For SNAPC5, an EMSA experiment was performed with cell extract[91]. The protein is present in several initiation complex structures[92] (PDB: 7ZWC, 7ZWD or 7ZX7) but does not contact DNA directly in any of them.

**Codebook experiment summary**: All experiment types were performed but none of them yielded specific PWMs. SNAPC2, SNAPC5

| SNAPC2 | ChIP | GHT | HT | SMS | PBM | Lysate | eGFP-IVT | IVT |
|---|---|---|---|---|---|---|---|---|
| Tested? | TRUE | TRUE | TRUE | TRUE | TRUE | TRUE | TRUE | TRUE |
| SNAPC5 | ChIP | GHT | HT | SMS | PBM | Lysate | eGFP-IVT | IVT |
| Tested? | FALSE | TRUE | TRUE | TRUE | TRUE | TRUE | TRUE | TRUE |

**SON (Unknown) is not likely to be a TF**. This protein was not annotated to have any likely DBDs. The main source of evidence for a role as a TF was study in which SELEX and EMSA were carried out with transfected cell extracts[93], which indicates that it (or something it associates with) binds “GAKANSRCC”. SON is thought to be mainly an RNA-binding protein that acts as a mRNA splicing cofactor, however.
**Codebook experiment summary**: Only ChIP-seq and Lysate based SELEX were performed. ChIP-seq did not pass quality control. No specific PWM enrichment. SON

| SON | ChIP | GHT | HT | SMS | PBM | Lysate | eGFP-IVT | IVT |
|---|---|---|---|---|---|---|---|---|
| Tested? | TRUE | TRUE | TRUE | FALSE | FALSE | TRUE | FALSE | FALSE |

**TET2 (Unknown) is likely a low specificity DNA-binding cofactor**. This protein was not annotated to have any likely DBDs, however, it is an enzyme that oxidises methylated CpG sequences to hydroxymethyls. It was included based on crystal structure PDB:4NM6[94] that shows the enzymatic domain of the protein complexed with a CpG methylated DNA. Protein binds DNA on the minor groove side, making backbone contacts and showing that the methylated base is flipped out of double-stranded context to face toward the catalytic core of the enzyme. Based on many lines of evidence, the removal of CpG methylation is guided with multiple mechanisms, including both sequence specific TFs and chromatin modifications, making it unlikely that there is significant contribution of intrinsic specificity[95].
**Codebook experiment summary**: ChIP-seq enriched specific PWMs, most prominent of which were for posterior homeodomains, whereas all other assays were unsuccessful. TET2

| TET2 | ChIP | GHT | HT | SMS | PBM | Lysate | eGFP-IVT | IVT |
|---|---|---|---|---|---|---|---|---|
| Tested? | TRUE | TRUE | TRUE | TRUE | TRUE | TRUE | TRUE | TRUE |

**THYN1 (Unknown) is a low specificity DNA-binding protein.** This protein was included in the study based on X-ray crystallography structure (**PDB:5J3E,** structure is not connected to a publication) with CpG methylated palindromic DNA sequence “GCCAAmCGTTGGC”. In this structure, the protein makes extensive contacts with the

DNA backbone, but contacts bases only through a single arginine inserted into the minor groove. In the structure, the protein does not contact the methylated cytosine. These features of the structure and the fact that protein did not enrich any motifs in experiments suggest that its DNA binding is nonspecific.
**Codebook experiment summary**: All experiment types were performed but none of them yielded specific PWMs. THYN1

| THYN1 | ChIP | GHT | HT | SMS | PBM | Lysate | eGFP-IVT | IVT |
|---|---|---|---|---|---|---|---|---|
| Tested? | TRUE | TRUE | TRUE | TRUE | TRUE | TRUE | TRUE | TRUE |

**TMF1 (Unknown) is likely not a TF.** This protein has not been annotated to have any likely DBDs and based on AF3 prediction, it is composed entirely of long alpha helices and unstructured regions. The protein was reported to bind DNA specifically based on EMSA experiments, where bacterially expressed recombinant protein bound specifically to the TATA element[96]. A second transcriptional regulation connecting study from 1999 described it as a co-activator of androgen receptor (AR), based on it interacting well with AR and up-regulating the expression of AR targets[97]. Later on, in a study from 2007 the protein was indicated to function in Rab6 mediated retrograde transport on the levels of both, endosomes to Golgi and Golgi to ER[98]. Interactions between TMF1, AR and Golgi transport proteins are validated by systematic analyzes of protein-protein interactions, and thus the protein's role in retrograde transport activity could potentially explain AR target upregulation seen in 1999 study. To our knowledge, however, no studies since the original EMSA-experiments have shown direct DNA-binding activity to this protein, and together with negative results in all our assays and a predicted structure that appears incompatible with DNA binding activity, it seems unlikely that the protein is a TF.
**Codebook experiment summary**: All experiment types were performed but none of them yielded specific PWMs. TMF1

| TMF1 | ChIP | GHT | HT | SMS | PBM | Lysate | eGFP-IVT | IVT |
|---|---|---|---|---|---|---|---|---|
| Tested? | TRUE | TRUE | TRUE | TRUE | TRUE | TRUE | TRUE | TRUE |

**TSC22D1 (Unknown) is likely a transcriptional cofactor.** This protein does not have any likely DBDs as it contains a potential leucine zipper, but without the basic region used to bind DNA in bZIP proteins. It was included based on an EMSA experiments by Ohta et al. 1996, performed with *E. coli* expressed recombinant protein that indicated it to bind CG-rich sequences[99]. TSC22D1 interacts with known TFs, including SMAD family members TFs to affect transcription, however[100,101], and to our knowledge there has been no evidence of direct DNA binding activity.
**Codebook experiment summary**: Only ChIP-seq and Lysate based SELEX experiments. ChIP-seq did not pass quality control. No enrichment for specific PWMs. TSC22D1

| TSC22D1 | ChIP | GHT | HT | SMS | PBM | Lysate | eGFP-IVT | IVT |
|---|---|---|---|---|---|---|---|---|
| Tested? | TRUE | TRUE | TRUE | FALSE | FALSE | TRUE | FALSE | FALSE |

### Potential TFs with unknown DNA-binding domain

**RAG1 (Unknown) is likely a sequence specific TF.** The main evidence for sequence specific DNA binding activity of RAG1 is X-ray structures showing the protein binding to DNA by itself (PDB: 3GNA) and also as a heteromultimer with RAG2 3(JBW)[102]. It binds DNA specifically as a part of sequence specific nuclease complex that works in V(D)J recombination, but as a sequence specific DNA binder it could also have regulatory activity.
**Codebook experiment summary**: No SMS experiment. Codebook ChIP-seq data are enriched in a "ACAAAAACC" motif that is concordant with DNA sequence in crystal structure PDB: 3GNA. Other experiments were performed but without specific PWM enrichment. RAG1

| RAG1 | ChIP | GHT | HT | SMS | PBM | Lysate | eGFP-IVT | IVT |
|---|---|---|---|---|---|---|---|---|
| Tested? | TRUE | TRUE | TRUE | FALSE | TRUE | TRUE | TRUE | TRUE |

**CENPS and CENPX (Unknown) are likely sequence-specific DNA-binding proteins that bind together as a cooperative complex.** Their potential for sequence specific DNA binding activity was based on crystal structure **PDB:4NDY[103]**, which shows CENPS and CENPX binding DNA as a complex that is consistent of them being obligate heterodimerization partners of each other, as the protein chains are intertwined. DNA in the structure is an AAAAAAA-stretch.
**Codebook experiment summary**: No SMS experiment. ChIP-seq experiments did not pass quality control and *in vitro* experiments were tested only in individual contexts. None of the experiments enriched specific PWMs. CENPS, CENPX

| CENPS | ChIP | GHT | HT | SMS | PBM | Lysate | eGFP-IVT | IVT |
|---|---|---|---|---|---|---|---|---|
| Tested? | TRUE | TRUE | TRUE | FALSE | TRUE | TRUE | TRUE | TRUE |
| CENPX | ChIP | GHT | HT | SMS | PBM | Lysate | eGFP-IVT | IVT |
| Tested? | TRUE | TRUE | TRUE | FALSE | TRUE | TRUE | TRUE | TRUE |

**SKI and SKIL (Unknown) are likely sequence specific TFs**. These proteins have a SKI/SNO/DACH DNA binding domain (See **Supplementary Information** for more details). The main evidence of sequence specific DNA binding activity is a study (Nicol et al. 1998) reporting that, by SELEX and EMSA, SKI (produced from cell extract) binds a "GTCTAGAC" consensus[104].
**Codebook experiment summary**: ChIP-seq experiments did not pass quality control. The same consensus sequence described by Nicol et al. clearly enriches in our lysate-based HT-SELEX for SKIL, however. This site enriched only in one weak experiment, and therefore the PWM did not pass our quality control. Nonetheless, it clearly validates the sequence specific DNA-binding activity of the protein. Both our SKIL experiment and

the Nicol et, al. data are based on cellular extracts, and as these proteins were unsuccessful with other protein production methods, it is possible that they require a post-transcriptional modification, bind as heterodimers, or potentially contact DNA indirectly. SKI, SKIL

| SKI | ChIP | GHT | HT | SMS | PBM | Lysate | eGFP-IVT | IVT |
|---|---|---|---|---|---|---|---|---|
| Tested? | TRUE | TRUE | TRUE | FALSE | FALSE | FALSE | TRUE | FALSE |
| SKIL | ChIP | GHT | HT | SMS | PBM | Lysate | eGFP-IVT | IVT |
| Tested? | TRUE | TRUE | TRUE | FALSE | FALSE | TRUE | TRUE | FALSE |

**DRAP1 (Unknown) is a potential TF.** This protein was not annotated to have any likely DBDs. DRAP1 produced AT-rich motif in external ChIP-seq that is consistent with its known function of binding DNA as cooperative complex of DR1, DRAP1 and TBP in structure PDB: 1JFI[105]
**Codebook experiment summary**: No SMS experiment. ChIP-seq is enriched in NFY-like PWMs. Other experiment types did not yield clear enrichment. DRAP1

| DRAP1 | ChIP | GHT | HT | SMS | PBM | Lysate | eGFP-IVT | IVT |
|---|---|---|---|---|---|---|---|---|
| Tested? | TRUE | TRUE | TRUE | FALSE | TRUE | TRUE | TRUE | TRUE |

**BRF2 (Unknown) is likely a sequence specific TF.** This protein was not annotated to have any likely DBDs. Based on PDB:4ROC it binds cooperatively with TBP and defines flanking specificity[106]; proper interaction needs the DNA bending caused by TBP and it is likely an obligate heterodimer.
**Codebook experiment summary**: No SMS experiment. ChIP-seq did not pass quality control. All other experiment types were performed but none of them yielded specific PWMs. BRF2

| BRF2 | ChIP | GHT | HT | SMS | PBM | Lysate | eGFP-IVT | IVT |
|---|---|---|---|---|---|---|---|---|
| Tested? | TRUE | TRUE | TRUE | FALSE | TRUE | TRUE | TRUE | TRUE |

**DR1 (Unknown) is a potential TF.** This protein was not annotated to have any likely DBDs. The main source of evidence for a role as a TF is crystal structure PDB:1JFI[105], showing it as an obligate heterodimer with DRAP1; together they bind TBP to inhibit its function. DR1 and DRAP1 make interactions with DNA but binding could be dependent on TBP1. DRAP1 produced AT-rich motif in external ChIP-seq that is consistent with this function
**Codebook experiment summary**: No SMS experiment. Our ChIP-seq shows enrichment of “TATAAA” motif consistently with it binding together with TBP1 (and possibly DRAP1). No other experiment types enrich in clear motifs. DR1

| DR1 | ChIP | GHT | HT | SMS | PBM | Lysate | eGFP-IVT | IVT |
|---|---|---|---|---|---|---|---|---|
| Tested? | TRUE | TRUE | TRUE | FALSE | TRUE | TRUE | TRUE | TRUE |

**PHF19 (Unknown) is likely a sequence specific TF.** This protein was not annotated to have any likely DBDs. A recent study shows with PBM that it bound a CG-rich PWM and also that its homologs PHF1 and MTF2 bound DNA specifically with a WH domain[107]. This protein is also known to bind trimethylated Lys-36 of histone H3.
**Codebook experiment summary**: No eGFP-IVT based SELEX or SMS experiments. ChIP-seq is enriched in CG-rich sequences All other experiment types were performed but none of them yielded specific PWMs. PHF19

| PHF19 | ChIP | GHT | HT | SMS | PBM | Lysate | eGFP-IVT | IVT |
|---|---|---|---|---|---|---|---|---|
| Tested? | TRUE | TRUE | TRUE | FALSE | TRUE | TRUE | FALSE | TRUE |

**PURG (Unknown) is a potential TF.** This protein was not annotated to have any likely DBDs. Its paralog PURB was successful with PBM, and paralog PURA has a Transfac PWM. An existing cell extract EMSA based results indicated binding to GGA repeats[108], but this is discrepant with our PBM data for paralog PURB. These proteins have been also linked on binding single stranded DNA[109].
**Codebook experiment summary**: No Lysate and eGFP-IVT based SELEX or SMS experiments. ChIP-seq did not pass quality control. Other experiment types did not enriched in anything. PURG

| PURG | ChIP | GHT | HT | SMS | PBM | Lysate | eGFP-IVT | IVT |
|---|---|---|---|---|---|---|---|---|
| Tested? | TRUE | TRUE | TRUE | FALSE | TRUE | FALSE | FALSE | TRUE |
