## Supplementary Discussion & Methods for "Codebook: sequence specificity and genomic binding of poorly-characterized human transcription factors"

#### Under-characterized DBDs, including the SKI/SNO/DAC and C-Clamp domains

##### Introduction

Six Codebook proteins that lacked canonical DBDs yielded successful experiments, and therefore motifs (CGGBP1, NACC2, TCF20, PURB, DACH1, and DACH2). All six appear to represent DBDs that were poorly described at the outset of the Codebook study. We and others have recently described CGGBP1 as the founding member of an extensive family of eukaryotic TFs derived from the DBDs of transposons, which may be related to the BED-type Zinc Finger found in ZBED TFs[62,108]. NACC2 contains a BEN domain, which over the last decade has been established as typically a sequence-specific DBD[109-111]. TCF20 contains a potential AT-hook[112] (below the conventional Pfam scoring threshold) and yielded an AT-hook-like motif. PURB is composed largely of three copies of the PUR (Purine-rich-element binding) domain; it yielded a motif on four different PBM assays (resembling ACCnAC/GTnGGT), which is unlike its previously established binding site (CTTCCCTGGAAG)[113]. The sequence specificity of this protein thus remains enigmatic.

DACH1 and DACH2 are paralogs that yielded very similar motifs. They contain a SKI/SNO/DAC domain, a winged helix-turn-helix (HTH)-type DBD shared with their *Drosophila melanogaster* ortholog Dachshund. Members of this family have been previously described as binding to specific DNA sequences: a Forkhead-like motif (different from the one we obtained) was previously described for human DACH1[19], and human SKI was reported to bind GTCTAGAC in SELEX[114]. The SKI/SNO/DAC domain is also believed to possess other, non-DNA binding biochemical functions, however. For example, the SKI/SNO/DAC of *D. melanogaster* Dac is necessary for transcription activation in yeast two-hybrid assays, potentially implying a protein-protein interaction function[115]. Previous structural and biochemical analysis of the SKI/SNO/DAC domain of SKOR1/2 also suggests that in these proteins the domain has a protein-interaction function[21,116]

In addition to these six examples, the sequence specificity of SLC2A4RG and ZNF395 – both Codebook proteins with only a single C2H2-zf – resides in their C-clamp, rather than the C2H2-zf. The C-clamp is also present in ZNF704, which has a published HT-SELEX PWM that is virtually identical to that of SLC2A4RG and ZNF395 (resembling the sequence CCGGCCGG)[101]. The C-clamp domain is perhaps best known for its presence in two members of the TCF/LEF family in humans, in which it augments the DNA sequence specificity of the HMG domain[117]. The C-clamps in ZNF704, ZNF395, and SLC2A4RG have also been previously described as binding to shorter, unmethylated sequences containing mainly C and G[118,119], most typically “CCGG”. These sequences are also enriched in the Codebook datasets, but the apparent dimeric site “CCGGCCGG” is even more preferred in several different assays[14].

**SKI/SNO/DAC and C-clamp potentially represent expansive families of transcription factors**

***SKI/SNO/DAC.*** Both of the last two domain classes (SKI/SNO/DAC and C-clamp) are distributed broadly across animals, and thus may represent large classes of unexplored DBDs. We therefore sought to further characterize their phylogenetic distribution, the structural basis of their ability to bind DNA, and the DNA-binding specificities of putative transcription factors encoding these domains across animals.

Interpro[120] lists over 7,000 proteins containing a SKI/SNO/DAC domain, entirely in metazoans. DACH1 and DACH2 orthologs are prevalent across vertebrates, with many duplicates retained in salmonids due to their lineage-specific whole-genome duplication (**Extended Data Fig. 2a**). Phylogenetic analysis of SKI/SNO/DAC domains reveals four major groups. DACH (containing DACH1/2, and invertebrate Dac), SKOR (SKOR1/2 in vertebrates and *D. melanogaster* Fuss), and SKIDA1, which appears to be restricted to vertebrates, are clearly defined (**Extended Data Fig. 2b**). Vertebrate SKI and SKIL and *D. melanogaster* Snoo appear to be early diverging genes; more detailed analysis would be necessary to confidently state that they form a clade, but they are more similar to each other than to members of the other three groups.

We tested a representative set of 21 SKI/SNO/DAC domains across animals, as well as all seven human proteins encoding these domains, using HT-SELEX. DACH1 and DACH2 yielded the same motifs as previously. SKIDA1 also produced a clear DNA-binding motif (**Extended Data Fig. 2b**). In addition, while SKI failed in our follow-up assays, its paralog SKIL was among the original Codebook set, in which it produced a PWM in only a single HT-SELEX experiment, and thus did not pass our quality control. The PWM generated in this assay is, however, concordant with the SELEX-derived consensus sequence GTCTAGAC by Nicol et al. 1998 for SKI, from only 33 SELEX ligands[114]. Both our SKIL experiment and the Nicol et al. data are based on cellular extracts, and as these proteins were unsuccessful with other protein production methods, they may require a post-transcriptional modification, bind as heterodimers with an unknown protein, or potentially contact DNA indirectly. None of the other SKI/SNO/DAC family members yielded a clear positive result.

In support of direct DNA binding by at least a subset of the SKI/SNO/DAC class, Alphafold3[73] predicted that the DACH1 HTH inserts into the major groove precisely at the PWM-predicted binding site within an extended DNA sequence (**Extended Data Fig. 2c, left**). We also modeled full-length and SKI/SNO/DAC-only SKI and SKIL protein sequences with a sequence from Nicol et al 1998[114] in AlphaFold3, but the molecules were not predicted to interact. By contrast, the SKIDA1 SKI/SNO/DAC was predicted (albeit with a low iPTM = 0.26) to interact with a high-scoring HT-SELEX sequence at precisely its motif, with a similar configuration as DACH1 (**Extended Data Fig. 2c, right**).

***C-clamp.*** Interpro[120] lists over 8,000 proteins containing the SMART[121] C-clamp domain, entirely in metazoans and a handful of closely related protists. Ensembl has called one-to-one orthologs across most animals for all but one of the human C-clamp containing proteins (**Extended Data Fig. 2d**), again with whole-genome derived duplicates in salmonids across most of the genes. By contrast, SLC2A4RG appears to be a tetrapod-specific paralog, as synteny-supported one-to-one orthologs only exist through to amphibians in Ensembl[122]. Phylogenetic analysis of the C-clamp from human and representative non-human organisms clearly divides it into two major clades – the Glut4EF/ZNF395 family containing ZN704 and SLC2A4G, and the LEF/TCF family (**Extended Data Fig. 2e**).

Like SKI/SNO/DAC, we assayed a representative set of C-clamp domains on HT-SELEX, selected from diverse animals as well as all five human proteins encoding C-clamps. Of the 17 proteins tested, we successfully derived 9 high-quality C-clamp-like motifs (**Extended Data Fig. 2e**). The TCF-proteins were tested with both the C-clamp alone, the HMG alone, and the full-length protein including both domains.

The C-clamp domains of human TCF7 and TCF7L2 both failed when assayed alone. The full-length protein for TCF7L2, however, did derive both a clear HMG motif as well as a C-clamp-like motif (A/G)CCG, albeit at a lower enrichment (**Extended Data Fig. 2e**). We did not derive a compound motif containing the signatures of both DBDs, but the enrichment of the C-clamp motif is consistent with its involvement in binding "helper sites" with the consensus (A/G)CCG near HMG motifs, with the C-clamp's inclusion in some TCF isoforms modulating binding site usage downstream of Wnt signaling[117]. The failure to enrich a bipartite motif with static spacing may be due to flexibility in orientation and distance between the HMG and helper motifs[123]. The C-clamp domain alone from *C. elegans* Pop-1 (a TCF-like protein) binds a nearly identical motif to the C-clamp motif obtained from the longer TCF7L2 construct, however, consistent with the deep conservation of Wnt signaling across animals, and the importance of the bipartite motif in activation of downstream targets[124].

An Alphafold3-predicted structure of SLC2A4RG is consistent with the C2H2-zf not serving as the main determinant of DNA-binding, and instead with the C-clamp(s) binding the major groove precisely at the PWM-predicted binding site(s) within an extended DNA sequence (**Extended Data Fig. 2f**). The predicted structure is virtually identical to the experimentally determined structure: the SLC2A4RG C-clamp in complex with a single GCCG-containing sequence[118] possesses the same configuration a monomer, with the four cysteine-residues in the C-clamp domain coordinating a Zn2+ ion. A unique contribution of our predicted structure is that the C2H2 zinc-finger may contact the major groove, potentially stabilizing the interaction. Several of our tested C-clamp constructs exclude the C2H2-zf, however (see next section), so at least in the case of the GlutEF/ZNF395 family, the C-clamp is sufficient for high-affinity binding.

An Alphafold3-predicted structure of SLC2A4RG is consistent with the C2H2-zf not serving as the main determinant of DNA-binding, and instead with the C-clamp(s) binding the major groove precisely at the PWM-predicted binding site(s) within an extended DNA sequence (**Extended Data Fig. 2f**). The predicted structure is virtually identical to the experimentally determined structure: The SLC2A4RG C-clamp in complex with a single GCCG-containing sequence[118] possesses the same configuration a monomer, with the four cysteine-residues in the C-clamp domain coordinating a Zn2+ ion. A unique contribution of our predicted structure is that the C2H2 zinc-finger may contact the major groove, potentially stabilizing the interaction. Several of our tested C-clamp constructs exclude the C2H2-zf, however (see next section of **Supplementary Discussion**), so at least in the case of the GlutEF/ZNF395 family, the C-clamp is sufficient for high-affinity binding.

#### Unsuccessful putative TFs

Here, we discuss potential reasons for the failure of 155 Codebook proteins to yield motifs. We first examined overall trends observed amongst the unsuccessful proteins, which we discuss here, and then consider each individual protein (compiled in **Supplementary Document 1**). By extension from the success rate for positive controls (58/61, or 95%), we would expect 314 successes out of the 332 Codebook proteins but obtained only 177. Thus, we sought explanations for the remaining 155 (= 332 - 177).

We first broke the 155 proteins into three main classes (**Extended Data Fig. 3a**), which we consider in turn: the C2H2-zfs (59 failures out of 179 attempted), all other known DBDs (53 failures out of 104 attempted), and those lacking a known DBD (43 failures out of 49 attempted). Each class has specific characteristics: the C2H2-zfs are best known for binding as tandem arrays, other DBDs generally have individual known structural characteristics (and most are represented by known structures), while those lacking a known DBD are mainly isolated cases with limited support in the literature and no DNA-protein structures.

The Codebook Consortium analysis was based on our previous curation of potential TFs, which we published in the database “HumanTFs”[125], and as our purpose was to compile a list for direct experimental evaluation of their DNA binding capability, our inclusion criteria were lenient. A large fraction of the proteins were included simply based on homology (typically, because they contained a likely DNA-binding domain), even if there was no direct evidence and on the other hand, we included proteins even if their sequence-specific DNA binding activity was based only on a single study and there was no other supporting evidence. Proteins that were included based on homology had overall high success rates, whereas proteins based on scant evidence were unsuccessful in all but a few notable exceptions.

##### C2H2-zf proteins

The C2H2-zf proteins typically bind DNA as a tandem array with each C2H2-zf offset from the next by three bases. This arrangement places constraints on the “linker” in between the C2H2-zf domains. Roughly half of all linkers in human C2H2-zf arrays are seven amino acids in length. Consistent with the importance binding as a tandem array with appropriate spacing, Codebook C2H2-zf proteins that contained at least a pair of DBDs separated by 3-10 amino acids showed a high success rate (112/132, or 84.9%), unlike those with only noncanonical gaps (4/14, or 28.6%) and those with only a single C2H2-zf domain (5/21, or 23.8%). Furthermore, the experiments were unsuccessful for 9 proteins that contain only “unconventional” C2H2-zf domains, which were included based on literature evidence of containing a C2H2, but that did not pass HMMer-based prediction as used in CisBP (P-value of less than $10^{-5}$) (**Extended Data Fig. 3b**).

We then manually examined all of the C2H2-zfs in these three categories that failed. For a minority, there is credible evidence in the literature that they may bind DNA (**Supplementary Table 6, Supplementary Document 1, and Extended Data Fig. 3c**).

For 21 of them, however, there is insufficient evidence and we have classified these as unlikely to be TFs.

Considering the low success rates for non-array C2H2 proteins, we examined, case-by-case, the successful proteins in this category. There were five proteins with only a single C2H2-domain that succeeded in yielding a motif; two of them (SLC2A4RG and ZNF395) appear to use the adjacent C-clamp to bind DNA (discussed above). Another two (ZNF503 and ZNF703) are close paralogs that were successful only in ChIP-seq experiments. According to previous data, ZNF703 is a chromatin corepressor component that may not bind DNA directly, but through other proteins[126]; we note that the ChIP-seq experiments enriched posterior homeodomain reminiscent CATAAA sites and several other motifs in both experiments. The last, ZBTB5, is likely to be a genuine TF that binds DNA specifically with just a single C2H2 domain by recognizing combinations of palindromic GATC-sequences. BTB domains are known to mediate homodimerization, suggesting that the domain manages to bind DNA with a single DBD through cooperative binding. Thus, it represents an unusual case.

Analyses resulted in four successful proteins with noncanonical gaps. MYT1L represents a well-characterized exception known through analysis of its paralog MYT1. Its C2H2 domains are of a specific C2H2C (IPR036060) subtype, that are known to be able to bind DNA either individually or as cooperative combinations with a different kind of mechanism than what is used in array based C2H2 proteins – the two domains do not enter major groove as an array, but instead the long disordered region separating them allows the domains to bind opposite to each other and recognizing a palindromic consensus sequence[127,128]. The second protein, TSHZ2, contains a homeodomain in addition to its widely gapped C2H2 domains (that are folded in the expected manner based on AlphaFold3), and the ATCA-consensus that it binds is similar to the sites bound by PBX homeodomains. ZNF618 is a poorly characterized protein that contains 4 canonical (and consistently folded on AlphaFold3) C2H2 domains, and additionally an enzymatic domain belonging to the Ribonuclease H-like domain family. The Codebook evidence is from lysate-based SELEX experiments and could potentially represent indirect binding. Finally, CASZ1 represents an interesting case, as it recognizes very long target sites with gapped A or T stretch PWMs. Both CisBP and protein domain databases predict that this protein has 11 canonical C2H2 domains separated by noncanonical gaps; however, AlphaFold3 predicts that all the repetitive domains predicted as C2H2 are in fact noncanonical C2H2 protein domains with a more complex fold. When compared against canonical DNA-binding C2H2 domains, they have an additional beta-sheet (i.e., three rather than two antiparallel beta-sheets), and there is a long linker between the first two beta-sheets that folds as part of the structure. Additionally, these domains have a longer putative DNA recognition helix than what is present in typical DNA-binding C2H2 domains. Conservation analysis of CASZ1 previously detected the sequence features that fold into these domains and predicted them as a novel type of zinc finger domain called "para-ZNF"[102] but this domain type does not appear to be listed in databases. **Extended Data Fig. 3d** shows a comparison of CASZ1 zinc fingers 1 and 2 against an example of canonical C2H2 domains of (GLI2, PDB:2GLI), with the predicted C2H2 region CASZ1 DBD2 structurally aligned to DBD4 of GLI2.

Overall, 21 out of the 59 failures of C2H2-zf protein can be explained by either the number, type, or spacing of the C2H2-zfs.

**Other DBD families**

Fifty-three (53) of the unsuccessful proteins had a predicted protein domain of a type that has been shown to bind DNA in sequence specific manner in either the same protein or homologous proteins (**Extended Data Fig. 3a**). For most of these DBDs, there are DNA-protein structures and other biochemical data available for either the same protein, or others of the same DBD class. We used these data, as well as Alphafold3 predictions, and other literature, to assess whether the unsuccessful proteins have properties that are consistent with sequence-specific DNA binding.

In many cases, the unsuccessful proteins had features that would likely prevent them from engaging in sequence-specific binding. For example, the RFX domain in RFX8 is truncated and missing a DNA-contacting region. SGSM2 is annotated as containing a BED family ZF; however, Alphafold3 predicted that the putative ZBED domain is not folded into a ZBED but is a part of a larger folded protein domain. Thus, these proteins are likely unable to bind DNA in a sequence-specific manner. In addition, some of the unsuccessful Codebook proteins now have motifs in databases and other studies in the literature.

We assessed all 59 proteins on the basis of available information, and gauged whether each appeared most likely to be a sequence-specific TF (24 cases), a low-specificity DNA-binding protein and/or cofactor (20 cases), or neither (9 cases).

Altogether there are 24 potential false negatives and 29 apparent true negatives, among the “Other” DBD families. Nine that we classified as true negatives represent the THAP domain, which appears to have both low DNA affinity and low DNA sequence specificity, and seven have the AT-hook DNA binding peptide motif, which has low sequence specificity and is often found in cofactors and chromatin remodeling enzymes **(Extended Data Fig. 3c)**[129].

**“Unknown” DBD proteins**

We applied the same criteria as for the “Other” DBD class to those with no known DNA-binding domain. Most of these proteins (34/43) had been included in the Codebook study on the basis of a single previous publication, typically including SELEX and/or EMSA. For most of these (30/34), no additional evidence has since emerged in support of sequence-specific DNA binding, while some now have other known functions; e.g. recombinant KCNIP3 was reported to bind DNA by EMSA[130], but it has a well-established role in binding and controlling specific potassium channels[131].

Ten of the 43 do appear to represent likely sequence-specific DNA-binding proteins (**Supplementary Table 6 and Supplementary Document 1**). Upon re-examination of our own data, several do appear to display sequence specificity, but did not pass our stringent success criteria. These include RAG1 and SKIL. Most of the remainder are believed to require cofactors to[27,95] bind DNA. For example, PDB:4NDY shows CENPS and CENPX binding DNA in an intertwined and asymmetric manner, consistent with binding as an obligate heterodimer. The Codebook *in vitro* experiments were performed with single proteins; thus, failure of these proteins could easily represent false negatives. Most (33/43) appear more likely to represent true negatives.

**Summary**

Compiling the results above, we can explain 21+29+33 = 83 of the 155 unsuccessful Codebook TFs as likely to be true negatives (54%). For others, the expert curation identified explanations for false negatives (e.g., obligate cofactors), which will guide future efforts to obtain high-quality PWMs for these proteins.

### Transposon-derived TFs

#### Introduction

Domestication is the process by which a transposon coding sequence is exapted by the host through fusion to an existing protein (often called 'exonization' as a TE exon is captured by a host gene), or as an entirely new gene[132]. The most widespread and consistent path of TE contribution to transcription factors is in the form of DBDs derived from DNA transposons[61]. In the transposon, this DBD is needed to identify the cognate genome-integrated transposon in *trans*, typically by recognizing motifs in the transposon's Terminal Inverted Repeats (TIRs), which flank the transposase ORF. TE-derived DBDs, now within host genes, have been and continue to be identified across animals.

We identified new DNA-binding motifs for sixteen previously known domesticated Codebook proteins (and two controls), with previously putative TF function. These include CGGBP1 from *hAT-19*[62], five proteins containing BED-zf domains derived from *hAT-Ac* and *hAT-Charlie*[63], six with the related CENBP or Brinker domains from *Tigger* and *Pogo* elements[64], three with *PIF-Harbinger*-derived Myb/SANT[65] or MADF domains, and FLYWCH1 from a *Mutator*-related transposon[66]. We therefore sought to characterize the evolutionary relationships of these proteins, as well as investigate their interactions with genomic transposon fragments.

#### Pogo and Tigger-derived transcription factors

The *Tigger*/*Pogo*-derived TFs with new Codebook motifs are involved in a range of biological processes and implicated in disease[133-135]. They include TIGD3, 4, 5, and 7, JRK, and the more distantly related POGK. All of these proteins share a bipartite DBD comprised of two HTH domains, with the N-terminal domain being of the "Pipsqueak (Psq)" family[136] in Interpro[120], and the C-terminal domain simply a "CENPB-type HTH"[137]. In POGK, the Psq domain is replaced with a "Brinker"-type HTH[138], but retains the C-terminal CENPB-type HTH. Based on phylogenetic analyses, *Tigger*/*Pogo* have been subject to multiple independent domestication events across tetrapods, although the precise evolutionary histories remain uncertain[104,105]. TIGD7 and JRK do at least seem to be closely related, and may have arisen from a single domestication event in the common ancestor of mammals[64]. The PWMs obtained for *Tigger*/*Pogo*-derived TFs are often long and information-rich, which may be one strategy by which their ancestral TEs bound specifically to cognate elements integrated across a large genome (**Extended Data Fig. 9a**).

All DNA transposons, including *Tigger*/*Pogo*, have been extinct in the human lineage for over 40 million years[65]. Nonetheless, genomic binding of JRK is enriched for binding to a subset of *Tigger*/*Pogo* elements, particularly Tigger15a, and the consensus sequence for Tigger15a has a PWM-predicted binding site for JRK in the TIRs of these elements (**Extended Data Fig. 9b**), consistent with its presumed ancestral role in transposition. A TBLASTN search of the JRK protein sequence against Dfam[139] *Tigger/Pogo* consensus DNA sequences also reveals that JRK has the strongest sequence homology to

Tigger15a, suggesting that Tigger15a is the family, or is closely related to the family, from which JRK (and perhaps TIGD7) was domesticated. To our knowledge, it is only the third human protein known to still bind its ancestral transposons, including SETMAR, derived from a fusion in anthropoid primates between a *Tc1-mariner* (a distantly related superfamily to *Tigger/Pogo*) transposon and a SET domain[70], and JRK's relative CENPB, as the CENPB box has been noted to resemble the Tigger2 transposon's TIRs[140]. These cases apparently represent the simultaneous introduction of a multitude of *cis*-regulatory elements and the TF that binds them, which may be a mechanism by which many transposons have been domesticated as TFs, such as the bat KRABINER[141].

**BED-type zinc fingers are universal DNA-binding domains**

Phylogenetic analysis of the ZBED family of transcription factors suggests that they are derived from at least three different domestication events: ZBED1 from *hAT-Ac,* ZBED2, 3, 4, 6 from a different *hAT-Ac* domestication, and ZBED5, 7 (aka ZMYM6), 8 (aka FAM200C), and 9 (aka SCAND3) from a distinct family, *hAT-Charlie*[63]. Of these, only ZBED1 and ZBED6 have been closely investigated for DNA-binding activity, binding TGTCG(C/T)GA(C/T)A and GCTCG consensus sequences, respectively[106,142]. Codebook tested all remaining ZBEDs other than ZBED7 and 8, and in addition, tested the BED-zf domains of FAM200B, a poorly characterized protein closely related to ZBED9, and GTF2IRD2, which is a fusion of two GTF2I DNA-binding domains with an N-terminal *hAT-Charlie*[143].

All yielded high-quality motifs save for ZBED3, consistent with a previous study that found it to have a largely cytosolic localization, instead functioning as a Wnt signaling regulator via direct interaction with Axin[144]. CGGBP1 possesses a BED-zf-like DBD and is domesticated from *hAT-19* transposons, which form a distinct family from *Ac* and *Charlie*[62,108]. A phylogeny created using zf-BED and zf-BED-like domains from these domesticated TFs clearly recapitulates the known relationships among the three *hAT* families, as well as the evolutionary domestication histories of the ZBEDs (**Extended Data Fig. 9c**).

Like *Tigger/Pogo*-derived TFs, the *hAT*-derived TFs exhibit broad motif diversity both in sequence and length, although they often feature a CpG dinucleotide followed by 1-2 A's and ~3 C's (**Extended Data Fig. 9c**). Unlike *Tigger/Pogo*, most BED-zf motifs are short, between 6-7 nucleotides; this may reflect how *hAT* transposases recognize and bind their cognate integrated transposons, with the BED-zf binding subterminal repeats distributed at both ends of the transposon, between the TIRs and transposase ORF[145].

Interestingly, while GTF2IRD2 has GTF2I DBDs, assays using constructs containing just the GTF2I DBDs failed, while full-length protein constructs including the BED-zf were successful. The derived motif is also reminiscent of the other *hAT-Charlie*-derived DBDs (**Extended Data Fig. 9c**); these lines of evidence suggest that it is, in fact, the BED-zf domain of GTF2IRD2 that is conferring sequence-specific binding.

#### Origin of *PIF-Harbinger*-derived transcription factors

While the origins of several functionally diverse classes of DNA-binding proteins can be attributed to domestication from transposons[61], some of these DBDs may have been originally acquired from a host gene through a very ancient domain capture event. For example, the Myb domain, a tri-helical HTH, is characteristic of *PIF-Harbinger* elements[146] and of *PIF-Harbinger*-derived TFs, but is also widely distributed across eukaryotes in genes with no known homology to TEs[147]. As a result, it cannot be confidently stated yet whether all Myb DBDs are derived from *PIF-Harbinger*, or if only a subset are.

Codebook tested a number of Myb/SANT encoding proteins, including DMTF1, TERF1, TTF1, MYSM1, TERB1, MSANTD1, MSANTD4, MYPOP. However, only the latter three have homology to *PIF-Harbinger* transposons[65], although all three yielded high-confidence motifs (**Extended Data Fig. 9d**). We have also previously derived motifs for the *PIF-Harbinger*-derived MSANTD3 and NAIF1[99] (the NAIF1 motif is also consistent with a previously identified consensus binding sequence[148]). A unique characteristic of *PIF-Harbinger* transposons is that the Myb DBD and the catalytic domain are encoded on separate ORFs[149]. In all of the TFs domesticated from *PIF-Harbinger,* the protein encodes the Myb DBD with no detectable catalytic domain remnants (unlike many of the *hAT* and *Tigger/Pogo* TFs above), which is perhaps mediated by it being encoded on a separate ORF that can be exonized into another protein, or domesticated in isolation.

#### Structural prediction of FLYWCH1-DNA binding

FLYWCH1 encodes five FLYWCH-type zinc-fingers, DBDs that are encoded by the *Phantom* family of *Mutator* DNA transposons[66]. We revealed that FLYWCH1 binds to a degenerate motif, with two high information content CpG dinucleotides, and two lower information content ones (**Extended Data Fig. 9d**). FLYWCH1 had poor overlap between ChIP-seq, GHT-SELEX, and PWM hits, and all TOPs are restricted to a ~6kb region on chromosome 10, and does not have enrichment for promoters, enhancers, or the "dark genome" in our datasets. FLYWCH1 has been tentatively identified as a tumour suppressor and DNA-damage response protein in a handful of studies[150-152], and may be involved in the recruitment of H3K9me3-associated proteins[153]. We therefore sought to gain a clearer understanding of FLYWCH1 binding activity, which could drive further study into its functional role in chromatin regulation.

While an unpublished NMR solution structure of the fifth FLYWCH1 zinc-finger exists (PDB: 2RPR), it is unclear which zinc-fingers are essential for conferring DNA-binding. To generate an initial hypothesis of how FLYWCH1 interacts with its binding site, we generated an Alphafold3[154] structure of the five FLYWCH1 zinc fingers with a high-confidence TOP sequence. While the pTM and iPTM scores are both low (0.29 and 0.33 respectively), the overall structure is consistent with our observed motif, with zinc-fingers 4 and 5 inserting into major grooves containing the highest-information content CpG dinucleotides (**Extended Data Fig. 9e**), and zinc-fingers 1 and 2 interacting with much lower-information positions, which at this binding site encode slightly suboptimal GG dinucleotides. As a CpG-binding protein, binding-site affinity might be influenced by

DNA methylation, which may partially explain the low overlap between ChIP-seq and GHT-SELEX. FLYWCH1 SMiLE-seq assays were unsuccessful, so it is currently unclear how DNA methylation may influence the binding activity of the protein[155].

#### Codebook TFs are frequently enriched for binding specific families of transposons

The Codebook data also underscore that many TFs bind preferentially and intrinsically to specific repeat classes. These interactions are explored in greater detail in the accompanying manuscripts[12,13]. Binding to endogenous retroelements is known to be a common property of the KRAB-domain-containing C2H2-zf (KZNF) subfamily in vivo[156], but until now it has not been clear that the recruitment is defined almost entirely by the sequence specificity of the KZNFs alone. The combination of assays, particularly GHT-SELEX, extends earlier observations by pinpointing the exact binding sites and demonstrating that these proteins typically have high specificity for these elements, because they bind preferentially to precisely the same elements *in vitro*. Binding preferentially to retroelements is not limited to KZNFs but includes other C2H2-zf proteins and other classes of TFs. For example, binding sites for ZBED9 are enriched for binding to Alus, and some *hAT-Charlie* elements[13], the family from which it was originally domesticated[63].

### Supplementary Methods

#### Allele specific binding analysis

We reasoned that the GHT-SELEX and ChIP-seq experiments allow direct assessment of allele-specific binding (ASB) of TFs by quantifying the allelic imbalance of read counts at single-nucleotide variants (SNVs). We note that the data were not initially intended for this purpose, and caveats include relatively low read counts, linked SNVs, and the fact that HEK293 has an abnormal karyotype and was derived from a single individual. Nonetheless, SNV calling (see below) yielded 924,997 variant calls overlapping with dbSNP common SNPs (889,814 variant calls from 361 ChIP-seq experiments and 35,183 from 370 GHT-SELEX multi-cycle experiments) at 122,364 unique genomic locations (corresponding to distinct rsSNP IDs,). Of these, 10,009 SNPs corresponded to 12,060 ASBs of 152 Codebook TFs and 39 positive controls, i.e., there was a significant imbalance in the number of sequencing reads supporting the reference or the alternative SNP alleles in ChIP-seq (10,575 ASBs) or GHT-SELEX (1,485 ASBs) read alignment **Extended Data Fig. 5** and, **Supplementary Table 8**. SNP calls and ASBs are available at Zenodo (https://doi.org/10.5281/zenodo.18224872).

##### Variant calling

For variant calling directly from ChIP-seq and GHT-SELEX data, we started by mapping raw ChIP-seq and pre-trimmed GHT-SELEX reads[14] to the hg38 human genome assembly using *bwa-mem* (v.0.7.1) with default settings (**Extended Data Fig. 5a)**. Next, we used *filter_reads.py* (originally taken from *stampipes* (https://github.com/StamLab/stampipes/tree/encode-release, accessed Sept 2022) to filter out reads with >2 mismatches and mapping quality <10. Then, we followed the workflow of [27] for SNV calling and read counting (https://github.com/autosome-ru/MixALime/tree/main/natcomm_supp_scripts, accessed Dec 2025):

(1) *samtools reheader* (v.1.16.1) was used to set the identical sample SM field in all alignment files;

(2) SNP calling was performed using *bcftools mpileup* (v.1.10.2) with *--redo-BAQ --adjust-MQ 50 --gap-frac 0.05 --max-depth 10000* and *bcftools call* with *--keep-alts --multiallelic-caller*;

(3) the resulting SNPs were split into biallelic records using *bcftools norm* with *--check-ref x -m -* followed by filtering with *bcftools filter -i "QUAL>=10 & FORMAT/GQ>=20 & FORMAT/DP>=10" --SnpGap 3 --IndelGap 10* and *bcftools view -m2 -M2 -v snps* leaving only biallelic SNPs covered by 10 or more reads;

(4) SNPs were annotated using *bcftools annotate* with *--columns ID,CAF,TOPMED* and dbSNP (v.151)[157];

(5) heterozygous variants located on the reference chromosomes with GQ ≥20, depth ≥10, and allelic counts ≥5 on each allele were filtered with *awk* (v.5.0.1);

(6) *WASP* (v.0.3.4)[96] was used with *bwa-mem* and *filter_reads.py* to account for reference mapping bias;

(7) *count_tags_pileup_new.py* (edited version of the original script *count_tags_pileup.py* from https://github.com/vierstralab/nf-allelic-mapping/tree/main/bin adapted by removing the settings related to DNase-seq specifics) was used to obtain allelic read counts with *pysam* (v.0.20.0);

(8) *recode_vcf.py* was used to convert the resulting BED files to VCF.

Of note, triallelic SNVs were split into two biallelic records.

**ASB calling and annotation**

ASB calling was performed independently for GHT-SELEX and ChIP-seq data. To account for aneuploidy and copy-number variation, the profiles of relative background allelic dosage were reconstructed with BABACHI (v.2.0.26) using default settings[97] (Abstract O3). The allelic imbalance was estimated with MIXALIME (v.2.14.17)[27], starting with mixalime create. Next, we fitted a marginalized compound negative binomial model (MCNB) using mixalime fit specifying MCNB and setting --*window-size* to 1000 and 10000 for GHT-SELEX and ChIP-Seq, respectively, taking into account lower coverage and fewer SNPs called from GHT-SELEX. Finally, we used mixalime *test* followed by TF-wise mixalime combine to obtain the TF-specific ASB calls. This resulted in 12,060 identified ASBs at 5% FDR.

Technically, the genotype quality (GQ) of SNPs and ASBs was much higher than the default threshold, and the majority of ASB SNPs were supported by two or more datasets (**Extended Data Fig. 5d,e**).

We then identified 3,564 ASBs that overlap a PWM hit (P-value < 0.001) for the associated TF. Of note, ASBs that do not overlap a PWM hit may be marker variants acting indirectly by being linked to “causative” SNVs. For ASBs with PWM hits, we calculated the PWM scores for both alleles and estimated the P-values of those scores against a uniform background distribution using PERFECTOS-APE[98]. The fold-change between Alt (alternative) and Ref (reference) allele P-values, $\log_2$(Alt/Ref), reflected the PWM-predicted allelic preferences, with positive (negative) values reflecting the preference for the Alt (Ref) allele, respectively. The ASBs with the absolute $\log_2$(fold-change)>1 were labelled as “motif-concordant” or “motif-discordant”, depending on whether the allelic preference exhibited by the greater ChIP-Seq or GHT-SELEX read coverage (Ref>Alt or Alt>Ref) was consistent with the difference in the respective PWM scores (**Extended Data Fig. 5c**).

To globally support the relevance of the Codebook ASB calls, we (1) obtained a joint set of 22,064 ASBs by running *MIXALIME* (v2.28.0) *multiple_combine* for joint aggregation of the allelic imbalance P-values over all processed datasets (**Supplementary Table 9**) and (2) for the resulting joint set of ASB-SNPs, estimated the significance of the overlap with GTEx v.8[30], ADASTRA v.6.1[28], and EBI GWAS Catalog v1 (e115_r2025-12-03_full)[29] using two-sided Fisher's exact test.

#### Motif Activity Response Analysis

**Identifying and scoring promoter sequences.** Motif activity response analysis requires promoter activity (gene expression) data across samples and motif scores across promoters. The former was downloaded from the FANTOM5 web resource (https://fantom.gsc.riken.jp/5/), *hg38_fair+new_CAGE_peaks_phase1and2_tpm_ann.osc.txt*, $\log_2$-transformed with a pseudocount of 0.05, and filtered, leaving only promoters of genes encoded in the nuclear genome as well as removing time courses, perturbations, and human total RNA samples. This resulted in 209374 individual promoters and 1020 samples (including replicates), belonging to 583 unique samples (142 tissues, 187 primary cells, and 254 cell lines), see **Supplementary Table 12**. Motif scanning was performed on regions from FANTOM5 *hg38_fair+new_CAGE_peaks_phase1and2.bed,* taking 250bp upstream and 10bp downstream from the representative TSS position as indicated in FANTOM5 data. Next, we used SPRY-SARUS (https://github.com/autosome-ru/sarus) to compute the sum-occupancy scores[90] for each representative motif of the motif clusters(obtained as described in Methods). The final analysis was performed with 632 motif clusters (representative motifs) corresponding to TFs that were jointly expressed >0 in at least one of the FANTOM5 samples.

**MARA model formulation and implementation.** For motif activity response analysis (MARA), we employed MARADONER (MARA-done-right) v0.13, a command-line tool written in Python and available in the PyPi repository. The analysis was performed using maradoner create, maradoner fit, and maradoner export with default parameters (https://github.com/autosome-ru/MARADONER).

Conceptually, MARADONER extends the original logic of MARA[45] and isMARA[46]. The basic assumption is that the promoter activity in each sample is a linear function of sample-specific motif activities, where each motif represents a set of TFs with shared binding specificity. We employ a matrix-variate linear mixed model:

$$Y = \mu_p {1_s}^T + 1_p \mu_s + B\,U + E, E \sim MN(0, I_p, D), U \sim MN(\mu_m {1_s}^T, \Sigma, G),$$

where $Y$ is a matrix of promoter activity in log-scale of shape $p \times s$ (where $p$ is the number of promoters and $s$ is the total number of samples), $\mu_p$ and $\mu_s$ are promoter-wise and sample-wise means, $1_n$ is a vector of ones of length $n$, $B$ is a matrix of promoter-level motif scores of shape $p \times m$, $U$ is a random matrix of motif activities of shape $m \times s$, $E$ is a random error/noise matrix of shape $p \times s$, $MN$ is a matrix-variate normal distribution, $I_p$ is an identity matrix of shape $p \times p$, $D$ is a diagonal matrix of noise variances of shape $s \times s$, $\Sigma$ is a $m \times m$ matrix of motif variances, $G$ is a diagonal $s \times s$ sample-wise scaling matrix, $\mu_m$ is a motif-wise mean value of motif activities. In contrast to the classical MARA, modeling $\mu_m$ allows explicitly distinguishing activators from repressors. The number of unique parameters in each of the diagonal matrices $D$ and $G$ is equal to $g$ (the number of groups, in our case, the number of unique samples excluding replicates), which is less than $s$ (the total number of samples), allowing for an increase in certainty in the parameter estimates of $D, G$.

MARADONER performs estimation via a four-stage restricted maximum likelihood (REML) procedure. Firstly, we isolate parameters in $E$ by finding a transformation that is orthogonal to $1_p, 1_s, B$ matrices, allowing us to focus on estimating $D$ solely. Secondly, we find a transformation that is orthogonal only to $1_p, 1_s$ vectors. This makes estimating parameters in $\Sigma, G$ possible given known parameters in $D$. Thirdly, we find a transformation that is orthogonal to $B$ only, and we estimate the total mean effect of the $\mu_p 1_s{}^T + 1_p \mu_s$ term. Finally, given the knowledge of all other parameters, we estimate the mean motif activity vector $\mu_m$ . Then, if we disentangle $U$ from $B\mu_m 1_s{}^T$, the deviation from motif mean quantity $\widehat{U} = U - B\mu_m 1_s{}^T$ can be interpreted as a sample-specific variation in motif activity. $\widehat{U}$ is then obtained as a maximum a posteriori (MAP) estimate. Alongside "raw" MAP estimates of $\widehat{U}$, for downstream analysis MARADONER also reports standardized motif activities, obtained by dividing each motif activity by the square root of its posterior variance.

The availability of the maximum likelihood estimates (MLEs) of motif-specific variances allows for an ANOVA-like test using the asymptotic properties of MLE for each motif. To this end, MARADONER performs the Wald test by extracting by square root of the diagonal entries from the asymptotic covariance matrix of parameter estimates that correspond to elements from $\Sigma$ (the standard errors).

MARADONER assumes that the gene expression (or promoter activity, in the case of FANTOM5 CAGE data) is provided in the log-scale. As for the motif-scores matrix $B$, by default, each column is normalized by taking the negative logarithm of its empirical survival function.
